# FICD activity and AMPylation remodelling modulate human neurogenesis

**DOI:** 10.1101/787929

**Authors:** Pavel Kielkowski, Isabel Y. Buchsbaum, Volker C. Kirsch, Nina C. Bach, Micha Drukker, Silvia Cappello, Stephan A. Sieber

**Affiliations:** Department of Chemistry, Technical University of Munich, Garching, Germany.; Max Planck Institute of Psychiatry, Munich, Germany.; Graduate School of Systemic Neurosciences, Ludwig-Maximilians-University Munich, Planegg, Germany; Helmholtz Center, Munich, Germany

## Abstract

Posttranslational modification (PTM) of proteins represents an important cellular mechanism for controlling diverse functions such as signalling, localisation or protein-protein interactions^1^. AMPylation (also termed adenylylation) has recently been discovered as a prevalent PTM for regulating protein activity^2^. In human cells AMPylation has been exclusively studied with the FICD protein^3–6^. Here we investigate the role of AMPylation in human neurogenesis by introducing a cell-permeable propargyl adenosine pronucleotide probe to infiltrate cellular AMPylation pathways and report distinct modifications in intact cancer cell lines, human-derived stem cells, neural progenitor cells (NPCs), neurons and cerebral organoids (COs) via LC-MS/MS as well as imaging methods. A total of 162 AMP modified proteins were identified. FICD-dependent AMPylation remodelling accelerates differentiation of neural progenitor cells into mature neurons in COs, demonstrating a so far unknown trigger of human neurogenesis.

---

Introduction of protein PTMs is a tightly controlled and almost ubiquitous process that often modulates critical protein function. PTMs such as tyrosination, acetylation and neddylation are known to play a crucial role in the development of the nervous system and in particular of neurons by broadening the diversity of the tubulin and microtubule proteoforms^7, 8^.

AMPylation was first discovered in *Escherichia coli* as regulator of glutamine synthetase activity^9^. Later, it was found that bacterial effectors from *Vibrio parahaemolyticus* and *Histophilus somni* AMPylate Rho guanosine triphosphatases (GTPases) in human host cells^10, 11^. These bacterial effectors contain highly conserved Fic (filamentation induced by cAMP) domains, which catalyse the transfer of AMP onto a serine, threonine or tyrosine residue of a substrate protein (Fig. 1a). Approximately 3,000 members of this family are known to contain the conserved HXFX(D/E)GNGRXXR sequence motif throughout all domains of life^12^. Despite their abundance in bacteria, only one human protein AMPylator containing the signature Fic domain, termed FICD (also known as Huntingtin yeast partner E, HYPE), has been discovered^12^. Structural and biochemical studies with FICD have revealed that its activity is tightly regulated and controlled by an autoinhibitory loop. Mutation of E234 to glycine overrides autoinhibition and results in a constitutively activated enzyme^12^; the mutant form H363 to alanine is catalytically inactive^4^. One known substrate of FICD is HSPA5, which is a chaperone located in the endoplasmic reticulum (ER) and master regulator of the unfolded protein response (UPR)^3–6^. Recent data show that FICD regulates the ATPase activity of HSPA5 and its interactions with unfolded proteins, but the exact function is not yet clear^13^. However, it was found that the HSPA5 AMPylation associates with changes in neuronal fitness in *drosophila*^3, 14–16^.

**Figure 1.**
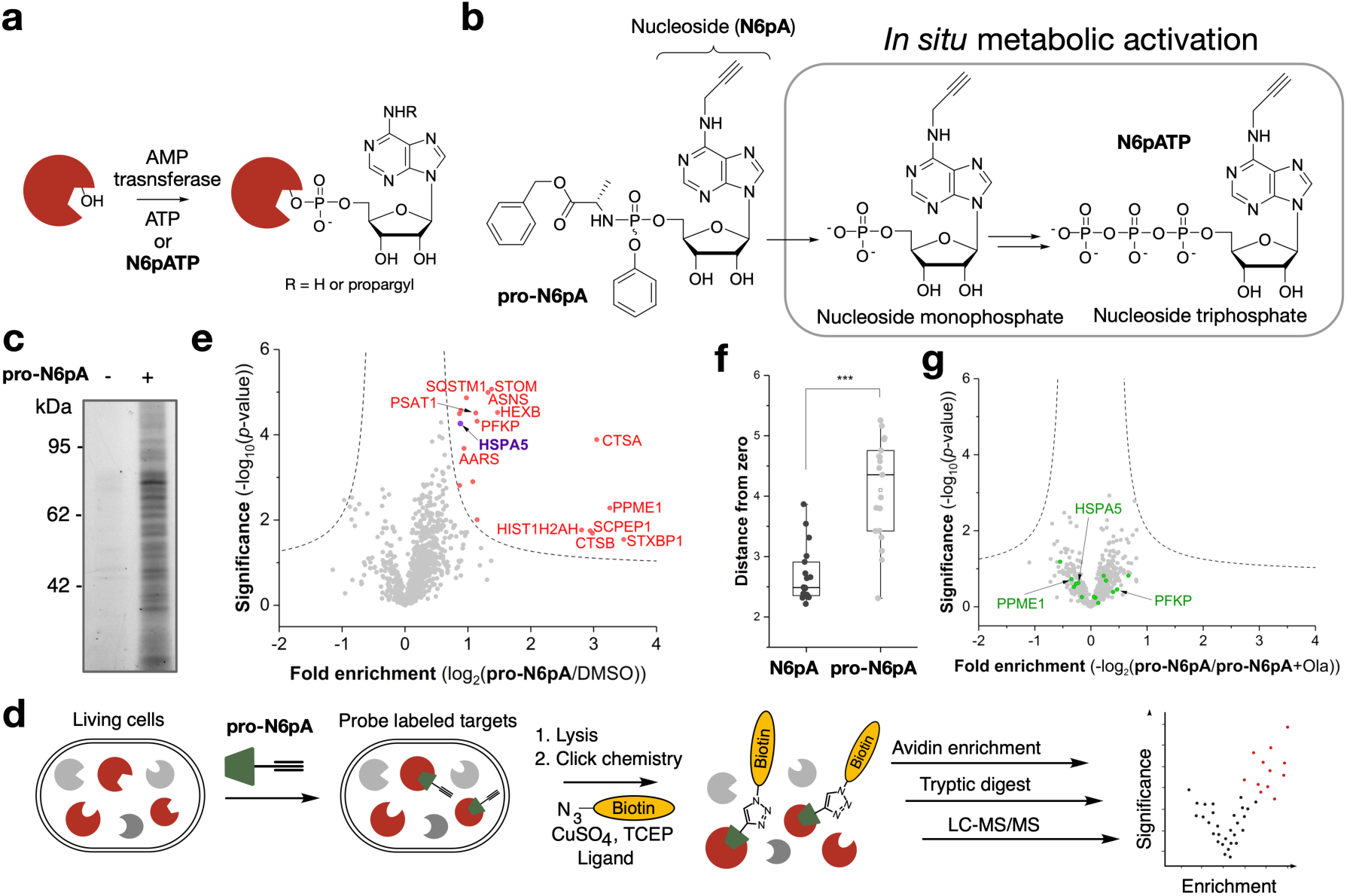
Pronucleotide probe reveals AMPylation of diverse proteins in HeLa cells. **a**, AMPylation on Ser, Thr or Tyr. **b**, Scheme of the pronucleotide probe **pro-N6pA** and parent adenosine derivative (**N6pA**) and its *in situ* activation. **c**, SDS-PAGE with in-gel fluorescence scanning showing *in situ* HeLa cell labelling by **pro-N6pA** compared to control (DMSO). **d**, Schematic representation of the chemical-proteomic approach used for *in situ* identification of AMPylated proteins. **e**, Volcano plot of fold-enrichment in HeLa cells by **pro-N6pA** labelling compared to DMSO versus significance upon two-sample *t*-test (FDR 0.05, s0 0.3; *n* = 12). **f**, Box plot representing comparison of labelling efficiency of pronucleotide **pro-N6pA** (light grey, *n* = 12) and parent nucleoside **N6pA** (black, *n* = 11); Squares represent the mean of the distances to zero for enriched proteins, lines represent the median of the distances to zero and whiskers stand for min and max values. Statistical significance was calculated with two-tailed Student’s *t*-test; \*\*\**P* < 0.001. **g**, Volcano plot of fold-enrichment by **pro-N6pA** labelling compared to probe and Ola treated HeLa cells versus significance upon two-sample *t*-test (FDR 0.05, s0 0.3; *n* = 8). Green circles represent proteins identified as AMPylated in HeLa cells.

Just recently, the highly conserved pseudokinase selenoprotein-O (SelO) was found to possess AMP transferase activity in eukaryotic cells^17^. Pseudokinases account for about 10% of the human kinome, but lack the characteristic active site residues and hence their function is largely unknown. However, their putative AMPylation activity is pointing to a possibly larger number of AMPylated proteins in human cells.

Lately, *N*^6^-propargyl adenosine-5‘-*O*-triphosphate (N^6^pATP)-derived probes have been applied to profile substrates of AMPylation in cell lysate^18–20^. Nevertheless, the most pressing, unaddressed challenge in discovering the function of AMPylation is the global analysis of AMPylated substrates under physiological conditions inside living cells. Particularly, ATP-derived probes suffer from restricted uptake of the charged nucleotides as well as competition with high endogenous ATP levels. Thus, new concepts are urgently needed to unravel the function of AMPylation in eukaryotic cells. Here we present a chemical-proteomic approach for identification of protein AMPylation in living cells using a pronucleotide probe and uncover FICD-dependent acceleration of neuronal differentiation in cerebral organoids (COs).

## Results

### 6-Propargyl adenosine pronucleotide reports on protein AMPylation

To approach the challenge of identifying AMPylated proteins *in situ*, we selected a phosphoramidate pronucleotide strategy (Fig. 1a, b)^21^. This delivery method not only enhances the probes’ cell membrane permeability but also bypasses the first phosphorylation of the nucleoside analogue by kinases. Based on these considerations, we designed and synthesized a *N*^6^-propargyl adenosine phosphoramidate pronucleotide (**pro-N6pA,** Fig. S1). We initiated our investigations with metabolomics experiments to determine **pro-N6pA** *in situ* metabolic activation to the corresponding **N6pATP**. A maximum concentration is reached 8 hours after **pro-N6pA** addition and it is maintained for at least 24 hours (Fig. S2). For the subsequent analysis of AMPylated proteins, we treated living (intact) HeLa cells with **pro-N6pA** (100 µM in DMSO) or dimethylsulfoxide (DMSO). Subsequent click-chemistry to a rhodamine-biotin-azide tag, enrichment on avidin beads and SDS-PAGE analysis via in-gel fluorescence detection revealed several distinct protein bands in the soluble fraction (Fig. 1c). Next, we performed quantitative proteome profiling in HeLa cells^22^. Enriched proteins were trypsin digested and resulting peptides were either isotopically marked by dimethyl labelling (DiMe) prior to LC-MS/MS measurement or analysed directly using label-free quantification (LFQ) (Fig. 1d)^23, 24^. Comparing **pro-N6pA** labelling (Fig. 1e) with parent *N*^6^-propargyl adenosine (**N6pA**, Fig. 1f, Fig. S3) yielded a larger number of significantly enriched proteins with the pronucleotide. Using **pro-N6pA**, a diverse group of 19 proteins was identified in HeLa cells, including the known FICD substrate HSPA5 (Fig. 1e, Table S1). Immunoprecipitation of the two selected proteins PFKP and PPME1 from the probe treated HeLa cells followed by click reaction with rhodamine-azide tag confirmed incorporation of the probe into these proteins (Fig. S3).

Although the *N*^6^-propargyl ATP analogue could, in principle, serve as precursor for ADP-ribosylation^25^, our controls indicate that ADP-ribosylation is not a major route. ADP-ribosylation is usually induced by stress conditions e.g. by addition of hydrogen peroxide to the cells’ media^26^. First, HeLa cells were pre-treated with poly(ADP-ribose)polymerases (PARP) inhibitors 4-aminobenzamide (4-ABA) or olaparib (Ola) prior to **pro-N6pA** labelling. For both PARP inhibitors, no influence on labelling intensity was observed based on in-gel fluorescence analysis (Fig. S3). In addition, MS-based chemical-proteomic experiments with Ola and **pro-N6pA** treated cells confirmed no changes in AMPylation (Fig. 1g and Table S2). Second, only two of our identified AMPylated proteins in HeLa cells (HIST1H2AH, RPS10) matched known ADP-ribosylated proteins (Fig. S3)^27^. Known ADP-ribosylated proteins were excluded as potential hits in the following experiments (Table S3).

### AMPylation of cathepsin B inhibits its peptidase activity *in vitro*

In order to validate our approach in more detail, we have employed an azide-TEV-cleavable-biotin linker during the pull-down procedure to identify the corresponding AMPylation sites of modification via MS/MS (Fig. 2a, b)^28^. We were able to directly analyse AMPylated peptides on three different cysteine cathepsin proteases CTSB (S104 and S107), CTSC (S254) and CTSL (S137) in HeLa cells (Fig. S4 and S5). All of the AMPylation sites were located on serine residues within the conserved sequence surrounding the catalytically active cysteine (Fig. 2c), suggesting that the bulky AMP modification might obstruct the binding of the peptide substrates and thus inhibit protease activity^29^. To determine whether FICD is the AMPylator of these cathepsins, we used an *in vitro* peptidase activity assay and found that cathepsin B is indeed inhibited upon FICD (wild-type (wt) or E234A mutant) treatment and did not observe any inhibition without the addition of ATP (Fig. 2d, Fig. S4). The direct measurement of AMPylation sites in CTSB (S104,107) *in vitro* was restricted by preparation of the recombinant double mutant CTSB which did not fold into the active protein, likely due to the mutation of the crucial amino acid residues within the conserved active site. Moreover, the TEV-linker based enrichment of modified peptides was performed with other cell types used in this study and three additional sites on MYH9, RAI14 and AASS (on Thr, Ser and Tyr residues respectively) were detected (Table S4). Of note, the MS-based identification of AMPylated sites in living cells is limited by the endogenous degree of modification. We thus assume that site identifications of proteins with lower AMP abundance are challenged by the detection limit. Here, previous trials in cell lysates using an active recombinant FICD E234G mutant yielded a complementary set of proteins likely due to an increased degree of modification (Fig. S6 and Table S5)^20^.

**Figure 2.**
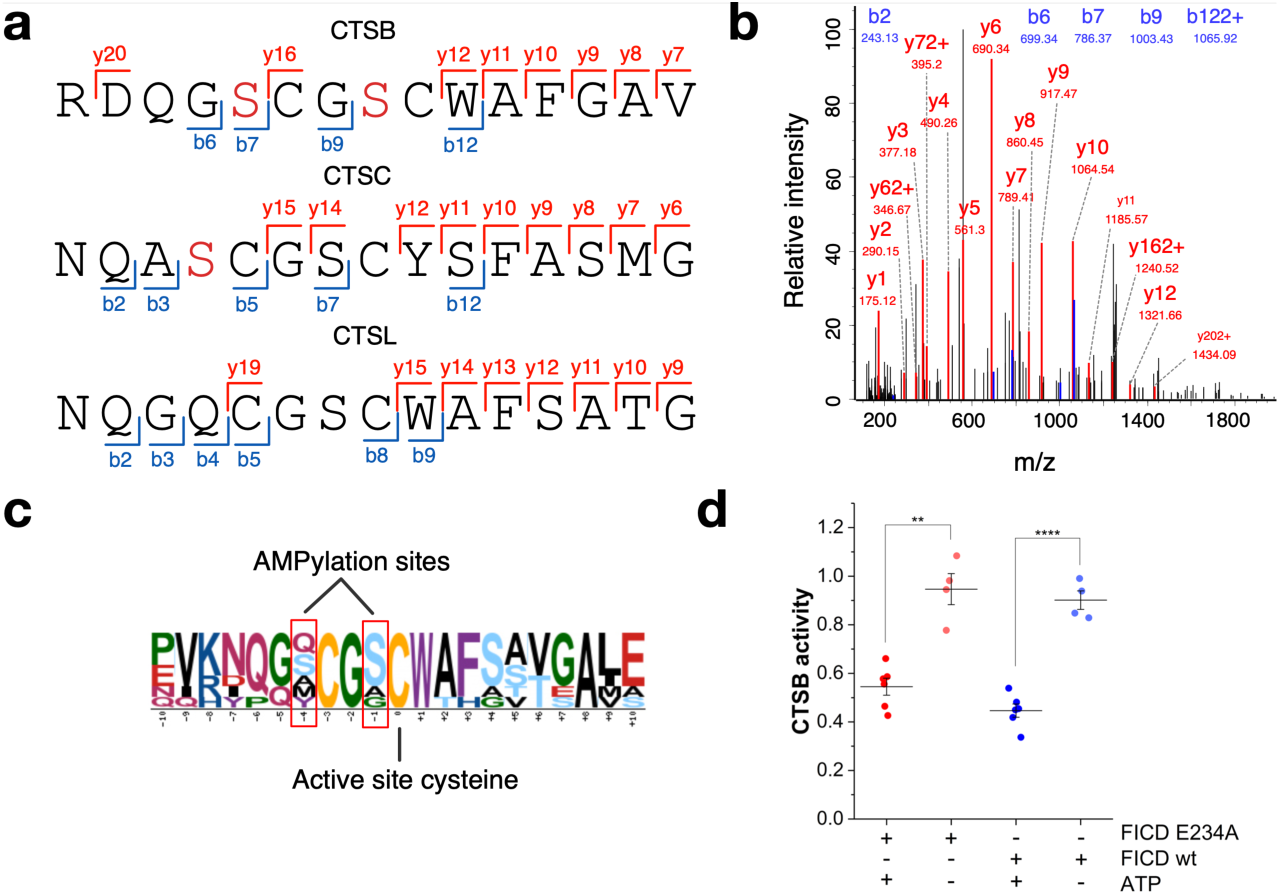
CTSB peptidase activity is inhibited by FICD catalysed AMPylation. **a**, Identified AMPylation sites on serine residues (red) of cathepsins B, C and L using **pro-N6pA** in HeLa cells. **b**, Exemplary MS/MS spectrum (MaxQuant) for the CTSB AMPylation site identification on S107 (see Fig. S4 and S5). **c**, Amino acid motif surrounding the active site cysteine of cysteine cathepsins. **d**, *In vitro* peptidase assay of CTSB activity after incubation with wt FICD or E234A mutant and with or without ATP for 6 h. Normalized to CTSB activity without FICD protein. Lines represent the mean and whiskers stand for 25^th^ and 75^th^ percentile. Two-tailed Student’s *t*-test; \*\**P* < 0.01, **** *P* < 0.0001.

### Chemical-proteomics profiling shows cell type dependent AMPylation pattern and in part independency of FICD

We performed proteome profiling in three different cancer cell lines, HeLa, A549 and SH-SY5Y, which revealed a total of 58 significantly enriched proteins, of which 38 were contributed solely from the latter neuroblastoma cells (Fig. 3a, Fig. S7). Overall, AMPylated proteins identified here are involved in diverse metabolic pathways including a widely conserved key regulator of glycolysis ATP-dependent 6-phosphofructokinase (PFKP)^30^, proteolysis (CTSA, CTSB)^31^, regulation of PTMs (PPME1)^32^ and UPR (HSPA5 and SQSTM1)^33^. Intriguingly, only PFKP was found to be AMPylated in all three cell lines, which otherwise exhibited unique AMPylation patterns.

**Figure 3.**
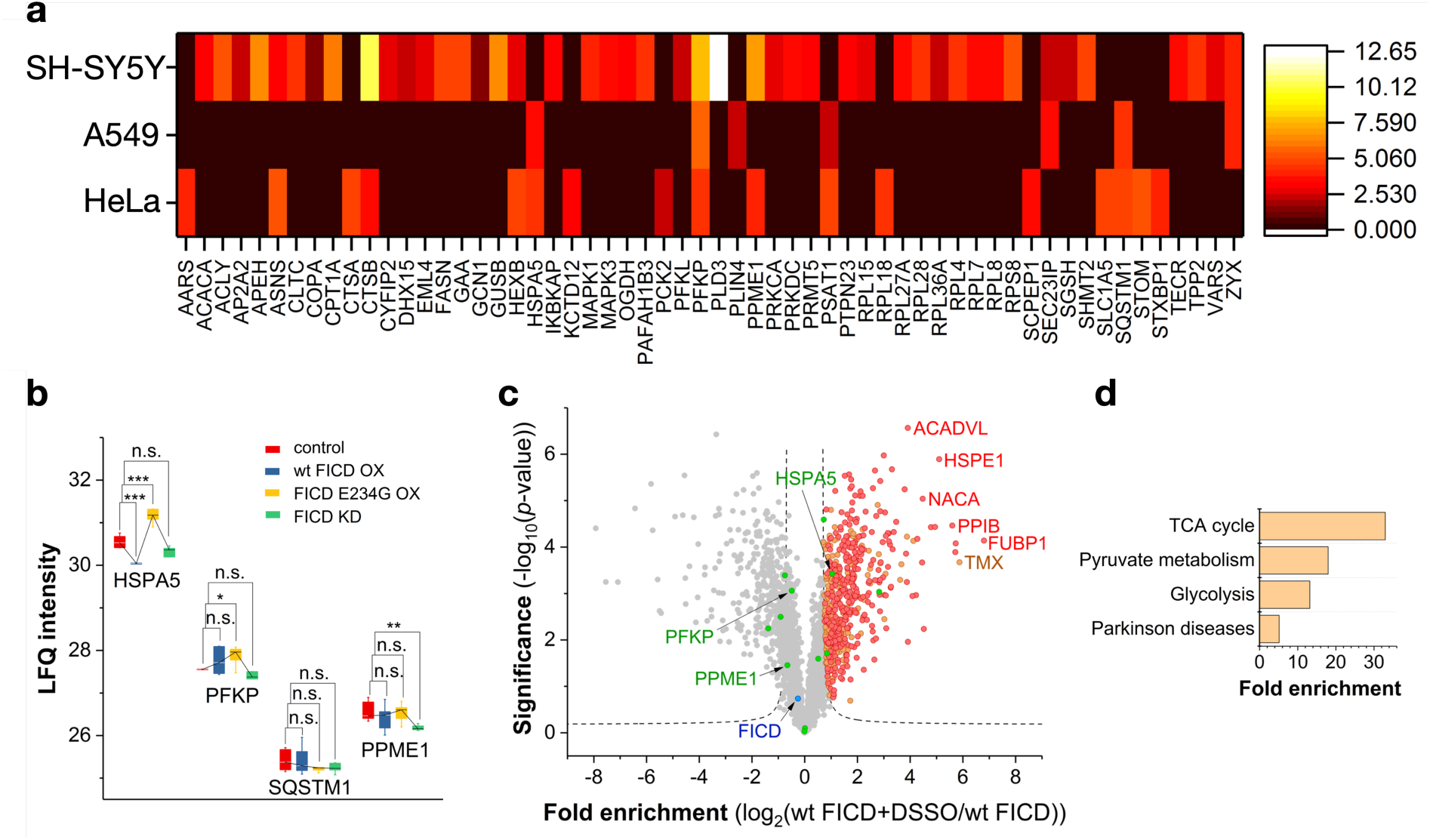
AMPylation in cancer cell lines, FICD-dependent AMPylation and FICD-interacting proteins. **a**, Heatmap representation of enriched proteins identified in cancer cell lines. Colour represents distance to zero of enriched proteins from respective volcano plot (FDR 0.05, s0 0.3; *n* = 8 or 9). **b**, Changes in AMPylation on selected proteins (those identified in HeLa cells from Fig. 1e) upon FICD overexpression (OX) or siRNA-mediated knock-down in (KD) HeLa cells. Statistical significance was tested using two-tailed Student’s t-test; \**P* < 0.05 ** *P* < 0.01, *** *P* < 0.001. **c**, Volcano plot representing FICD interacting proteins identified in the pull-down experiment of his tag labelled FICD and DSSO cross-linking reagent (FDR 0.01; s0 1.5; *n* = 3). Green circles represent proteins identified as AMPylated in HeLa cells. Red circles represent hits overlapping with parallel experiment with FICD E234G mutant. Orange circles are hits enriched only with wt FICD. **d**, Panther Pathways enriched from FICD-interacting proteins, cut-off FDR 0.05.

To directly dissect the descent of AMPylated proteins from FICD, we compared the AMPylation levels of proteins in probe treated HeLa cells comprising FICD knockdown, wt FICD overexpression (OX) and activated FICD E234G OX (Fig. 3b, Fig. S8 and Table S6). Interestingly, HSPA5 is a clear FICD-dependent responder where AMPylation is significantly upregulated in FICD E234G OX and downregulated in wt FICD OX, which is also known to perform de-AMPylation^6^. Remarkably, while all previous studies have been carried out *in vitro*^18–20^, we here independently confirm this data by the first *in situ* experiments. A direct *in situ* interaction is further corroborated by MS-based pulldown experiments of wt FICD and FICD E234G in the presence of a chemical crosslinker, which revealed HSPA5 together with other sets of proteins as interacting partners, while proteins like SQSTM1, PFKP and PPME1 were not enriched and thus considered as not interacting with FICD (Fig. 3c)^34^. Of note, FICD E234G revealed a more pronounced interaction with HSPA5, confirming our OX studies (Fig. S9 and Table S7). GO term analysis of the overlapping interacting partners indicated a link to basal metabolism (Fig. 3d). With HSPA5 as a validated candidate, we moved on and analysed other AMPylated proteins. The set of hits including PPME1, PFKP and SQSTM1 exhibited no significant changes in AMPylation levels upon FICD KD and OX, suggesting an FICD-independent mode of AMPylation (Fig. 3b) for which the origin of the AMP transfer could not be fully deduced. Given the recent discovery of an additional AMPylating enzyme also in eukaryotic cells, it is likely that the other proteins detected here descent form (a) yet undiscovered AMPylator(s)^17^.

Next, proteome profiling under endoplasmic reticulum (ER) stress conditions was performed to determine whether modifications are altered as previously reported for thapsigargin (Tg)-treated cells^6^. Only a slight increase by 2.5-fold in AMPylated HSPA5 was observed in HeLa and A549 cells. Quantification of HSPA5 by western blot shows an increase in HSPA5 expression by more than 11-fold after Tg treatment. Thus, normalization of total AMPylation to the expression of HSPA5 results in an overall reduction of its AMPylation, which is in line with previously published results (Fig. S7)^6^. Despite the moderate impact of ER stress on AMPylation in these cells, we found 145 dysregulated proteins in SH-SY5Y neuroblastoma cells (Fig. S7). The high amount of AMPylated proteins in SH-SY5Y under baseline and ER stress conditions indicates that AMPylation may have a specific role in the nervous system.

### Chemical-proteomics profiling reveals remodelling of AMPylation during neuronal differentiation

The large number of hits in neuroblastoma cells (Fig. 3a) indicates a specific importance of AMPylation in neural cells. To study AMPylation in a model system of developing neurons, neural progenitor cells (NPCs) and neurons were generated from human induced pluripotent stem cells (iPSCs, Fig. S10)^35^. Successively, iPSCs, NPCs and neurons were each treated with **pro-N6pA** and the enriched proteins were analysed via LFQ LC-MS/MS (Fig. 4a, Fig. S11 and S12 and Table S1 and S8). While PFKP was AMPylated in both the proliferating cell lines and neurons, the proteins CTSB, PSAT1 and PPME1 were only AMPylated in proliferating cells. Importantly, neurons exhibited the largest number of significantly and differentially AMPylated proteins (55 total), including transport proteins (KIF21A, KIF5C, MYH3, MYH7, MYH8) and cytoskeletal proteins (TUBB, TUBB2B, TUBB3B, TUBB4B, MAP2) (Fig. 4a). This is of particular interest as the cytoskeletal remodelling, which is required for neuronal polarisation, migration, and proper axon guidance, is a highly dynamic processes precisely regulated by several PTMs on tubulin and microtubules - and AMPylation may indeed be an additional one (Fig. S13)^7, 8, 36, 37^. AMPylation remodelling could be involved in the process of cell type specification and differentiation from iPSCs through NPCs to neurons, with cellular proteins undergoing substantial de- and re-AMPylation following an hourglass-like model (Fig. 4b, Fig. S11)^2, 6, 20^. Further, parallel chemical-proteomics studies of AMPylation under ER stress induced by Tg in iPSCs, NPCs and neurons showed distinct responses ranging from a strong change of AMPylation of several proteins in iPSCs over mild alterations in NPCs to an obvious upregulation on two proteins (HSPA5 and SQSTM1) in neurons (Fig. S11).

**Figure 4.**
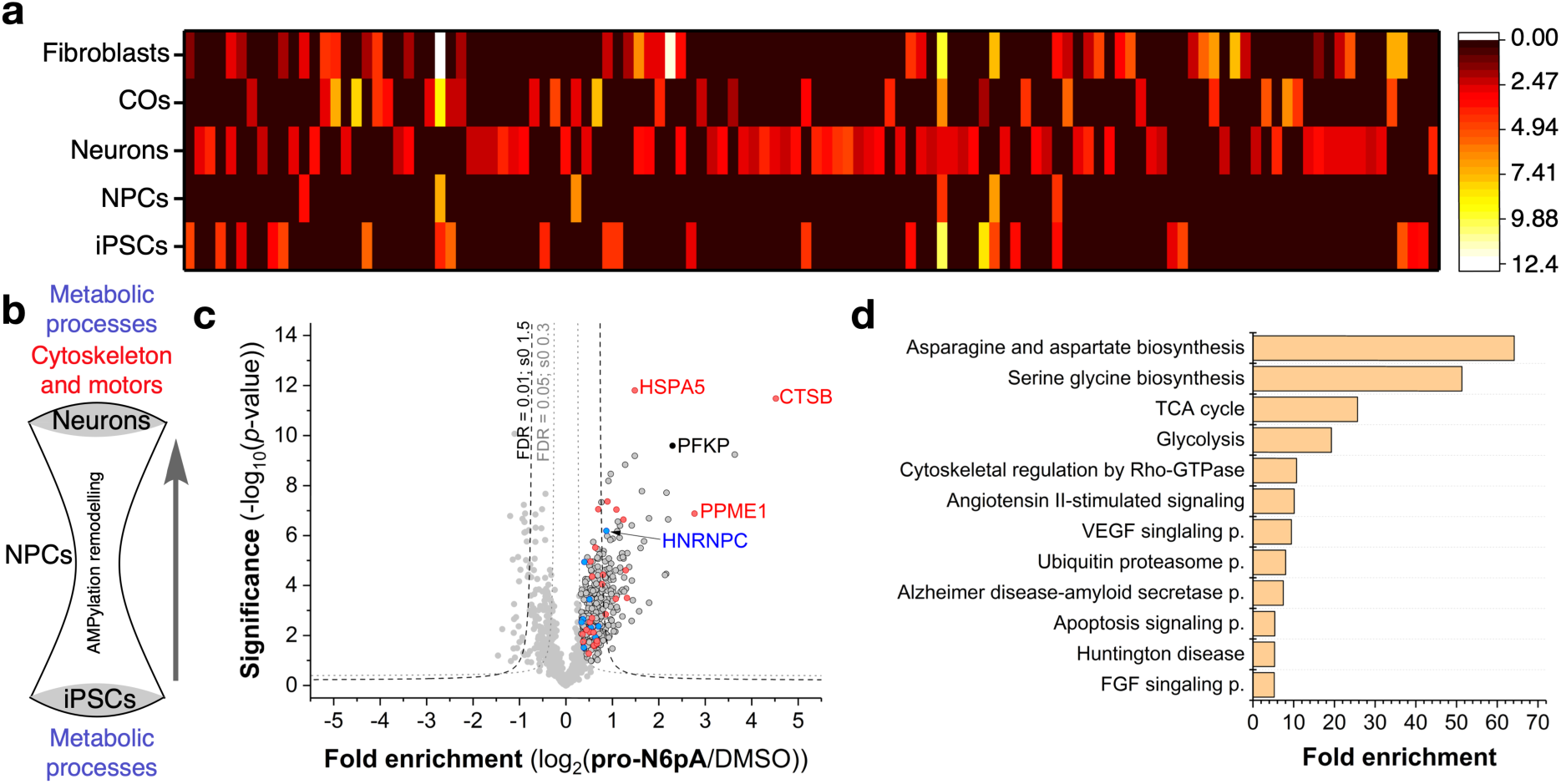
AMPylation remodelling is specific for the development of the neuronal cells. **a**, Heatmap representation of enriched proteins identified in different cell types and COs. Color represents distance to zero of enriched proteins from respective volcano plot (FDR 0.05, s0 0.3; *n* = 8 or 9). **b**, The hourglass model of AMPylation remodelling hypothesises a complete de- and re-AMPylation in the process of neuronal differentiation: from a high number of AMPylated proteins in proliferative iPSCs, most of which are involved in metabolic processes and possess catalytic activity, differentiating cells pass through a state of very sparse AMPylation as NPCs, with final neuronal differentiation resulting in neuronal identity with a high number of newly and uniquely AMPylated proteins which are enriched in metabolic functions on the one hand and in cytoskeletal and molecular motor functions on the other hand. **c**, Volcano plot of fold-enrichment by **pro-N6pA** labelling compared to DMSO versus significance upon two-sample t-test (FDR 0.05, s0 0.3; *n* = 9) in fibroblasts. Red circles represent proteins identified AMPylated in proliferating cell types while blue circles stand for overlap with hits in neurons. **d**, Panther Pathways enriched within the identified AMPylated proteins in all tested cell types, cut-off FDR 0.05.

To specify if the observed AMPylation in neurons is common for differentiated postmitotic cells we performed chemical profiling in fibroblasts (Fig. 4c, Fig. S14). Analysis of the enriched proteins revealed similarities with tested cancer cells and proliferating cell types. Most significantly enriched proteins included HSPA5, CTSB, PFKP and PPME1, all common to the proliferating cells. This highlights a distinct AMPylation remodelling in neurons.

GO term analysis of AMPylated proteins found in all screened cell types using the Panther Pathway tool displayed enrichment of basal metabolism such as TCA cycle and glycolysis as well as neuronal specific pathways including cytoskeletal regulation by Rho-GTPase and FGF signalling. Interestingly, pathways marking neurodegenerative diseases, e.g. Alzheimer, the disease-amyloid secretase pathway and Huntingtin disease were identified as well (Fig. 4d).

### FICD and most AMPylated proteins have different cellular localisations

The chemical-proteomic results were corroborated by fluorescence imaging of probe-treated HeLa, iPSCs, NPCs and neurons (Fig. 5, Fig. S15 and Table S9). In order to rule out signals derived from *N*^6^-propargyladenosine nucleotide incorporation into RNAs (e.g. in polyA tails of mRNA)^38^, we performed a control experiment in which the RNA of the fixed cells was digested with different concentrations of RNase prior to click-chemistry and as positive control of the RNase digest 5-ethynyl uridine (5-EU) stained RNAs were degraded in parallel to **pro-N6pA** labelling. Indeed, we observed only a slight decrease in overall cell staining by **pro-N6pA** rather than disappearance of the bright AMPylation spots, while we observed a strong decrease in 5-EU labelling (Fig. S16).

**Figure 5.**
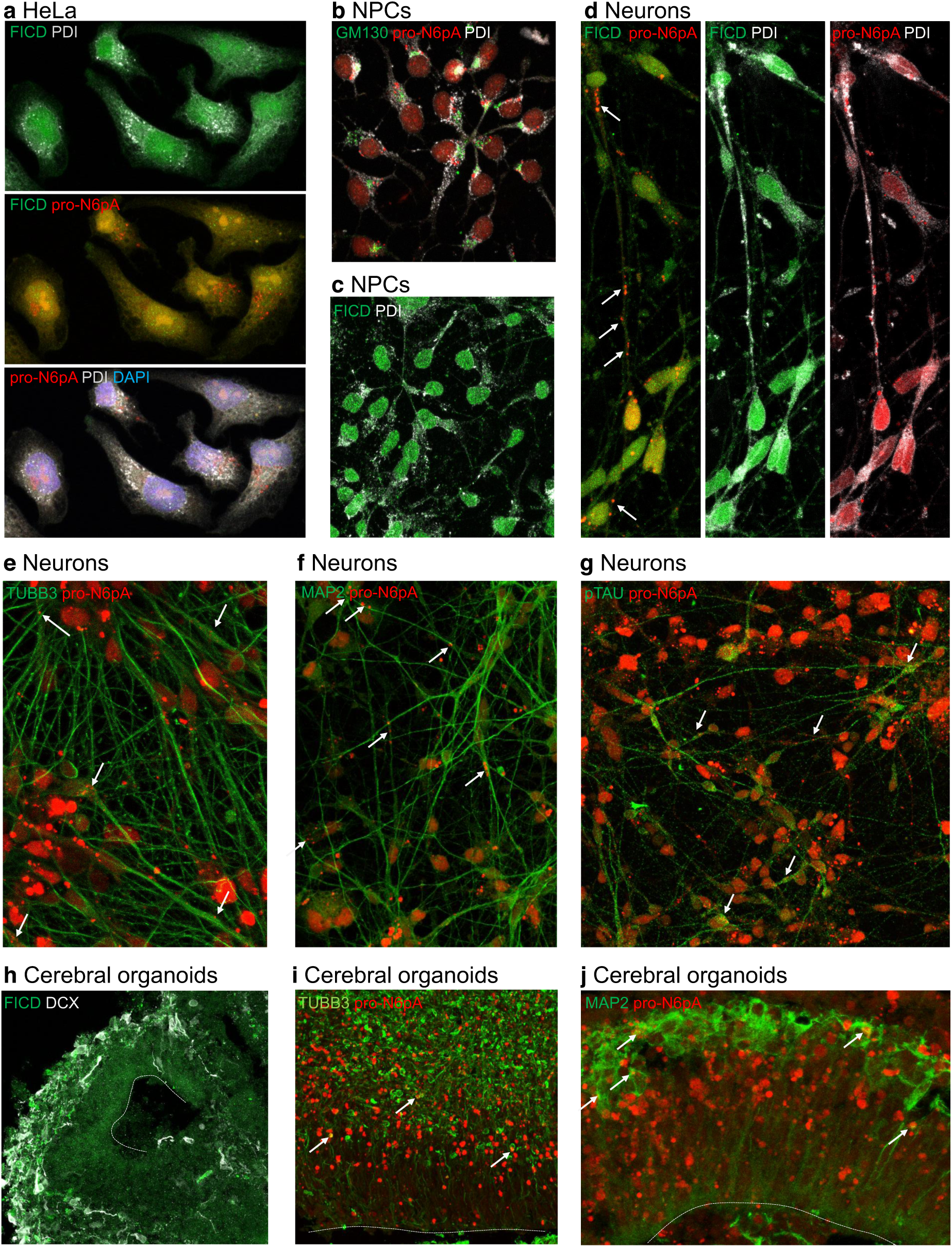
Characterization of intracellular FICD and probe localisation in HeLa (a), NPCs (b, c), neurons (d-g) and cerebral organoids (h-j). Click chemistry of **pro-N6pA** with rhodamine-azide and immunohistochemical staining. **a,** HeLa cells contain big nuclear (DAPI – blue) clusters of AMPylated proteins (pro-N6pA, red) and additional, small cytoplasmic spots of probe localisation. FICD shows characteristic ER distribution and colocalises with PDI marker, FICD rarely colocalises with probe-containing proteins. **b,** In NPCs, probe-containing AMPylated proteins (pro-N6pA, red) localize mostly to ER and only rarely with Golgi (GM130, green) with additional small nuclear and cytoplasmic clusters. **c,** FICD localizes to the ER. **d,** In differentiated neurons, clusters of AMPylated proteins localise to nucleus and processes (white arrows), both inside and outside rough ER, partly overlapping with FICD. **e,** Pro-N6pA partly colocalises with the TUBB3+ neuronal cytoskeleton (green, white arrows), with TUBB3 being identified as AMPylation target in neurons. **f,** AMPylated proteins can be found both in neuronal dendrites (MAP2+, green, white arrows indicating colocalisation; also identified as neuronal AMPylation target) and **g,** in axons (phosphoTAU+, green, white arrows for colocalisation). **h,** In cerebral organoids, FICD (green) is enriched in the DCX+ neuronal layer (white), which is in line with qPCR data from 2D in vitro generated NPCs and neurons (Fig. S19). **i, j,** AMPylated proteins are enriched right below and within the neuronal layer (TUBB3+ and MAP2+, green) and include TUBB3 (**g,** white arrows indicating colocalisation) and MAP2 (**h,** white arrows indicating colocalisation). See also Fig. S15.

Given that the cellular localisation of FICD and AMPylated proteins might play an important functional role, we combined click chemistry with rhodamine-azide for intracellular probe visualisation with immunohistochemistry (IHC) for FICD and various cellular markers: PDI for rough endoplasmic reticulum, GM130 for Golgi complex, TUBB3 for total neuronal microtubule cytoskeleton, MAP2 for neuronal dendrites, phospho-TAU for neuronal axons, and DAPI to visualize nuclei (Fig. 5, Fig. S15). Staining performed in HeLa cells revealed that AMPylated proteins are enriched in the nucleus, additional small spots were found in the cytoplasm partially overlapping with the ER. As expected, FICD is localized in the ER (Fig. 5a). This observation was further corroborated by overexpression of FICD with a C-terminal FLAG tag (Fig. S17) and by analysis of the FICD’s glycosylation using endoglycosidase H assay (Fig. S17). On the contrary, in NPCs AMPylation is strictly localized next to the rough ER and in the nucleus (Fig. 5b, c). In neurons, AMPylation was observed in nucleus and to a lesser extent in neurites, including MAP2+ dendrites and phospho-TAU+ axons (Fig. 5d-g). Finally, fibroblasts showed another specific localisation pattern with AMPylation accumulated around the nucleus and its’ complete absence inside. (Fig. S15). Differences in localisation of FICD and AMPylated proteins support the presence of additional AMP transferases with complementary cellular distribution.

### Decrease of FICD levels in *in vitro* neural model systems reveals a role of FICD-dependent AMPylation remodelling in neuronal differentiation

To understand if AMPylation plays a role in neuronal differentiation, we utilized both NPC-to-neuron differentiation and the recently developed 3-dimensional human cerebral organoids (COs)^39, 40^. COs contain areas which closely resemble the structure and organisation of the germinal zones of developing human neocortex (Fig. S18)^41^. Treatment of COs with **pro-N6pA** and subsequent analysis via LFQ LC-MS/MS confirmed the AMPylation of PFKP, found in all cell types, and CTSB, another prevalent target in other studied cell lines (Fig. 4a and Fig. S5 and S12). Analysis of the significantly enriched proteins using a STRING database revealed that several proteins are located in extracellular space (Fig. S13). Interestingly, visualization of the **pro-N6pA** probe-treated COs via click-chemistry with rhodamine-azide revealed strongest fluorescence in the neuronal layer (Fig. 5i, j and Fig. S15), which is in line with the highest number of AMPylated proteins identified in neurons (Fig. 4a).

To examine the function of AMPylation in neurogenesis and neuronal differentiation in more detail, we first characterized the expression of FICD in NPCs, neurons, neuroblastoma cells and COs and found a clear enrichment of FICD in the neurites of neurons, SH-SY5Y and in the neuronal layer of COs compared to the progenitor zone (Fig. 5d, h; Fig. S15). Results of imaging were paralleled by qPCR studies demonstrating higher baseline expression levels of FICD in neurons compared to iPSCs and NPCs (Fig. S19). We knocked down FICD levels (Fig. S20) in NPCs differentiating to neurons (Fig. 6a, b) and found a significant increase in transfected cells that remain in cell cycle (KI67+) (Fig. 6a). This result suggested a potential role of FICD-mediated AMPylation in neurogenesis. We then performed down- or upregulation of FICD expression in ventricle-like germinal zones of 50 days old COs by electroporation, as this model system better resembles the 3-dimensional organization of the developing brain. Only apical radial glia cells (aRGs), which are bipolar neural stem cells that will subsequently give rise to intermediate progenitors and neurons directly, are capable of taking up the vectors via their apical process to the ventricle-like lumen (Fig. 6c). To asses if FICD-mediated activity has a function in neurogenesis during development, COs were analysed 7 (Fig. 6 and 7, Fig. S21) and 14 days post-electroporation (dpe) (Fig. S22). Cortical-like germinal zones were defined by immunohistochemical (IHC) analysis using PAX6 as a marker for dorsal aRGs (Fig. 6d, Fig. 7b) with mitotic cells labelled for PH3 (Fig. 6e, Fig. S21). The position and number of neurons was analysed by IHC using two different markers for mature neurons: MAP2, a microtubule-associated protein which is enriched in neuronal dendrites (Fig. 7c, Fig. S21) and the nuclear marker NEUN (Fig. 6g, Fig. 7d). Most of miRNA-transfected (GFP+) cells (FICD KD) were positive for PAX6 (Fig. 6d) 7 dpe. The proportion of mitotic PH3+GFP+ cells was significantly increased (Fig. 6e, f) at the expense of neurons, as shown by the significantly reduced number of NEUN+GFP+ cells (Fig. 6g, h).

**Figure 6.**
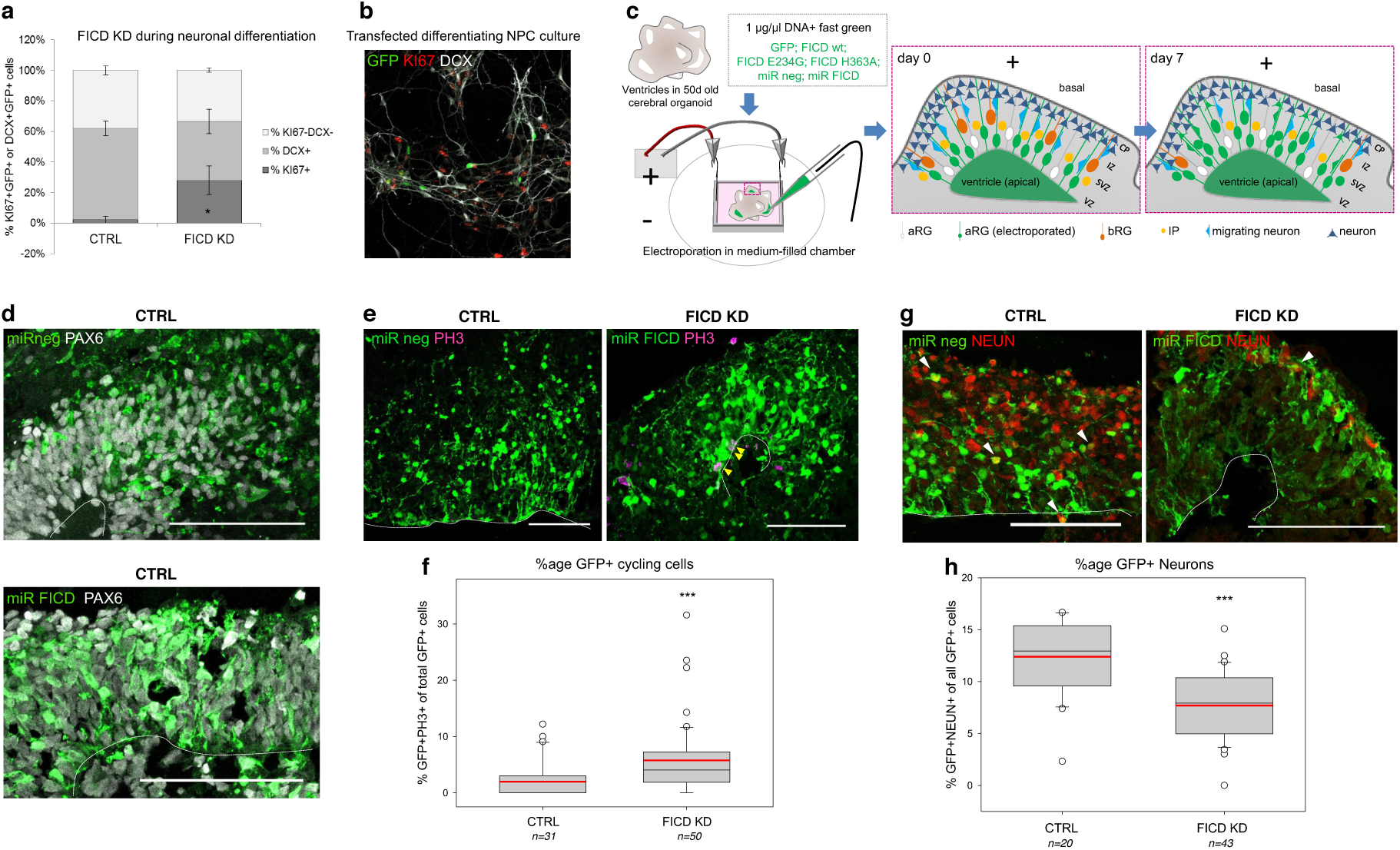
Downregulation of FICD levels keeps differentiating neurons and cerebral organoids in a cycling state. **a,** FICD knockdown (KD) in neural progenitors (NPCs) differentiating to neurons inhibits cells from cell cycle exit. NPCs were transfected with miRNAs targeting FICD (Fig. S20) and cultured under differentiating conditions for 7 days. IHC stainings for the proliferation marker KI67+ and the early neuronal marker doublecortin (DCX) showed that FICD KD leads to a significant increase in KI67+ compared to control, while the number of generated neurons tends to be decreased (analysis of 3 coverslips/condition with at least 20 transfected cells each; two-tailed Student’s *t-test:* KI67+: \**P* < 0.05; DCX+: *P = 0.068*). **b,** Example image of transfected and IHC stained culture with transfected cells (GFP+) in green, proliferating cells (KI67+) in red and differentiating neurons (DCX+) in white. **c,** Scheme showing the electroporation of DNA into ventricle-like structures of cerebral organoids (COs) and the organisation of different cell types within the germinal zone. DNA (constructs are listed; supplemented with fast green for visualisation) is injected into the lumen and taken up by aRG via their apical processes. The transfected construct can be found in IPs and neurons upon differentiation of transfected aRG (green, 7 days post electroporation (dpe)) (VZ = ventricular zone, SVZ = subventricular zone, IZ = intermediate zone, CP = cortical plate; aRG = apical radial glia, bRG = basal radial glia, IP = intermediate progenitor). **d,** Upon acute miRNA-mediated KD of FICD in ventricles of COs (50d+7), most GFP+ cells (green) have aRG identity (PAX6+, white). **e, f,** FICD KD leads to an increased number of cycling progenitors (**e,** IHC staining for PH3+ cells in M-Phase. GFP+ PH3+ cells marked by yellow arrowheads; **f,** Quantification of GFP+ PH3+ progenitor cells 7 dpe). **g, h,** aRG transfected with FICD-targeting miRNAs differentiate less to neurons (**g,** IHC staining for neuronal nuclei marker NEUN, red; GFP-positive neurons shown by white arrowheads; **h,** Quantification of GFP+ neurons 7 dpe shows significant decrease upon FICD knockdown). **d, e, g,** 50+7d old organoids; electroporated cells and their progeny shown in green; Scalebar = 50 µm, dotted line = apical surface. **f, h,** 1n = 1 electroporated germinal zone; box plot: mean (red line), median (black line), box represents 25th and 75th percentiles, whiskers extend to 10th and 90th percentiles, all outliers are shown; Significance was tested using Kruskal-Wallis One-way ANOVA on Ranks and Dunn’s Pairwise Multiple Comparison (\*\*\**P* < 0.001). See also Fig. S20 for miRNA validation, Fig. S18 for characterisation of COs, and Fig. S21 for analysis of MAP+ neuronal processes upon FICD KD in COs.

**Figure 7.**
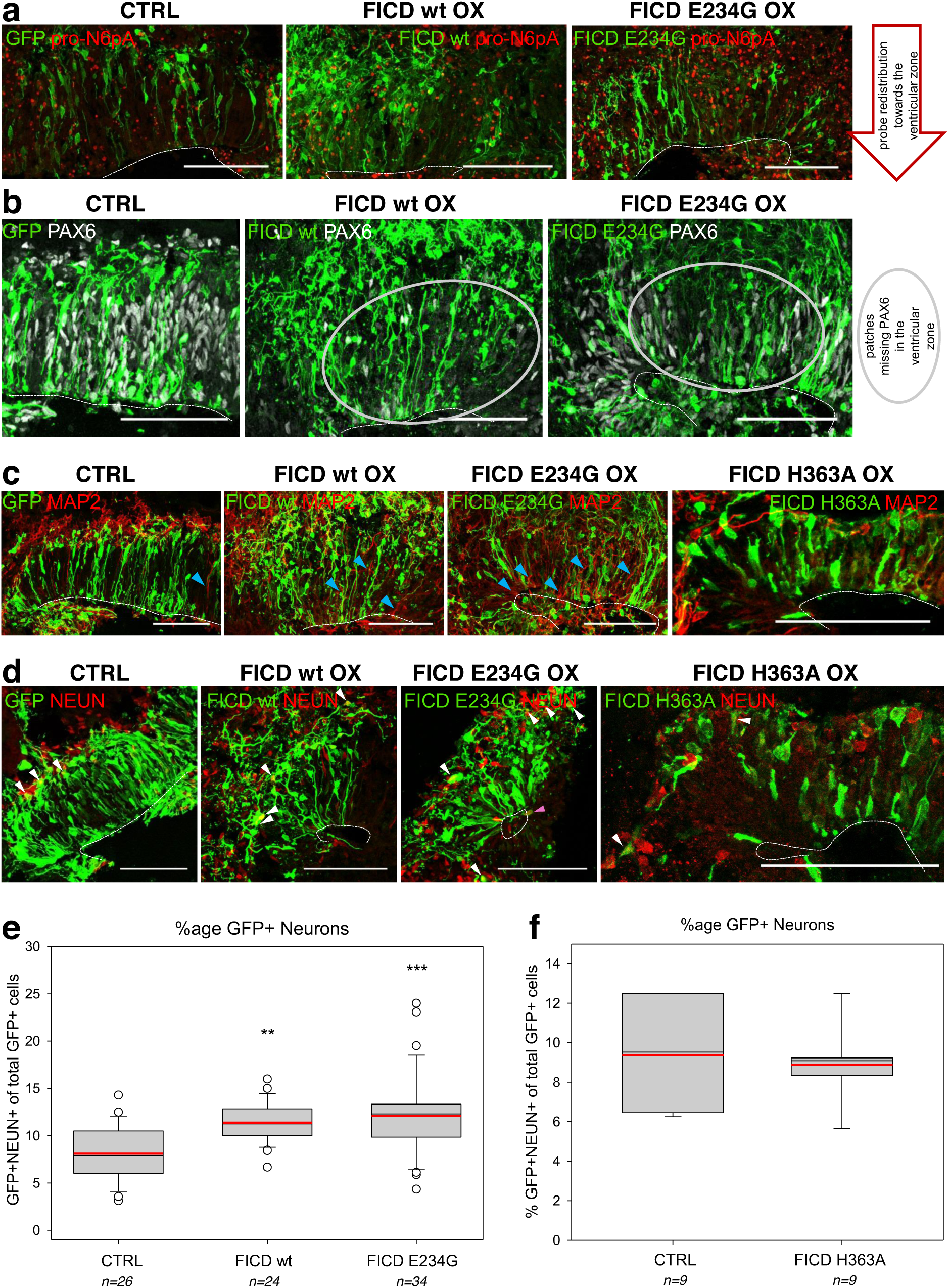
AMPylation remodelling in cerebral organoids by FICD overexpression increases neuronal differentiation. FICD wt, E234G and H363A were overexpressed in 50d old cerebral organoids (COs, see **Fig.6c** for electroporation scheme) and sections were analysed 7 days later by immunohistochemistry (IHC). FICD constructs do not bear any fluorescent tag and were co-transfected with GFP containing plasmid as a transfection control (green colour). **a,** Acute overexpression (OX) of FICD wt and E234G (green) in ventricles of COs (50+7d) leads to remodelling of AMPylation, visualised by the redistribution of fluorescence labelling using pro-N6pA (red). **b,** Germinal zones rich in cells overexpressing FICD wt or E234G (green) show patchy “holes” lacking PAX6+ dorsal NPCs (grey) in their ventricular zone (VZ) (indicated by grey circle). **c,** Upon FICD wt and E234G OX, MAP2+ neuronal processes (red) increasingly extend into the VZ (blue arrowheads), which does not occur upon control and FICD H363A OX. **d, e,** RGs overexpressing FICD wt or E234G show increased differentiation to neurons compared to control. (**d**, IHC staining for nuclei of differentiated neurons (NEUN, red; GFP+ neurons shown by white arrowheads, GFP+ neuron in the progenitor zone by blue arrowhead. **e,** Quantification of GFP+ neurons shows significant increase upon FICD wt or E234G OX). **f,** OX of the catalytically inactive FICD H363A does not lead to an increase in GFP+ neurons. **a, b, c, d,** 50+7d old organoids; electroporated cells and their progeny shown in green; Scalebar = 50 µm, dotted line = apical surface. **e, f,** 1n = 1 electroporated germinal zone; box plot: mean (red line), median (black line), box represents 25th and 75th percentiles, whiskers extend to 10th and 90th percentiles, all outliers are shown; significance was tested using Kruskal-Wallis One-way ANOVA on Ranks and Dunn’s Pairwise Multiple Comparison (\*\**P* < 0.01; *** *P* < 0.001).

### Forced expression of FICD wt and E234G *in vitro* leads to an increased neuronal differentiation

Conversely, when electroporating vectors carrying wt FICD, activated FICD E234G mutant or catalytically inactive FICD H363A mutant into ventricles of 50 days old COs, those transfected with FICD wt or E234G showed an increase and redistribution in fluorescent signal upon **pro-N6pA** treatment indicating a remodelling of AMPylation upon FICD overexpression (Fig. 7a), while there were no changes in distribution or intensity of the signal upon OX of FICD H363A used as a control (Fig. S21 and similarly in neuroblastoma cells Fig. S23). Moreover, upon FICD wt or E234G OX in COs, progenitor zones had regions sparse in PAX6+ cells (Fig. 7b). At the same time, MAP2+ neurites increasingly invaded these progenitor zones 7 dpe (Fig. 7c, blue arrowheads; Fig. S21) and 14 dpe (Fig. S22), which was not the case upon control or FICD H363A electroporation (Fig. 7c), nor upon FICD KD (Fig. S21). Interestingly, both the E234G mutant and wt FICD-transfected aRGs gave rise to a significantly higher number of neurons compared to H363A inactive mutant or control already at 7 dpe (Fig. 7e, f), which was consistent also at 14 dpe (Fig. S22). Additionally, we have excluded other cellular processes to be involved in the observed effect by whole proteome analysis of FICD transfected neuroblastoma cells (Fig. S24, Table S10), which showed only minimal changes in overall protein expression. Furthermore, pull-down of the AMPylated proteins from neuroblastoma cells under FICD E234G OX conditions revealed a general increase in AMPylation including proteins such as CTSB, TPP1, CAPZB and NSFL1C, which were found AMPylated in COs (Fig. S25, Table S11). This effect was not observed with FICD wt OX, which is in line with low AMPylation activity of the FICD wt.

Taken together, FICD may regulate the transition from neural progenitors to neurons. The direct comparison to catalytically inactive FICD H363A, showing no difference to control condition, demonstrates the importance of FICD catalytic activity in proper progenitor cell cycle exit and neuronal differentiation. These results suggest that remodelling of AMPylation may play a role in neuronal differentiation during human brain development. However, it remains to be investigated whether the specific AMPylation/de-AMPylation activity on HSPA5 and subsequent changes in the UPR are responsible for modulation of the neuronal differentiation, which would be supported by the known connection between UPR and brain development.^42^ Alternatively, the synergistic action of AMPylation on cytoskeletal protein targets catalyzed by putative AMPylators and associated changes in cellular polarization as described for example for MAP6 palmitoylation could be responsible for these effects.^43^

See also Fig. S21 for analysis of PH3+ progenitors upon FICD wt/E234G/H363A OX in COs and for scoring of MAP2+ progenitor cells intruding the VZ upon FICD KD or FICD wt/E234G/H363A OX in COs and Fig. S22 for the analysis of 2 weeks after electroporation of COs with FICD wt/E234G OX constructs.

## Discussion

Our **pro-N6pA** phosphoramidate probe design facilitated *in situ* identification of 162 potentially AMPylated proteins in different cell types and uncovered FICD as a modulator of neuronal differentiation. We successfully identified FICD dependent AMPylation as exemplified on HSPA5 and FICD independent AMPylation as shown for other proteins like PFKP, PPME and SQSTM1. FICD is the only known human AMPylator and all previous studies utilize this enzyme for deciphering substrates *in vitro*. Our *in situ* approach is global and does not only depend on FICD. Thus FICD-independent AMPylation supports the existence of additional AMP transferases such as an emerging group of pseudokinases which were identified as AMPylators in eukaryotic cells^17^. Moreover, our *in situ* profiling allowed to screen AMPylation remodelling during neuronal development from iPSCs in 2D and 3D *in vitro* approaches, connecting biological implications of FICD dependent AMPylation/de-AMPylation with human brain development: Acute KD of FICD in differentiating neurons (2D) and in aRG in cerebral organoids (3D) kept cells in a cycling state, while OX of the only known human AMPylating enzyme was shown to drive the differentiation of NPCs to neurons in cerebral organoids. The subtle but always significant dysregulation of neurogenesis resulting from FICD OX and KD may be caused by impaired AMPylation remodelling, influencing catalytic activity of metabolic enzymes or stability of cytoskeletal proteins. The remarkable number of AMPylated targets identified altogether in NPCs, neurons and COs indicates a synergistic influence in fine-tuning neurogenesis, but it is not trivial to pinpoint the function of each target protein individually, leaving the precise molecular mechanism unresolved. Furthermore, alteration of the neuronal differentiation process might be influenced as well through the AMPylation of HSPA5 and successive changes in UPR^16^.

Our study highlights both the promises and challenges of using chemical-proteomics for identification of protein PTMs. Although we have successfully identified a large group of AMPylated proteins in various cell types and elucidated its functional implications, the method itself yielded rather low rate of identified sites needed for biochemical testing of the AMPylation function *in vitro*. Nevertheless, we were able to identify seven sites and show that this PTM can inhibit target protein activity, as exemplified by CTSB, the abundance of AMPylation likely limits *in situ* detection. Future studies will thus focus on methods to quantify AMPylation levels and fine-tune enrichment and MS-based detection procedures. Interestingly, taking together both approaches of chemical-proteomics and fluorescence imaging utilizing the **pro-N6pA** probe suggests a cell type-specific AMPylation pattern. This is a combination of the particular AMPylated proteins and their intracellular localisation in a certain cell type. For example, postmitotic fibroblasts exhibit highly enriched proteins shared with the proliferating cell lines, but their subcellular localisation is very distinct from the localisation in the cycling cells. Aside the dependence of AMPylation on the cell type, we have shown with the example of thapsigargin-induced ER stress that the prevalent environmental condition can affect AMPylation.

With these features, we believe our method will lead to discovery of new functions for protein AMPylation beyond neuronal development e.g. in stem cell differentiation, unfolded protein response or regulation of complex network of cysteine cathepsins.

## Supplementary Information

## Acknowledgements

This work was supported by the European Research Council (ERC) consolidator grant (725085 - CHEMMINE), SFB749 and Alexander von Humboldt fellowship to P.K. We thank to A. Itzen for helpful suggestions and providing us with recombinant FICDs and B. F. Cravatt for providing us with the azide-TEV-cleavable-biotin linker. S. M. Hacker and A. Hoegl for manuscript proofreading, and T. Öztürk and G. Giorgio for technical assistance.

## Author Contribution

S.A.S. and P.K. conceived of the project. P.K., I.Y.B., S.C. and S.A.S. designed the experiments. P.K. and I.Y.B. performed the experiments. P.K. synthesized the probes, conducted chemical-proteomic experiments, data analysis and *in vitro* assays. I.Y.B. cultured the fibroblasts, neuroblastoma cells, iPSCs, NPCs, and neurons, performed all KD and OX experiments and analysis on COs, fluorescent imaging, qPCR, and STRING. V.C.K. performed the metabolomics experiments. N.C.B. set up the MS methods and helped with MS data analysis. M.D. generated and provided iPSC. P.K., I.Y.B., S.C. and S.A.S. wrote the manuscript.

## Author Information

## Reviewer Information

## METHODS

### Synthesis

The nucleoside (*N*^6^-propargyl adenosine, N6pA) and the rhodamine-biotin-azide tags were synthesized as described previously^44, 45^. Synthesis of the phosphoramidate probe pro-N6pA is described in the Supplementary Information following a methodology described previously^46^. Chemical identity and samples purity were established using NMR, HRMS and HPLC analysis.

### Cell lines

Human epitheloid cervix carcinoma cells (HeLa, CCL-2) and human lung carcinoma cells (A549) were cultivated in high glucose Dulbeccośs Modified Eaglés Medium (DMEM) supplemented with 10% (v/v) fetal bovine serum (FBS) and 2 mM L-glutamine. Cells were grown under a humidified atmosphere at 37 °C and 5% CO_2_. Cells were seeded into 6 cm diameter dishes and grown to 80-90% confluency. Human neuroblastoma cells SH-SY5Y (CRL-226) were cultivated in DMEM/F12 1:1 media supplemented with 10% (v/v) FBS.

### Chemical-proteomics

Cells were treated with the probes at 80-90% confluency (*n* represents number of cell culture dishes). Culture medium was removed and the cells or COs were labelled in fresh media containing 100µM N6pA or 100µM pro-N6pA (both stocks 100mM in DMSO) for 16 h at 37 °C in cells incubator. Subsequent cell lysis, click chemistry, avidin beads enrichment and MS sample preparation were performed as described previously^22–24, 47^. A total amount of 500 µg (HeLa, A549, SH-SY5Y or COs) or 250 µg (iPSCs, NPCs, neurons) of proteins in lysate was used for each MS sample preparation. For details see Supplementary Information.

### Site identifications

Site identification experiments were performed in HeLa and SH-SY5Y cells and COs. Cells or COs were cultivated and treated with pro-N6pA. After the cells lysis and protein concentration measurement, 3.6 and 16 mg of HeLa or 6 mg of SH-SY5Y or 8 mg of CO protein lysates were used for further MS sample preparations. The protocol used for enrichment and digest with TEV-cleavable linker was adapted from ref. 28. For details see Supplementary Information.

### Mass Spectrometry

Nanoflow LC-MS/MS analysis was performed with an UltiMate 3,000 Nano HPLC system coupled to an Orbitrap Fusion or Q Exactive Plus (*Thermo Fisher Scientific*). Fragments were generated using higher-energy collisional dissociation (HCD) and detected in the ion trap at a rapid scan rate. Raw files were analysed using MaxQuant software with the Andromeda search engine. Searches were performed against the Uniprot database for Homo sapiens (taxon identifier: 9606, 7th July 2015, including isoforms). At least two unique peptides were required for protein identification. False discovery rate determination was carried out using a decoy database and thresholds were set to 1 % FDR both at peptide-spectrum match and at protein levels. For AMPylation site identification spectra were searched for AMP conjugated with TEV tag (+694.2700) and only one unique or razor peptide was required. For details of MS measurement and data analysis see Supplementary Information and Tables S1, S4 and S8.

### *In vitro* CTSB activity assay

Cathepsin B (CTSB, 0.4 μg/μL, *R&D Systems*) was diluted in activation buffer (25 mM MES, 5 mM DTT, pH 5.0) to 10 μg/mL and incubated at 25 °C for 25 min. The activated CTSB was further diluted to 2 μg/mL in AMPylation buffer (20 mM Hepes, 100 mM NaCl, 5 mM MgCl_2_, 1 mM DTT, 0.1 mg/mL BSA, pH 7.5) and supplemented with 100μM ATP, 2.8 μM wt FICD or FICD E234G mutant (gift from A. Itzen, TUM) or ddH_2_O and incubated at 25 °C for 0 – 6 h. Subsequently, 3 μL of the mixture was used in 57 μL assay buffer (25 mM MES, 10 μM Z-Arg-Arg-7-amido-4-methylcoumarin hydrochloride (*Sigma*), pH 5.0) in 96-well plate and the fluorescent intensity was read by TECAN 200M Pro after 20 min using 380 nm and 460 nm as excitation and emission wavelengths.

### iPSC culture

Induced pluripotent stem cells reprogrammed from fibroblasts (for reprogramming see Supplementary Information) were cultured at 37 °C, 5 % CO_2_ and ambient oxygen level on Geltrex coated plates (*Thermo Fisher Scientific*) in mTeSR1 medium (*StemCell Technologies*) with daily medium change. For passaging, iPSC colonies were washed with PBS and incubated with StemPro Accutase Cell Dissociation Reagent (A1110501, *Life Technologies*) diluted 1:4 in PBS for 3 minutes. Pieces of colonies were washed off with DMEM/F12, collected by 5 min centrifugation at 300 × g and resuspended in mTeSR1 supplemented with 10 µM Rock inhibitor Y-27632(2HCl) (72304, *StemCell Technologies*) for the first day.

### Generation of neural progenitor cells (NPCs) and neurons from iPSCs

Neural progenitors were generated as described previously^35^ with the following modifications. Embryoid bodies (EBs) were generated from feeder-free iPSCs by incubating colonies with Collagenase Type IV (7909, *StemCell Technologies*) for 10 min, followed by washing with DMEM/F12, manual disruption and scraping with a cell lifter (3008, *Corning Life Sciences*). Resulting pieces of colonies were plated in suspension in Neural Induction Medium (NIM) consisting of DMEM/F12+Hepes (31330095, *Life Technologies*) with 1x N2 and B27 supplements (without vitamin A, *Thermo Fisher*) with medium change every other day. Resulting NPCs were passaged using Accutase (*StemCell Technologies*) and split at a maximum ratio of 1:4. NPCs were only used for up to seven passages. For differentiation to neurons, single NPCs were plated at a density of 10^4^ cells/cm^2^ on Polyornithine/Laminin plates and cultured in NPM for 1 more day to reach about 30% cell density. Afterwards, medium was changed to Neuronal Differentiation Medium NDM (NIM containing 20 ng/mL BDNF (248-BD, *R&D Systems*) and 20 ng/mL GDNF (212-GD, *R&D Systems*)) and cells were differentiated for 40 days with medium change every 5 days.

### Cerebral organoids

Cerebral organoids were generated starting from 9,000 single iPS cells/well as previously described^40^. Organoids were cultured in 10 cm dishes on an orbital shaker at 37 °C, 5% CO_2_ and ambient oxygen level with medium changes twice a week. Organoids were electroporated at 50 days after plating (see *Electroporation of cerebral organoids*) and analysed 7 and 14 dpe. For immunostaining, 16 µm sections of organoids were prepared using a cryotome. For analysis 7 dpe, 24-34 different ventricles in 7-12 organoids from 2 independent batches were analysed per construct. For 14 days, 4 organoids per construct with altogether 13-21 electroporated ventricles per construct were analysed.

### Generation and validation of microRNAs targeting FICD

MicroRNAs (miRNAs) targeting FICD were generated using the BLOCK-iT system from Invitrogen (Thermo Fisher, Waltham, MA, USA). MiRNA sequences were determined using Invitrogens RNAi design tool https://rnaidesigner.thermofisher.com/rnaiexpress/setOption.do?designOption=mirnapid=1961720787891316464, accessed on December 6th, 2017, with the NCBI Reference Sequence NM_007076.2 as seed sequence. Three miRNA sequences were chosen and ordered as oligonucleotides from Sigma. FICD miRNA oligonucleotides were annealed and ligated into a GFP-containing entry vector pENTR-GW/EmGFP-miR using T4 DNA Ligase (Thermo Fisher, Waltham, MA, USA). Subsequently, the miRNA sequences were cloned into the pCAG-GS destination vector using the Gateway system (Thermo Fisher). The resulting miRNA expression plasmids were sequenced, the knockdown efficiency was validated in Hela cells via qPCR and Westernblot and the most efficient construct was used for NPC transfection and for electroporation of COs (Fig. S19, Fig. 6).

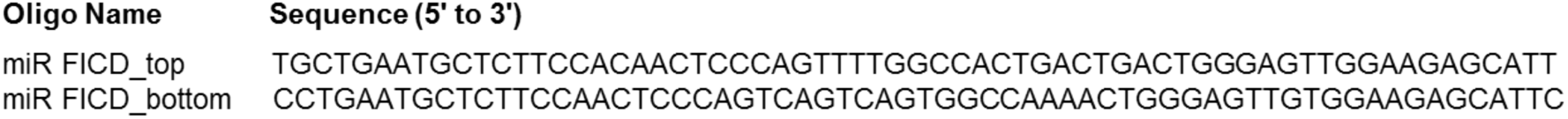

### Transfection of differentiating NPCs

For transfection of differentiating NPCs, 10^4^ cells/cm^2^ were plated on Polyornithine/Laminin-coated coverslips in 24-well plates. After one day in NPM (see *Generation of neural progenitor cells (NPCs) and neurons from iPSCs*), medium was changed to growth-factor free NIM (see *Generation of neural progenitor cells (NPCs) and neurons from iPSCs.*) to generate differentiating conditions. 4 days after plating, NPCs were transfected with 500 ng DNA/well following Lipofectamine® 3000 protocol (*Thermofisher*) and continuously cultured in NIM with medium change every other day. Cells were fixed 7 days post transfection with 4 % PFA for 20min at RT and processed by immunohistochemistry.

### Electroporation of cerebral organoids

For electroporation (see scheme in Fig. 6c), cerebral organoids were kept in NDM+A without antibiotics. The organoids were placed in an electroporation chamber (*Harvard Apparatus*) and pCMV-SPORT6 plasmid with FICD wt, FICD E234G (gift from A. Itzen, TUM), or FICD H363A plus pCAG-IRES-GFP (FICD to GFP ratio 2:1), GFP only as overexpression control, miRNA against FICD (or scrambled miRNA negative control) in pCAG-GS at a concentration of 1 µg/µl, supplemented with fast green for visualization, was injected into ventricle-like cavities at several positions per organoid. Electroporation was performed with an ECM830 electroporation device (*Harvard Apparatus*) by subjecting the organoids to a 1 second interval with 5 pulses of 50 ms duration at 80mV.

### Immunohistochemistry

Frozen organoid sections were thawn to rt for 20 min and then rehydrated in PBS for 5 min. For nuclear antigens, an antigen retrieval step (HIER) was performed in which the sections were boiled in 0.01M citric buffer pH 6 for 1 min at 720 Watt and an additional 10 min at 120 W. Slides were then left to cool down for 20 min. Half of the citric buffer was replaced by H_2_O, slides were incubated for another 10 min and then washed in PBS. Subsequently, a postfixation step of 10 min was carried out with 4 % PFA in PBS. Then, the sections were permeabilized using 0.1 % Triton X100 in PBS for 5 min. After permeabilization, sections were blocked at rt for at least 1 h with 10 % Normal Goat Serum in 0.1 % Tween in PBS. The primary antibody (Table S12 in Supplementary information) in blocking solution was then incubated overnight at 4 °C. Following several washes with 0.1 % Tween in PBS, sections were incubated with 1:1,000 dilutions of Alexa Fluor-conjugated secondary antibodies (*Life Technologies*) in blocking solution for at least 1h at rt, using 0.1 µg/ml 4,6-diamidino-2-phenylindole (DAPI, *Sigma Aldrich*) to counterstain nuclei. Finally, sections were washed again several times with 0.1 % Tween in PBS and mounted with Aqua Polymount (18606, *Polysciences*). Sections were visualized using a Leica SP8 confocal laser scanning microscope. Cells were cultured on round coverslips (13 mm diameter, *VWR*) in 24 well plates, washed with PBS and fixed with 4 % PFA in PBS for 15 min at rt. HIER, permeabilization, blocking and staining were carried out as described for the organoid sections.

### Cell quantifications

For quantification of GFP^+^ mitotic cells or neurons upon NPC transfection or in electroporated CO ventricles, all GFP^+^PH3^+^, GFP^+^KI67^+^, GFP^+^DCX^+^, and GFP^+^NEUN^+^ cells were counted using the cell counter plugin in Fiji^48^. Double positive cells were normalized to the total number of GFP^+^ cells.

### Statistics

Statistical analysis of the MaxQuant result table proteinGroups.txt (Table S2) was done with Perseus 1.5.1.6. Putative contaminants and reverse hits were removed. Dimethyl-labelling ratios or normalized LFQ intensities were log_2_-transformed, hits with less than 3 valid values in each group were removed and −log_10_(*p*-values) were obtained by a two-sided one sample Student’s *t*-test over replicates with the initial significance level of *p* = 0.05 adjustment by the multiple testing correction method of Benjamini and Hochberg (FDR = 0.05), the −log_10_ of *p*-values were plotted against the log_2_ of geometric mean of ratios “heavy”/”light” (H/L) for dimethyl labelling or by volcano plot function for LFQ. Distance from zero was calculated from significance and fold enrichments from respective volcano plot as *d* = √((*fold enrichment*)^9^ + (*significance*)^9^). Venn diagrams were generated with a drawing tool at http://bioinfogp.cnb.csic.es/tools/venny/ using gene names as a key. All graphs were processed in Microsoft Excel or OriginPro 2017. Statistics for qPCR data and quantifications of immunohistochemical stainings in cells and COs was performed in SigmaPlot (Version 13.0; Systat Software, San Jose, CA) using Kruskal-Wallis ANOVA on Ranks with Dunn’s Pairwise Multiple Comparison. For NPC transfection, 3 coverslips with at least 20 transfected cells each were analysed. For COs, 2-4 batches of organoids were analysed for each construct (Data shown with *n* = total number of electroporated ventricles analysed per construct).

### Data availability

The mass spectrometry proteomics data have been deposited to the ProteomeXchange Consortium via the PRIDE partner repository with the dataset identifier PXD015062 reviewer account detail, username: password: 17wEcc5V.

