## Supplementary Information for "FICD activity and AMPylation remodelling modulate human neurogenesis"

### CONTENTS

**(A) Supplementary Figures**

**(B) Structures and synthesis**

**(C) Biological methods**

**(D) NMR spectra**

**(E) References**

### (A) Supplementary Figure

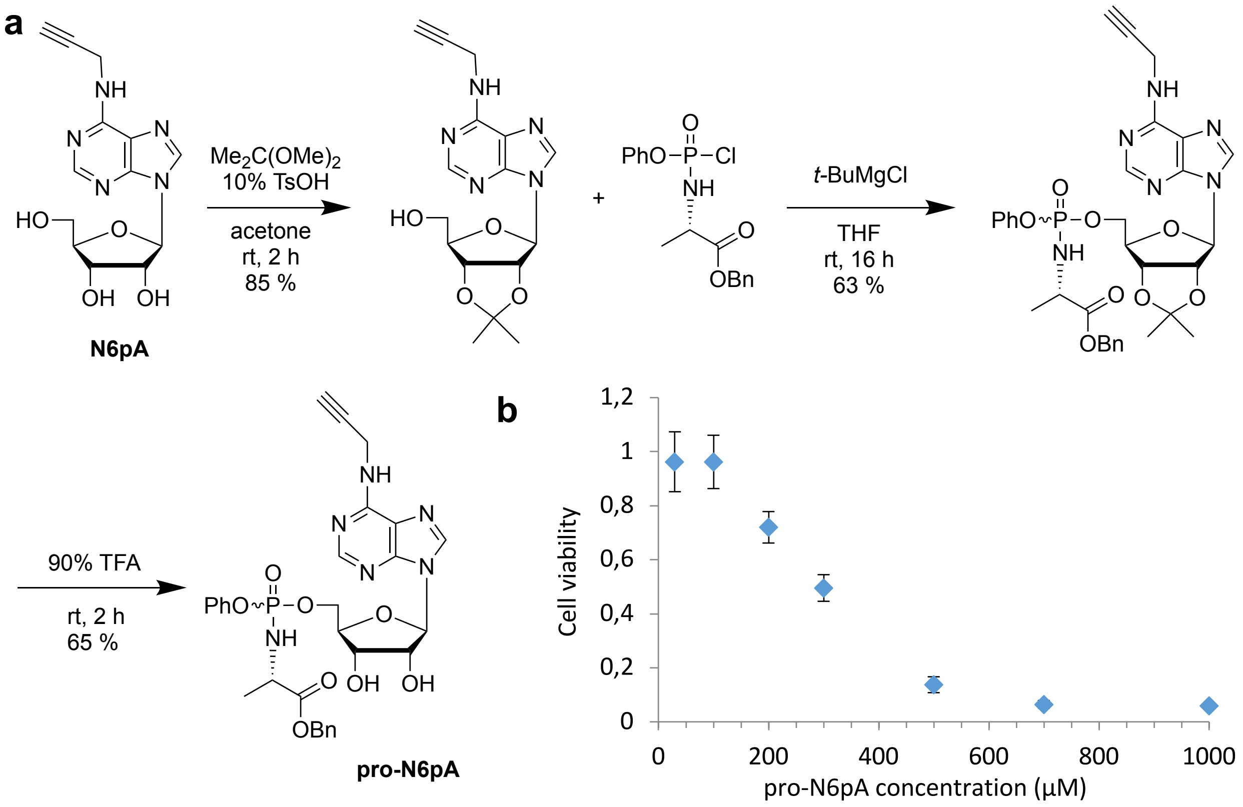

**Figure S1 | Synthesis and cytotoxicity of pro-N6pA. a,** Synthetic approach to pronucleotide probe. **b**, Cytotoxicity MTT assay of **pro-N6pA** probe. Dots represent the means of four replicates with standard deviation.

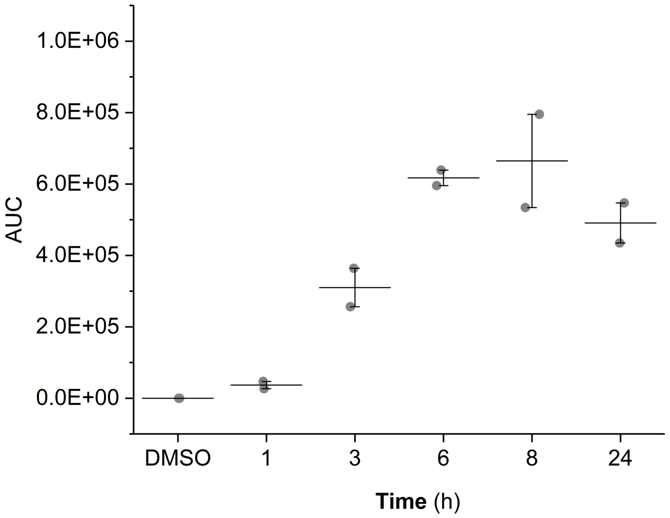


**Figure S2 | *In situ* metabolic activation of the pro-N6pA probe to N6pATP.** HeLa cells were treated with 100 µM pro-N6pA for indicated period of time. Horizontal represent the mean of the duplicates.

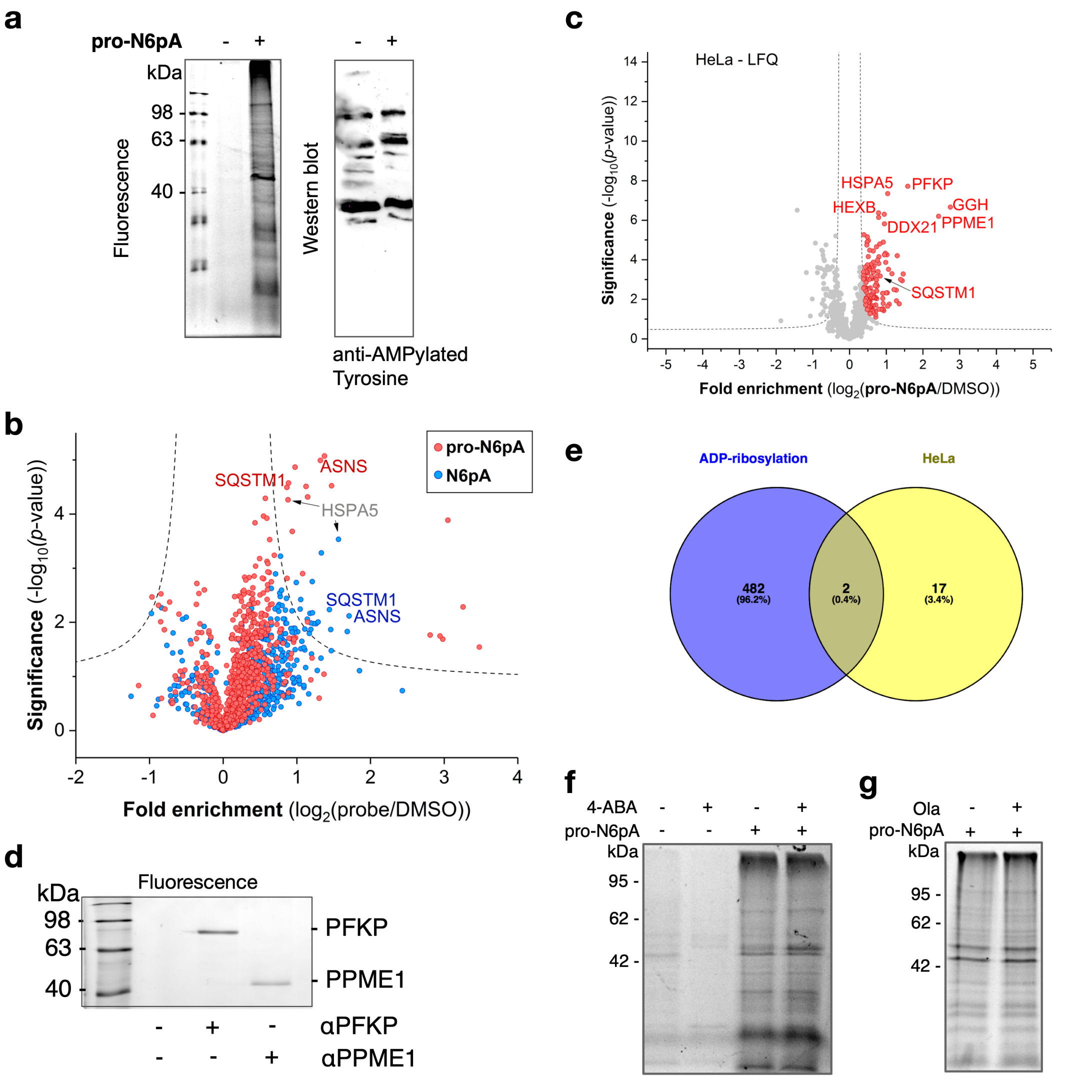


**Figure S3 | Validation of phosphoramidate pronucleotide probe pro-N6pA as probe for cellular AMPylation. a**, Direct comparison of the pro-N6pA labelling in HeLa cells with immunostaining of the western blot using anti-AMPylated tyrosine antibody. **b**, Volcano plot visualization of fold-enrichment using pro-N6pA or parent nucleoside (N6pA) compared to DMSO (control) versus significance upon performing a two-sample *t*-test (FDR 0.05, s0 0.3; *n* = 12 (pro-N6pA) or 11 (N6pA)). Pro-N6pA or N6pA treatment with 100 µM probe, 16 h incubation. **c**, Volcano plot visualising the pro-N6pA identified targets using LFQ method in HeLa cells. **d**, Fluorescence SDS-PAGE showing immunoprecipitated PFKP and PPME1 from pro-N6pA treated HeLa cells. Immunoprecipitated proteins were labelled via click-chemistry with rhodamine-azide. **e**, Venn plot representing overlap of known ADP-ribosylated proteins and hits identified by pro-N6pA probe in HeLa cells. **f**, Inhibition of poly(ADP-ribose) polymerases by pre-treatment of the HeLa cells with 20 mM 4-aminobenzamide (4-ABA) for

2 h before addition of pro-N6pA. **g**, Inhibition of poly(ADP-ribose) polymerases by pre-treatment of the HeLa cells with 10 µM Olaparib (Ola) for 2 h before addition of pro-N6pA.

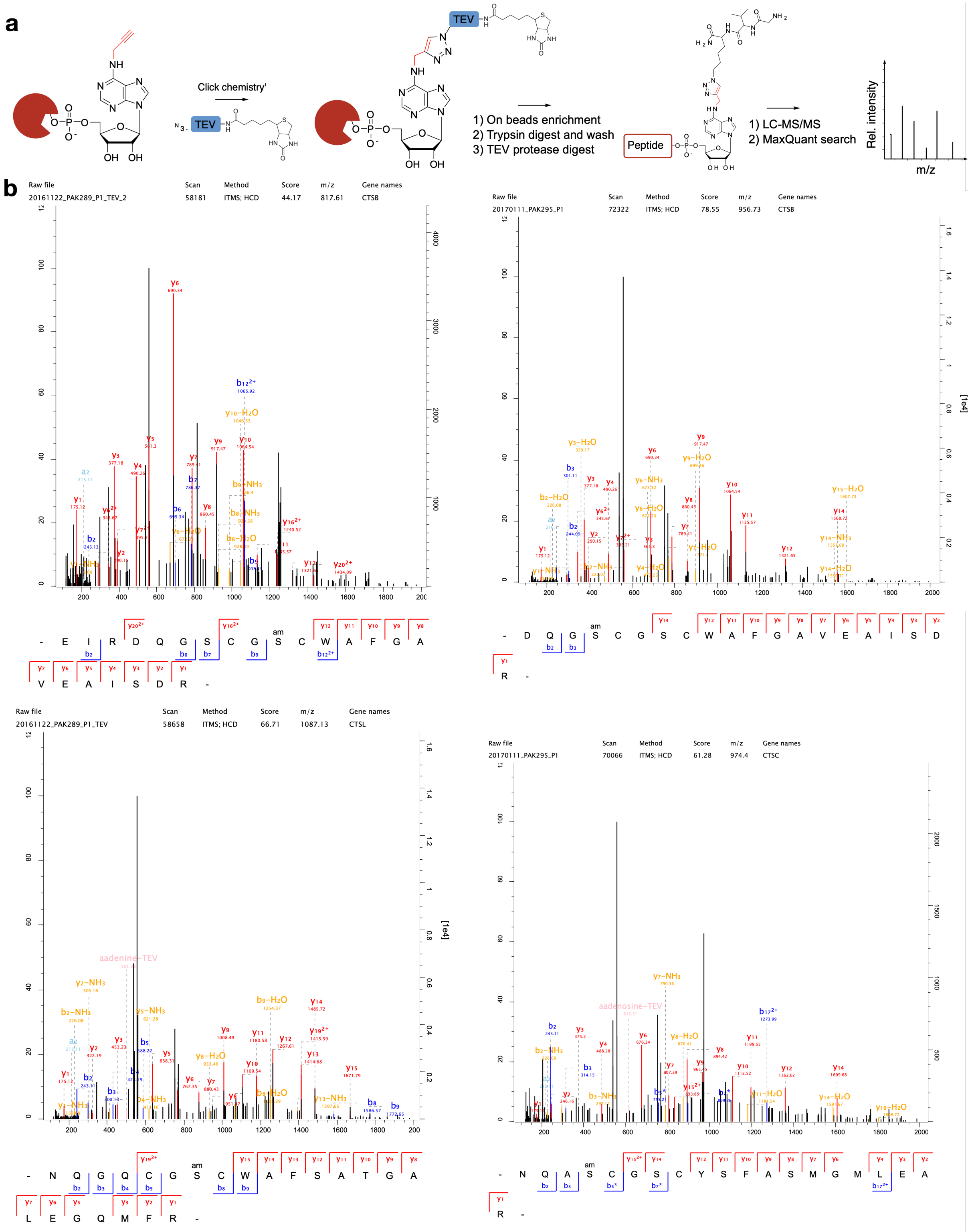

**c**

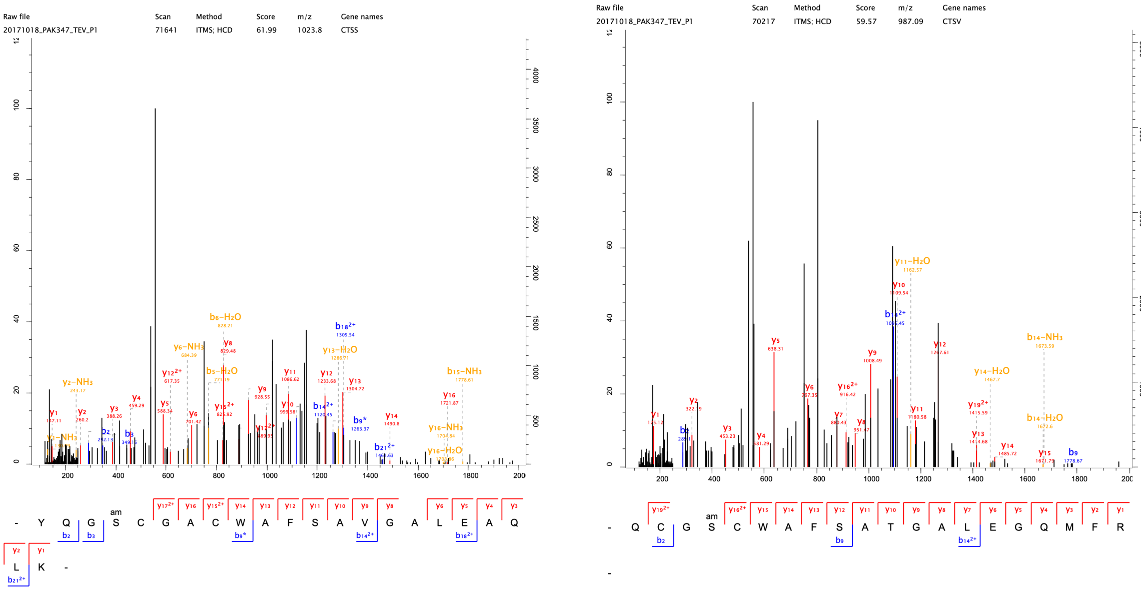


**d**

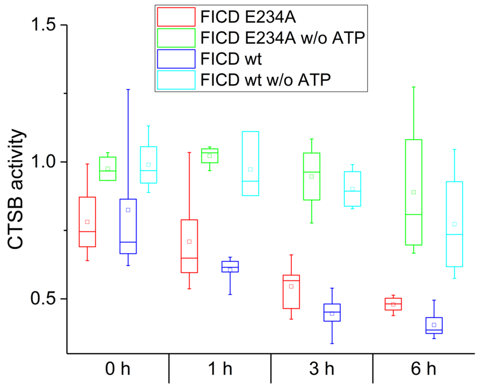


**Figure S4 | AMPylation site identification on cathepsins in HeLa cells and inhibition of CTSB activity upon AMPylation by FICD. a**, Scheme of site identification using the TEV-cleavable-linker. **b**, MS/MS spectra (MaxQuant) for the CTSB, CTSC and CTSL AMPylation site identification. CTSC AMPylation site was identified by two MS/MS scans with PEP scores 0.0224 and 0.0168. Additional sites on cathepsins were identified in SH-SY5Y cells and COs (Table S4). **c**, MS/MS spectra (MaxQuant) for the CTSS and CTSV identified in COs. **d**, *In vitro* CTSB peptidase activity upon incubation with wt FICD or E234A mutant at 25 °C for 0 – 6 h with or without ATP (0.1 mM). Boxes represent 25-75 percentile, whiskers stand for min and max values, square is mean value and line is median.

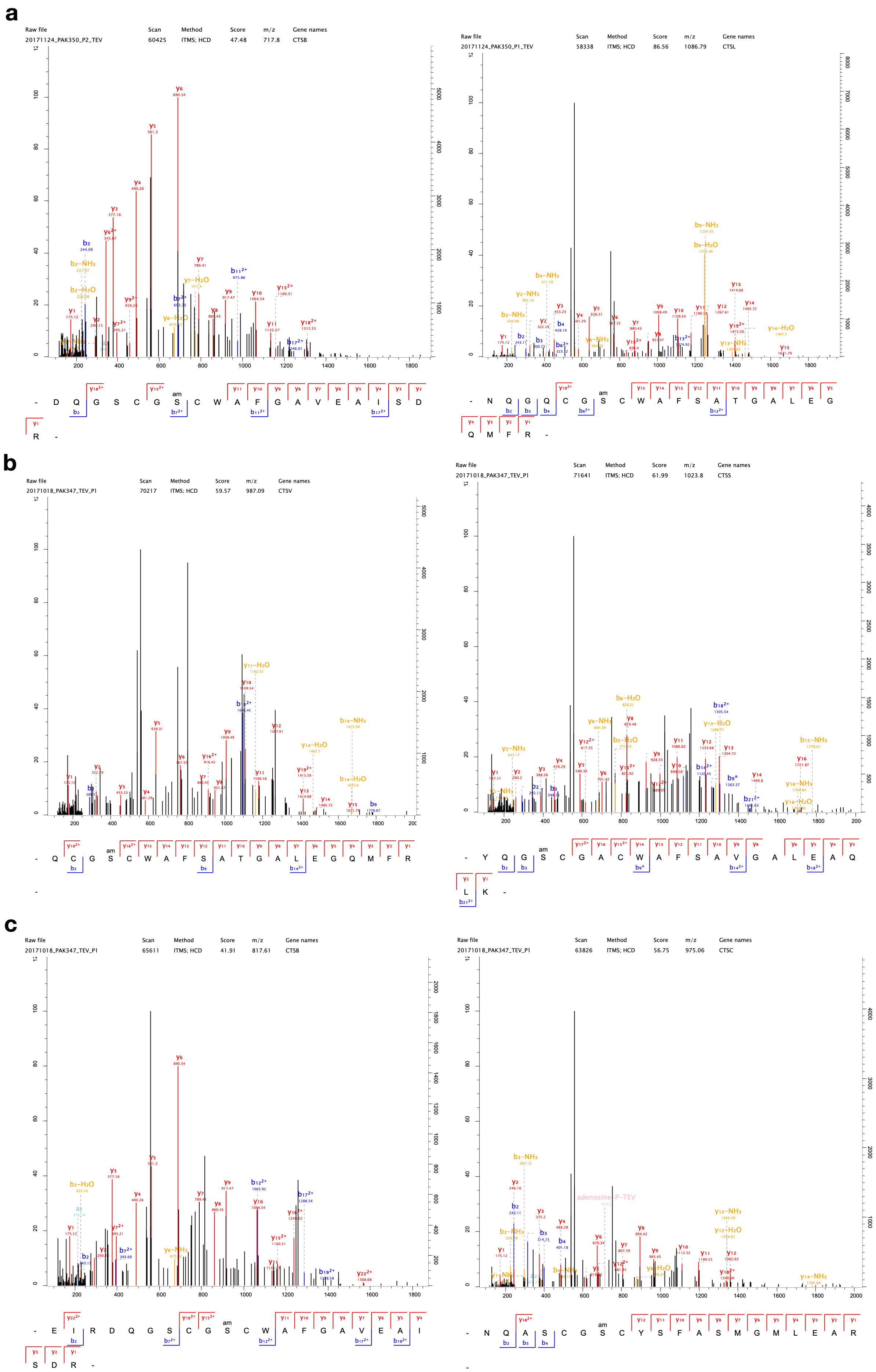
a
Raw file Scan Method Score m/z Gene names
20171124_PAK350_P2_TEV 60425 ITMS; HCD 47.48 717.8 CTSB
Raw file Scan Method Score m/z Gene names
20171124_PAK350_P1_TEV 58338 ITMS; HCD 86.56 1086.79 CTSL
b
Raw file Scan Method Score m/z Gene names
20171018_PAK347_TEV_P1 70217 ITMS; HCD 59.57 987.09 CTSV
Raw file Scan Method Score m/z Gene names
20171018_PAK347_TEV_P1 71641 ITMS; HCD 61.99 1023.8 CTSS
c
Raw file Scan Method Score m/z Gene names
20171018_PAK347_TEV_P1 65611 ITMS; HCD 41.91 817.61 CTSB
Raw file Scan Method Score m/z Gene names
20171018_PAK347_TEV_P1 63826 ITMS; HCD 56.75 975.06 CTSC
adenosine P-TEV

**Figure S5 | AMPylation sites identified in SH-SY5Y cells and cerebral organoids. a**, MS/MS spectra (MaxQuant) for the CTSB, CTSL AMPylation sites from SH-SY5Y cells. **b**, Cathepsins CTSV, CTSS AMPylation sites identified in cerebral organoids. **c**, Cathepsins CTSB, CTSC AMPylation sites identified in cerebral organoids (Table S4).

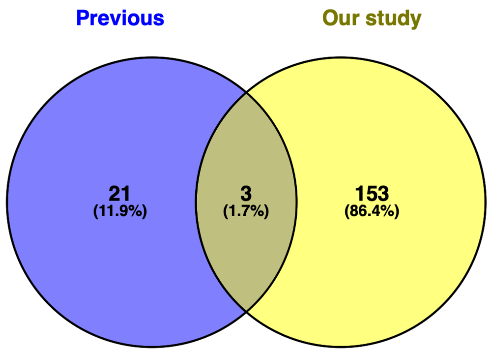


**Figure S6 | Comparison of proteins identified in previous studies[20] and our study.**

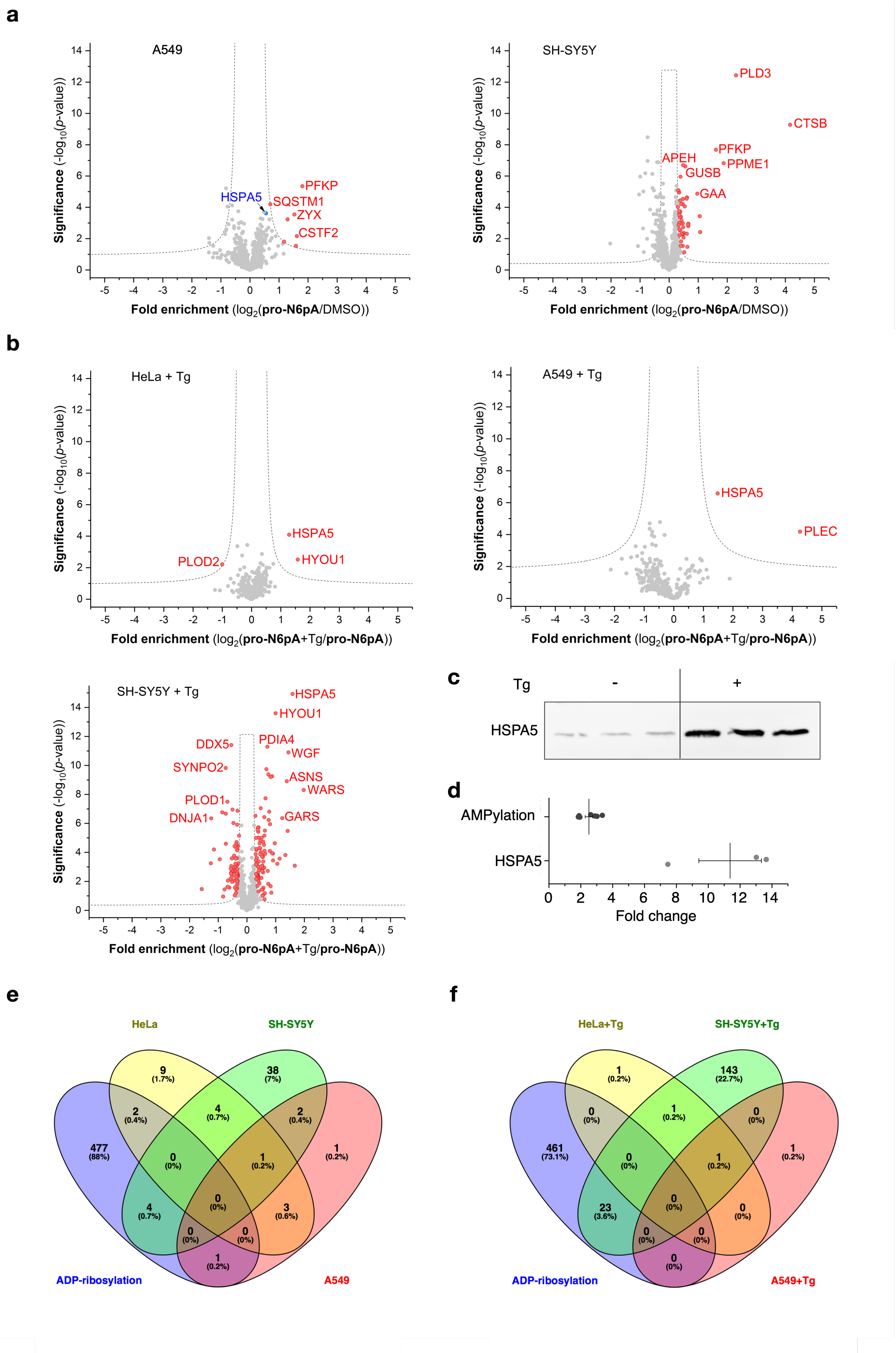

a
A549
HSPA5
PFKP
SQSTM1
ZYX
CSTF2
SH-SY5Y
PLD3
CTSB
PFKP
PPME1
APEH
GUSB
GAA
Significance (-log10(p-value))
Fold enrichment (log2(pro-N6pA/DMSO))
b
HeLa + Tg
HSPA5
HYOU1
PLOD2
A549 + Tg
HSPA5
PLEC
Fold enrichment (log2(pro-N6pA+Tg/pro-N6pA))
SH-SY5Y + Tg
HSPA5
HYOU1
DDX5
PDIA4
WGF
SYNPO2
ASNS
WARS
PLOD1
GARS
DNJA1
c
Tg
-
+
HSPA5
d
AMPylation
HSPA5
Fold change
e
HeLa
SH-SY5Y
ADP-ribosylation
A549
f
HeLa+Tg
SH-SY5Y+Tg
ADP-ribosylation
A549+Tg

**Figure S7 | AMPylation is dependent on cell type and changes upon ER stress induction. a, b,** Pro-N6pA 100 μM probe, 16 h incubation. Complete list of the proteins identified by chemical-proteomic experiments is in Table S1 and S8. **a**, Volcano plot visualizations of fold-enrichment using pro-N6pA compared to DMSO *versus* significance upon performing a two-sample *t*-test (FDR 0.05, s0 0.3; *n* = 8) from *in situ* labelling of A549 and SH-SY5Y cells. The MS intensities were quantified in A549 via DiMe, in SH-SY5Y via LFQ. **b**, Volcano plot visualizations of fold-enrichment using pro-N6pA upon Tg treatment (1 µg/mL, 24 h) compared to pro-N6pA treated cells *versus* significance upon performing a two-sample *t*-test (FDR 0.05, s0 0.3; *n* = 8 or 7 (HeLa + Tg)) from *in situ* labelling of HeLa, A549 and SH-SY5Y cells. The MS intensities were quantified in HeLa + Tg via DiMe, in A549 and SH-SY5Y via LFQ. **c**, Immunoblot of endogenous HSPA5 levels in HeLa cells without or with Tg treatment (1 µg/mL, 24 h). **d**, Graph representation of fold change in AMPylation and HSPA5 expression levels after Tg treatment based on normalized MS intensities (MaxQuant) and immunoblot analysis, respectively. **e**, **f**, Venn diagrams show minor overlap of known ADP-ribosylated proteins with identified AMPylated proteins in different cell lines with or without induction of ER stress by Tg.

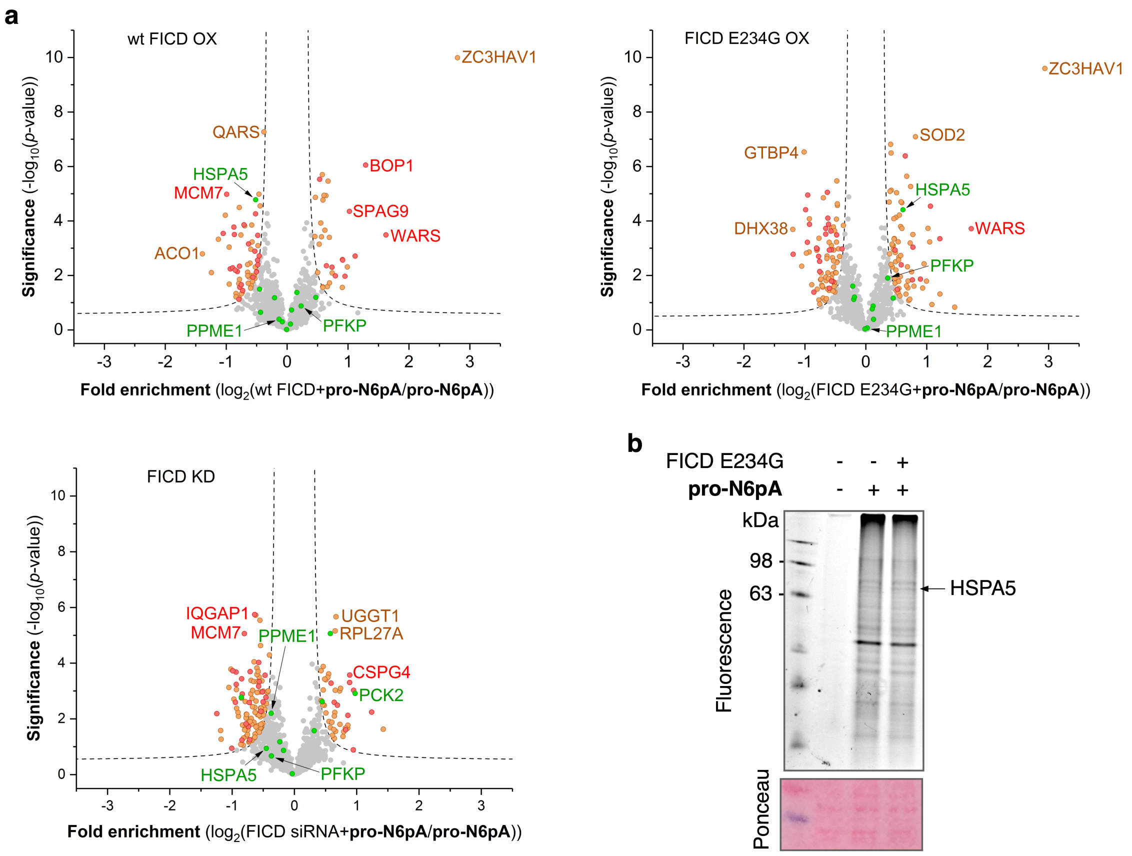


**Figure S8 | Volcano plots of FICD overexpression and knockdown in HeLa cells. a**, Volcano plot of fold-enrichment by pro-N6pA in HeLa transfected with FICD or siRNA compared to pro-N6pA treated cells versus significance upon performing a two-sample *t*-test (FDR 0.05, s0 0.3; *n* = 8). Green dots represent proteins identified as AMPylated in HeLa cells. Red dots represent proteins which overlap for the presented three volcano plots. Orange dots represent other significantly enriched proteins see **Table S6**. **b**, Overexpression of FICD E234G in HeLa cells. In-gel analysis of the labelling with pro-N6pA probe.

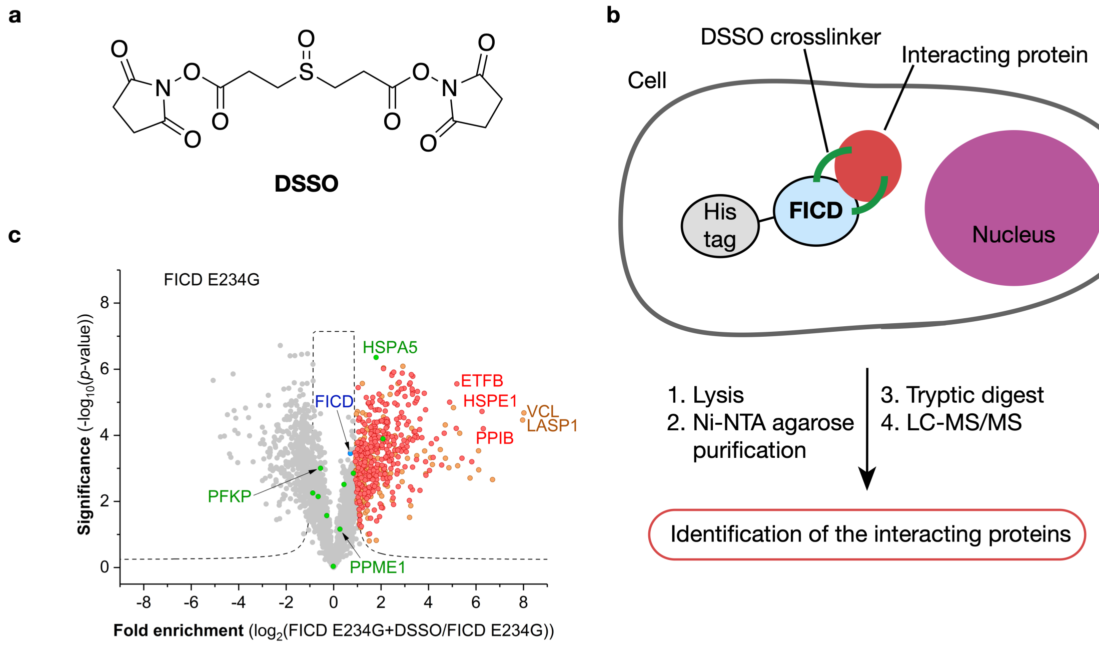


**Figure S9 | Pull-down of FICD interacting proteins using DSSO crosslinker. a**, Structure of the DSSO crosslinker. **b**, Workflow of identification of the FICD interacting proteins using the DSSO crosslinker. **c**, Volcano plot representing FICD E234G enriched proteins from HeLa cells. Green dots represent the proteins found AMPylated in HeLa cells (FDR 0.01; s0 1.5; *n* = 3). Red dots represent proteins which were significantly enriched in the parallel experiment with the wt FICD. Orange dots represent proteins significantly enriched only in this experiment see **Table S7**.

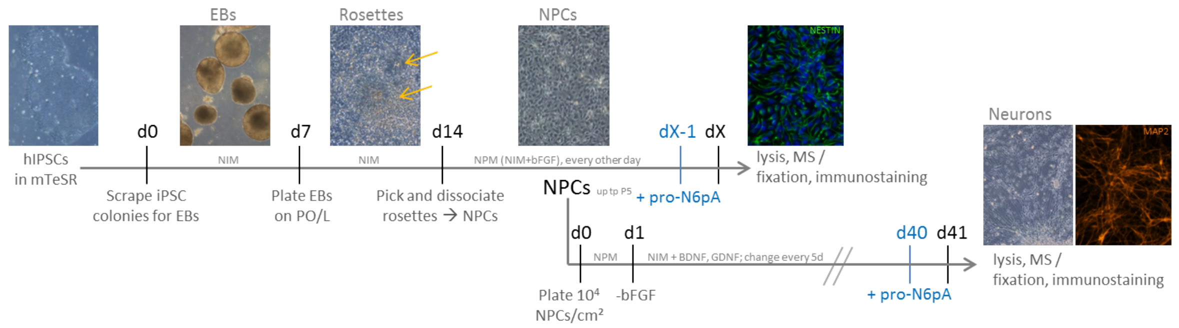


**Figure S10 | NPCs and neuron preparation.** Timeline for generation of Neural progenitor cells (NPCs) and neurons (according to ref. 35) and their treatment with pro-N6pA (neural rosettes: yellow arrows; hIPSCs = human induced pluripotent stem cells; EBs = embryoid bodies; PO/L

= poly-ornithine-laminin coated plates; NIM = neural induction medium; NPM = neural progenitor medium).

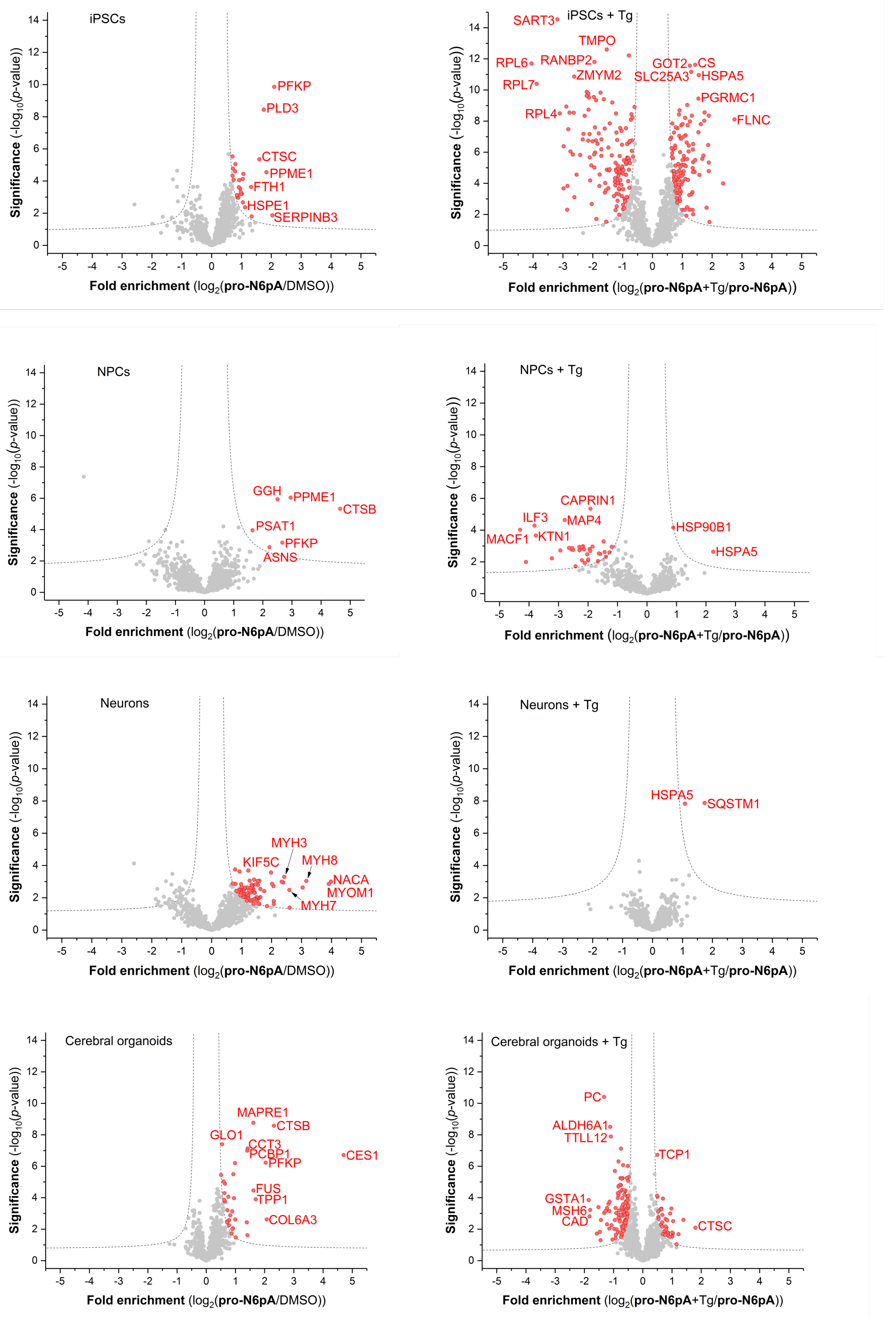
iPSCs
PFKP
PLD3
CTSC
PPME1
FTH1
HSPE1
SERPINB3
Significance (-log$_{10}$($p$-value))
Fold enrichment (log$_2$(pro-N6pA/DMSO))
iPSCs + Tg
SART3
TMPO
RPL6
RANBP2
ZMYM2
RPL7
RPL4
GOT2
CS
SLC25A3
HSPA5
PGRMC1
FLNC
Fold enrichment (log$_2$(pro-N6pA+Tg/pro-N6pA))
NPCs
GGH
PPME1
CTSB
PSAT1
PFKP
ASNS
NPCs + Tg
CAPRIN1
ILF3
MAP4
MACF1
KTN1
HSP90B1
HSPA5
Neurons
MYH3
KIF5C
MYH8
NACA
MYOM1
MYH7
Neurons + Tg
HSPA5
SQSTM1
Cerebral organoids
MAPRE1
CTSB
GLO1
CCT3
PCBP1
PFKP
CES1
FUS
TPP1
COL6A3
Cerebral organoids + Tg
PC
ALDH6A1
TTLL12
TCP1
GSTA1
MSH6
CAD
CTSC

**Figure S11 | Neurons show specific AMPylation of cytoskeletal and motor proteins. a**, Volcano plot visualizations of fold-enrichment using pro-N6pA with or without Tg treatment (1 µg/mL, 24 h) compared to DMSO or pro-N6pA treated cells versus significance upon performing a two-sample *t*-test (FDR 0.05, s0 0.3; *n* = 9) from *in vitro* labelling of iPSCs, NPCs, neurons and COs.

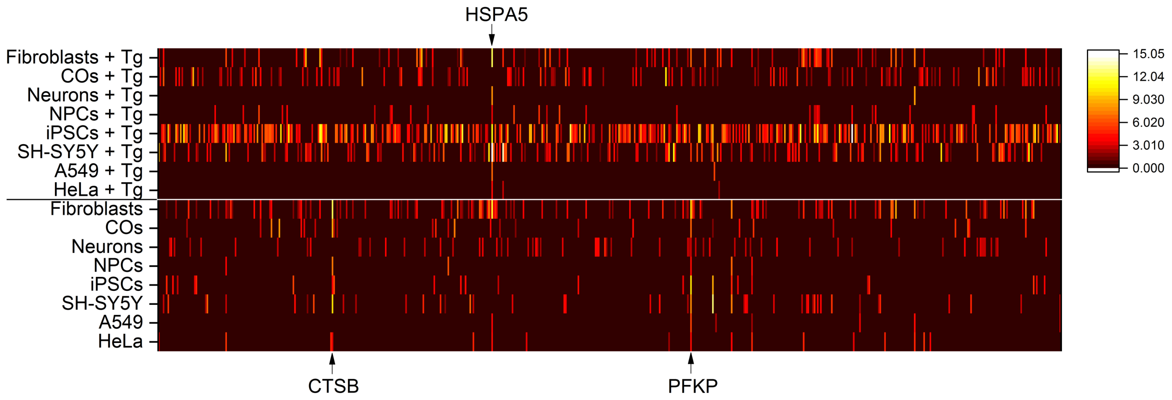


**Figure S12 | Heat map summarising the data from all tested cell lines**. **a**, Heatmap representation of enriched proteins identified in different cell types and COs; Colour represents the proteins' distance from zero of enriched proteins from respective volcano plot (FDR 0.05, s0 0.3; *n* = 7 – 9). **Table S1, S8**.

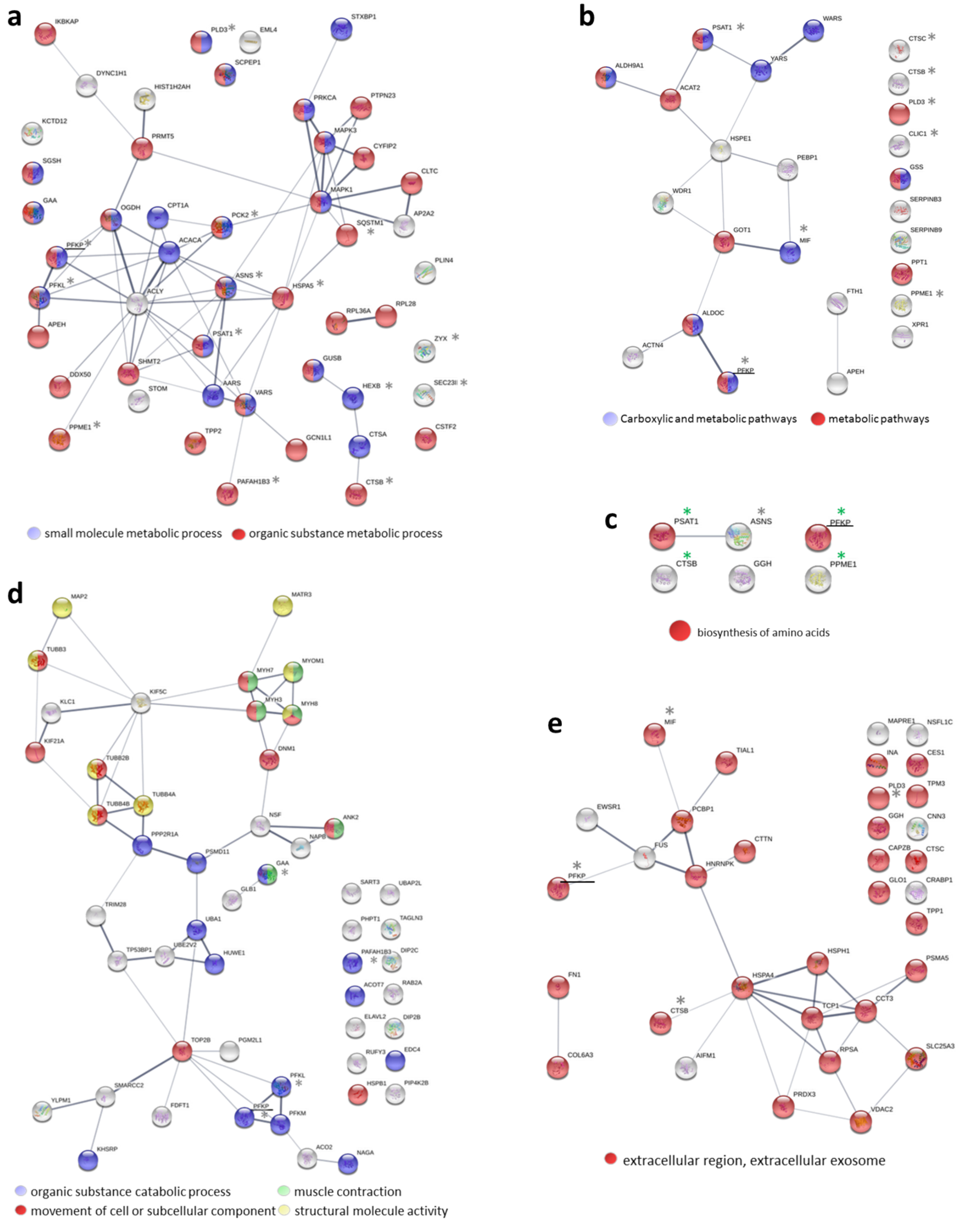


**Figure S13 | Network and GO Term Analysis of AMPylated Proteins.** Network analysis of AMPylated (and not ADP-ribosylated) hits in cancer lines (a), iPSCs (b), NPCs (c), neurons (d), and COs (e) with examples of enriched GO Terms highlighted in different colours and stars marking recurrent hits (PFKP underlined as only hit found in all analysed cell types;

green stars for hits in NPCs already present in iPSCs). Network Analysis was performed in STRING.db. See also **Table S13**.

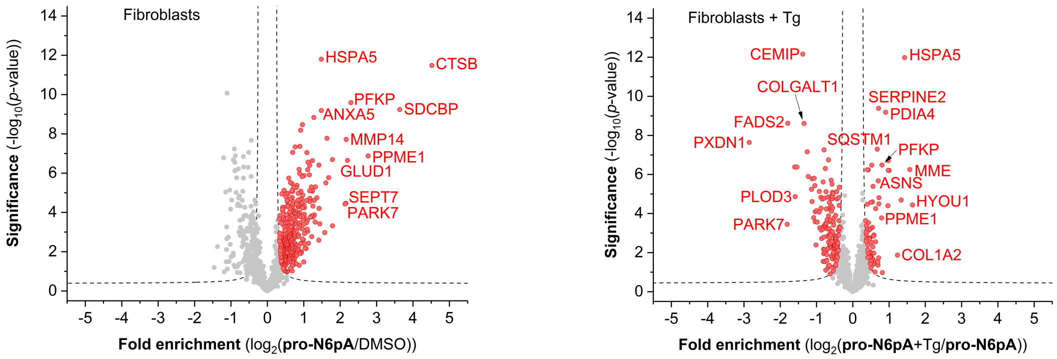


**Figure S14 | Fibroblasts chemical proteomics.** Volcano plot of fold-enrichment in fibroblasts by **pro-N6pA** labelling compared to DMSO or **pro-N6pA** + Tg versus significance upon two-sample *t*-test (FDR 0.05; s0 0.3; $n$ = 9).

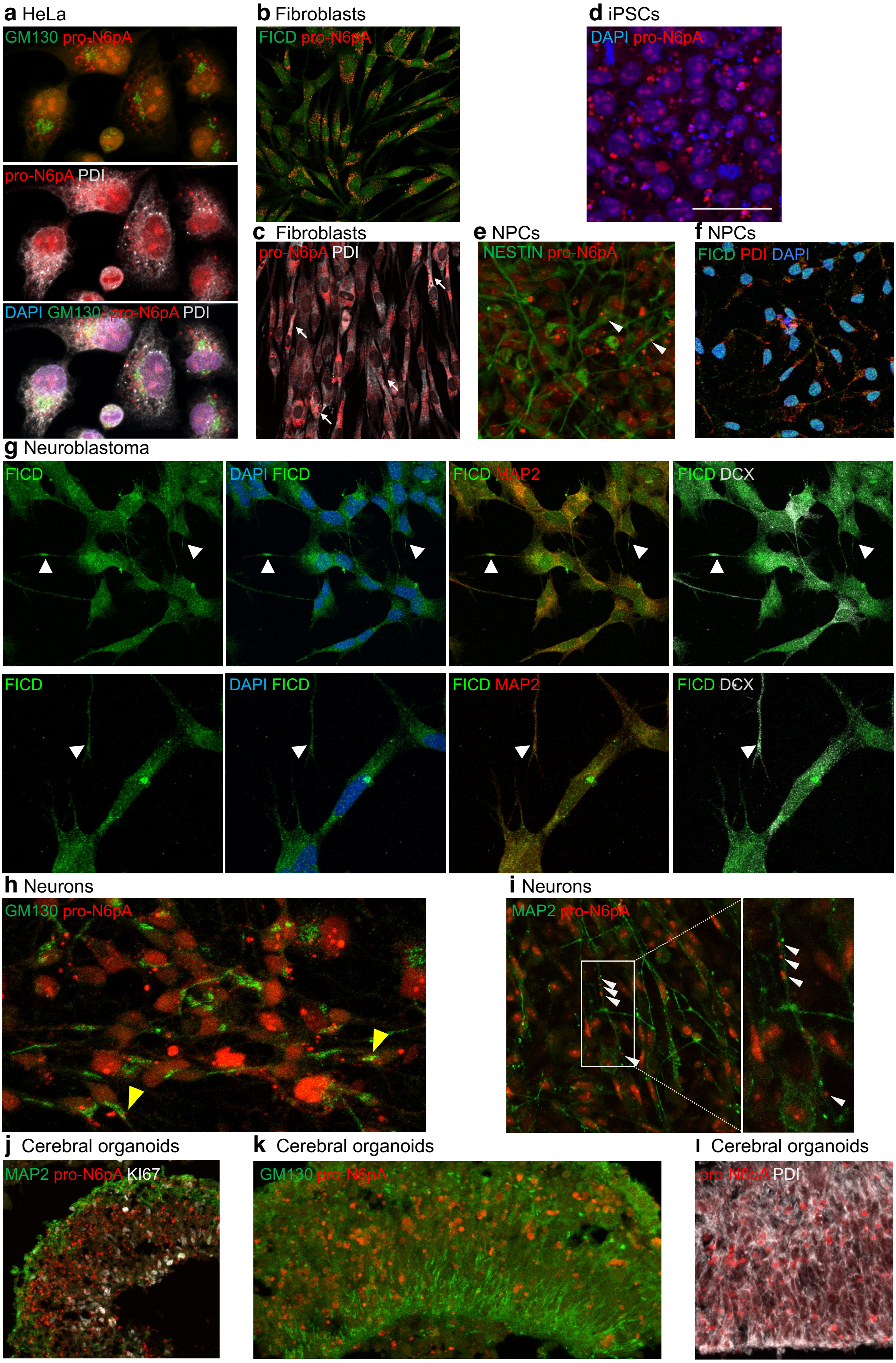
a HeLa
GM130 pro-N6pA
pro-N6pA PDI
DAPI GM130 pro-N6pA PDI
b Fibroblasts
FICD pro-N6pA
c Fibroblasts
pro-N6pA PDI
d iPSCs
DAPI pro-N6pA
e NPCs
NESTIN pro-N6pA
f NPCs
FICD PDI DAPI
g Neuroblastoma
FICD
DAPI FICD
FICD MAP2
FICD DCX
FICD
DAPI FICD
FICD MAP2
FICD DCX
h Neurons
GM130 pro-N6pA
i Neurons
MAP2 pro-N6pA
j Cerebral organoids
MAP2 pro-N6pA KI67
k Cerebral organoids
GM130 pro-N6pA
l Cerebral organoids
pro-N6pA PDI

**Figure S15 | Characterization of intracellular FICD and probe localisation in HeLa, fibroblasts, iPSCs, NPCs, neuroblastoma, neurons and cerebral organoids.** **a**, AMPylated proteins (red) localise to the nucleus and rough ER (PDI, white) in HeLa, but not to cis-Golgi (GM130, green). **b**, In fibroblasts, FICD (green) is localised in the nucleus and cytoplasm, including processes. AMPylated proteins (red; b, c) accumulate in the perinuclear area, including ER (PDI, white, white arrows). **c**, with additional probe localisation to the processes, while the nucleus is free of probe signal. **d**, In iPSCs, AMPylated proteins form nuclear clusters similar to those in HeLa. **e**, Probe localises to NESTIN+ (green) processes (white arrowheads) in NPCs. **f**, FICD (green) is both in the nucleus and in processes of NPCs, partly colocalising with rough ER (PDI, red). **g**, IHC of FICD in SH-SY5Y neuroblastoma cells overlaid with DAPI, DCX and MAP2 markers reveals FICD localisation in cytoplasm and neurites (white arrowheads). **h**, Probe rarely localises to Golgi (GM130, green) in neurons (yellow arrowheads). **i**, Additional example showing the localisation of AMPylated proteins in MAP2+ (green) neuronal processes (white arrowheads). **j-l**, In cerebral organoids, AMPylated proteins accumulate basally, below and within the neuronal layer (MAP2+, green, j), with some degree of colocalisation with Golgi (GM130, green; k) and more with ER (PDI, white; l).

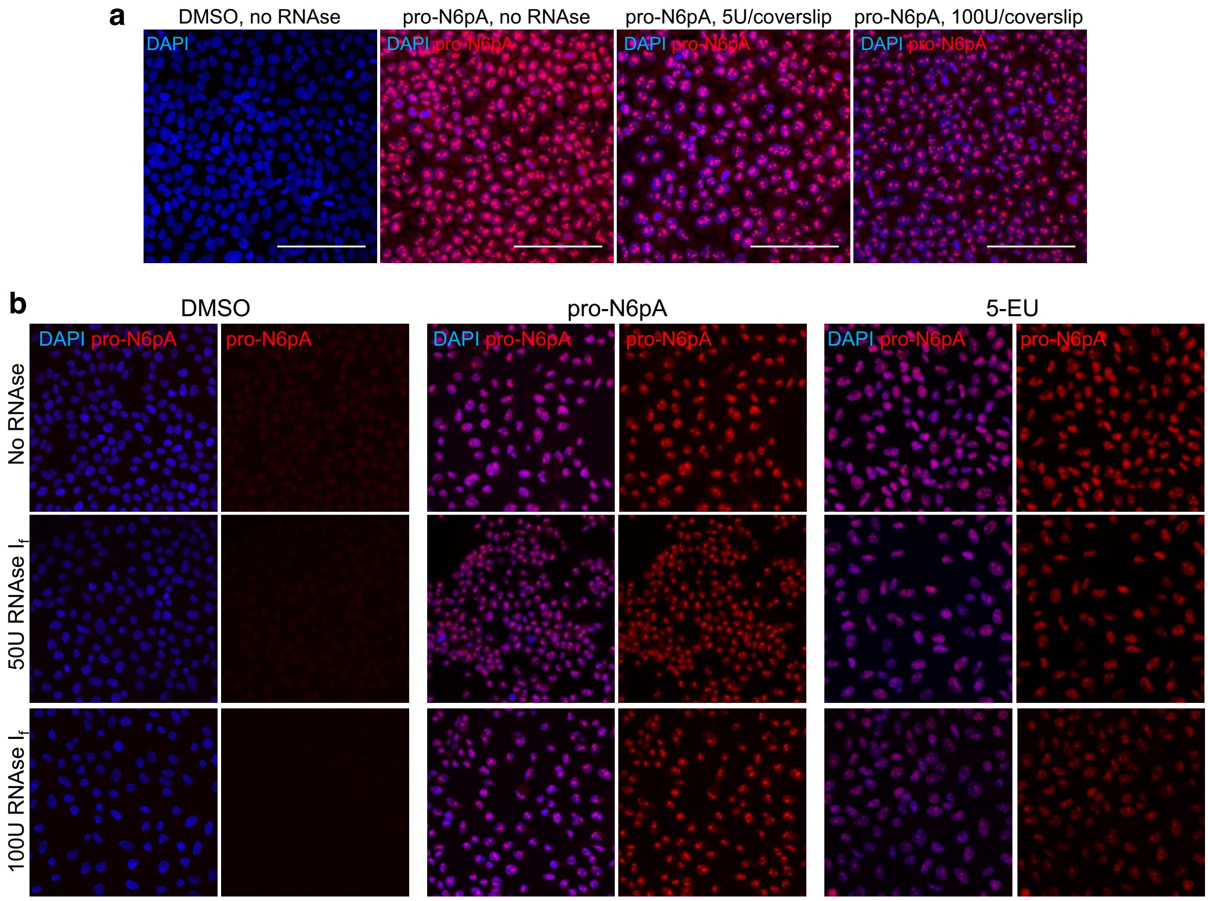


**Figure S16 | pro-N6pA is specifically incorporated into proteins.** RNAse treatment of cells cultured with pro-N6pA or 5-EU demonstrates that pro-N6pA is only to a small extent incorporated into RNAs. **a,** HeLa cells were grown on coverslips in 24-well plates and incubated with 100 µM pro-N6pA for 16 h before PFA-fixation. Prior to addition of rhodamine-azide by click reaction, they were treated with different amounts of RNAse $I_f$ (5U or 100U of RNAse per one well). The dotty pattern of localisation of the probe seems to be specific to the proteins it was incorporated in (Scalebar = 100 µm). **b**, 5-EU signal strongly vanishes with RNAse $I_f$ treatment, whereas pro-N6pA staining does not.

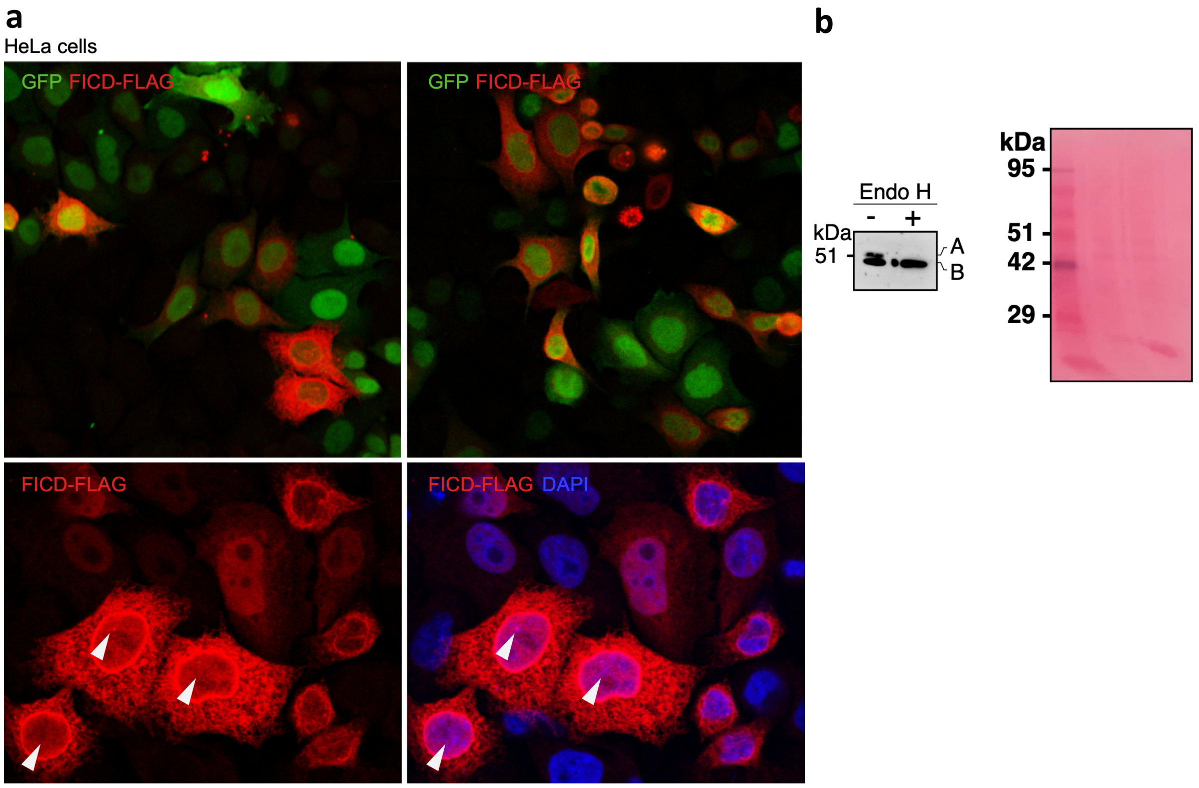


**Figure S17 | FICD localises in nucleus and cytoplasm in HeLa cells. a**, Overexpression of FICD with C-terminal 1 x FLAG-tag (red) in HeLa cells shows FICD localised to both nucleus and cytoplasm. GFP (green) was used as transfection control and DAPI (blue) for nuclear counterstaining. White arrows point towards nuclear FICD localisation. **b**, Endo H assay showing the presence of un-glycosylated FICD in HeLa cell lysate. A: glycosylated FICD and B: un-glycosylated FICD . The whole blot is shown stained with the Ponceau S. The experiment was performed in duplicate. Partial glycosylation of FICD was recently reported in ref 49.

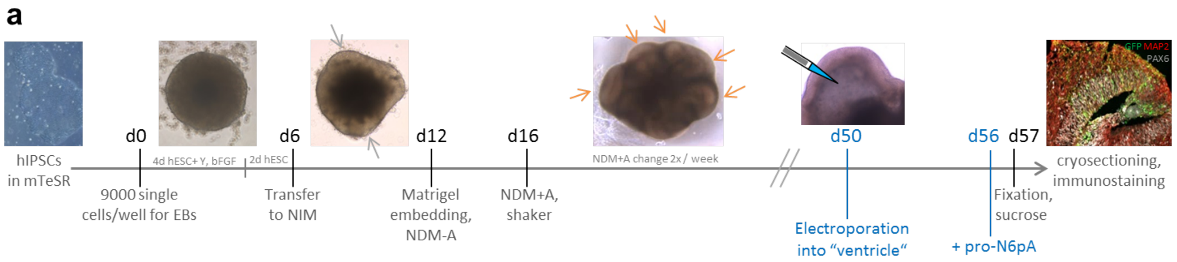

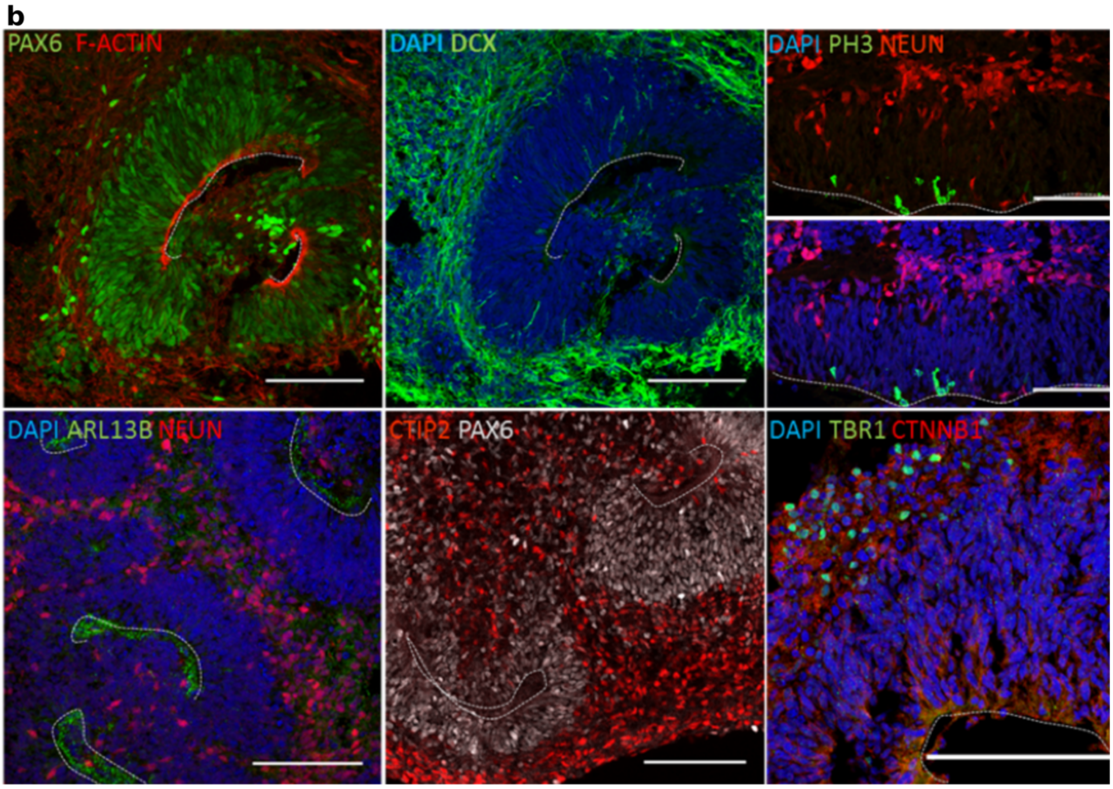


**Figure S18 | Cerebral organoids preparation and characterization**. **a,** Timeline for generation of cerebral organoids and their treatment with pro-N6pA (according to ref. 39), early neuroectoderm: grey arrows; expanded neuroepithelium: orange arrows; (EBs: embryoid bodies; hESC: low bFGF human embryonic stem cell medium; NIM: neural induction medium; Y: Rock inhibitor Y-27632; bFGF: basis fibroblast growth factor FGF-2; NDM: neural differentiation medium; +/- A: B27 supplement with or without vitamin A). **b,** Generation of cerebral organoids containing germinal zones comprised of different cell types that arise during dorsal cortical development. The progenitor zone, consisting of proliferative (e.g. PH3$^+$ for M-phase) PAX6$^+$ radial glia is apically delineated by ARL13B$^+$ primary cilia of radial glia, pointing into the ventricle-like lumen, and by apical junctions containing F-ACTIN and CTNNB1. Radial glia give rise to neuroblasts (DCX$^+$) that migrate basally, where they differentiate fully to become NEUN$^+$ neurons. The neocortical layers form in an inside-out manner with TBR1$^+$ and CTIP2$^+$ deep-layer projection neurons being generated first (Immunostainings of cerebral organoids cultured for 55-60 days, apical surface indicated by dotted line, Scalebar = 100 µm).

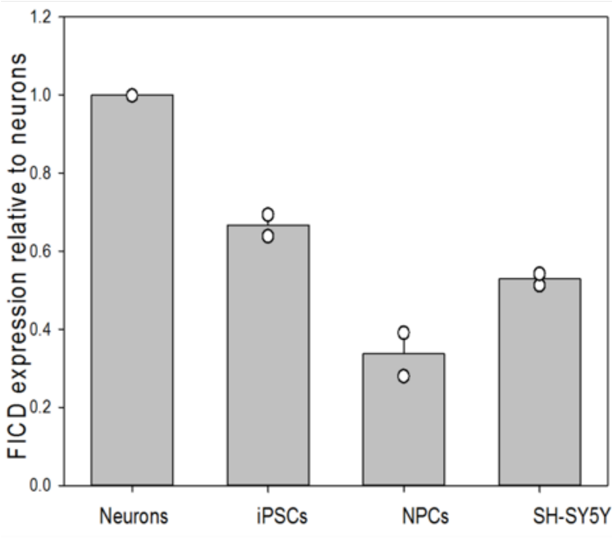


**Figure S19 | Neurons exhibit higher expression levels of FICD.** qPCR based analysis of FICD expression in different cell types confirms highest level in neurons. Error bars represent standard deviation.

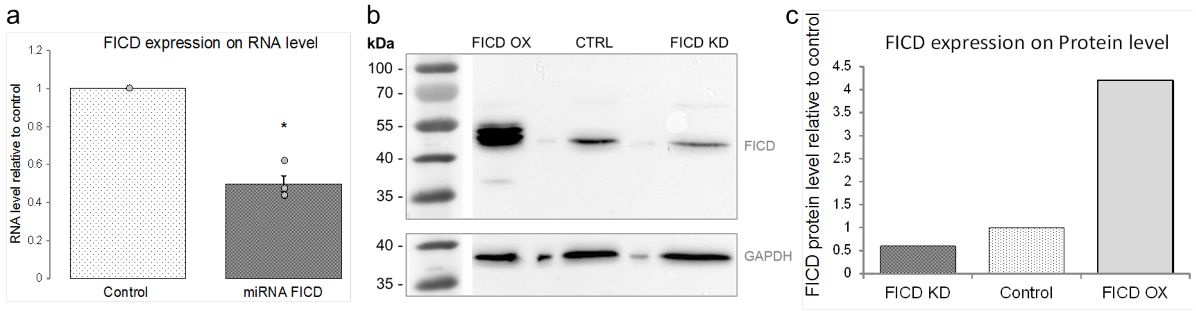


**Figure S20 | Validation of miRNAs targeting FICD.** miRNAs were generated following the BLOCK-iT™ Pol II miR RNAi Expression Vector Kits protocol by Invitrogen. Validation in HeLa via qPCR (**a**) and Westernblot (**b, c**) showed a knockdown efficiency to 50% of the control expression level both for RNA and protein (**a**: *$P=0.029$).

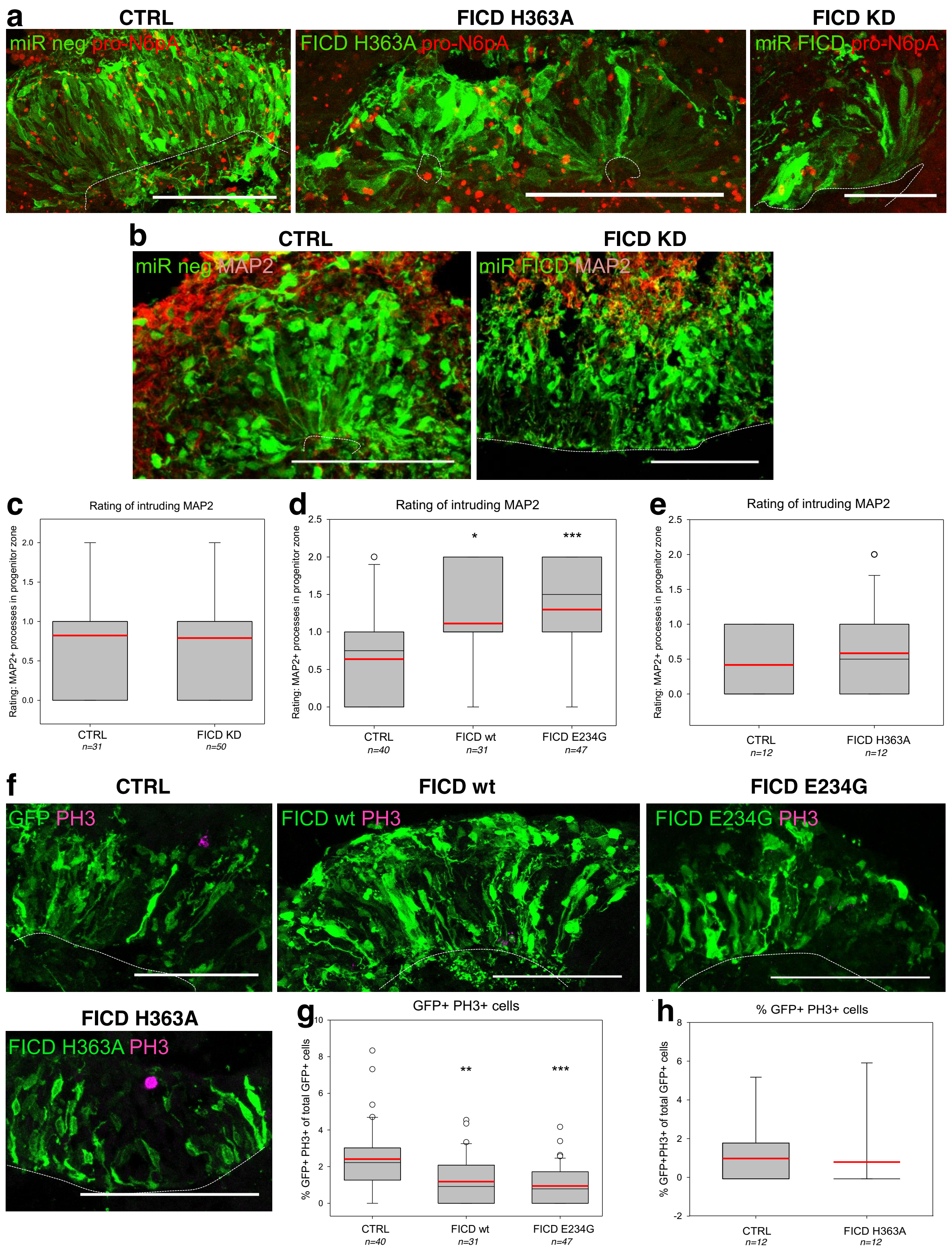


**Figure S21 | pro-N6pA signal and MAP2+ processes in COs upon FICD KD, rating of MAP2 intrusion, and mitotic cells upon FICD OX. a,** Click chemistry (red) in electroporated COs shows no difference in probe distribution in the case of catalytically inactive FICD H363A, but a decrease in signal upon FICD KD (transfected cells shown in green). **b,** IHC shows that MAP2+ neuronal processes (red) do not intrude the progenitor zone in COs upon FICD KD. **c, d, e,**

Rating of intrusion of MAP2+ neuronal processes into the progenitor zone of COs electroporated with constructs for FICD KD (**c**), FICD wt or E234G OX (**d**) or FICD H363A (**e**) shows increase upon FICD wt and E234G OX, but no change for KD and inactive FICD H363A. (Looking at IHC staining for MAP2, each ventricle was rated concerning the amount of neuronal processes inside the progenitor zone as *0 = none, 1 = few, 2 = many*.) **f,** Example images of IHC for PH3+ mitotic cells (magenta) upon FICD wt/E234G/H363A OX in COs. **g, h,** Quantification of PH3+ electroporated (GFP+) cells in germinal zones of COs eletroporated with FICD wt or E234G (**g**) or H363A (**h**) shows significant decrease when wt or activated FICD is overexpressed, but no change for H363A.

Electroporated organoids were analysed at 50d+7. **a, b, f,** electroporated cells and their progeny are shown in green; Scalebar = 50 µm; dotted line = apical surface. **c, d, e, g, h,** 1n = 1 electroporated germinal zone; box plot: mean (red line), median (black line), box represents 25th and 75th percentiles, whiskers extend to 10th and 90th percentiles, all outliers are shown; significance was tested using Kruskal-Wallis One-way ANOVA on Ranks and Dunn's Pairwise Multiple Comparison (**P<0.05*; ***P<0.01* ; ****P<0.001*).

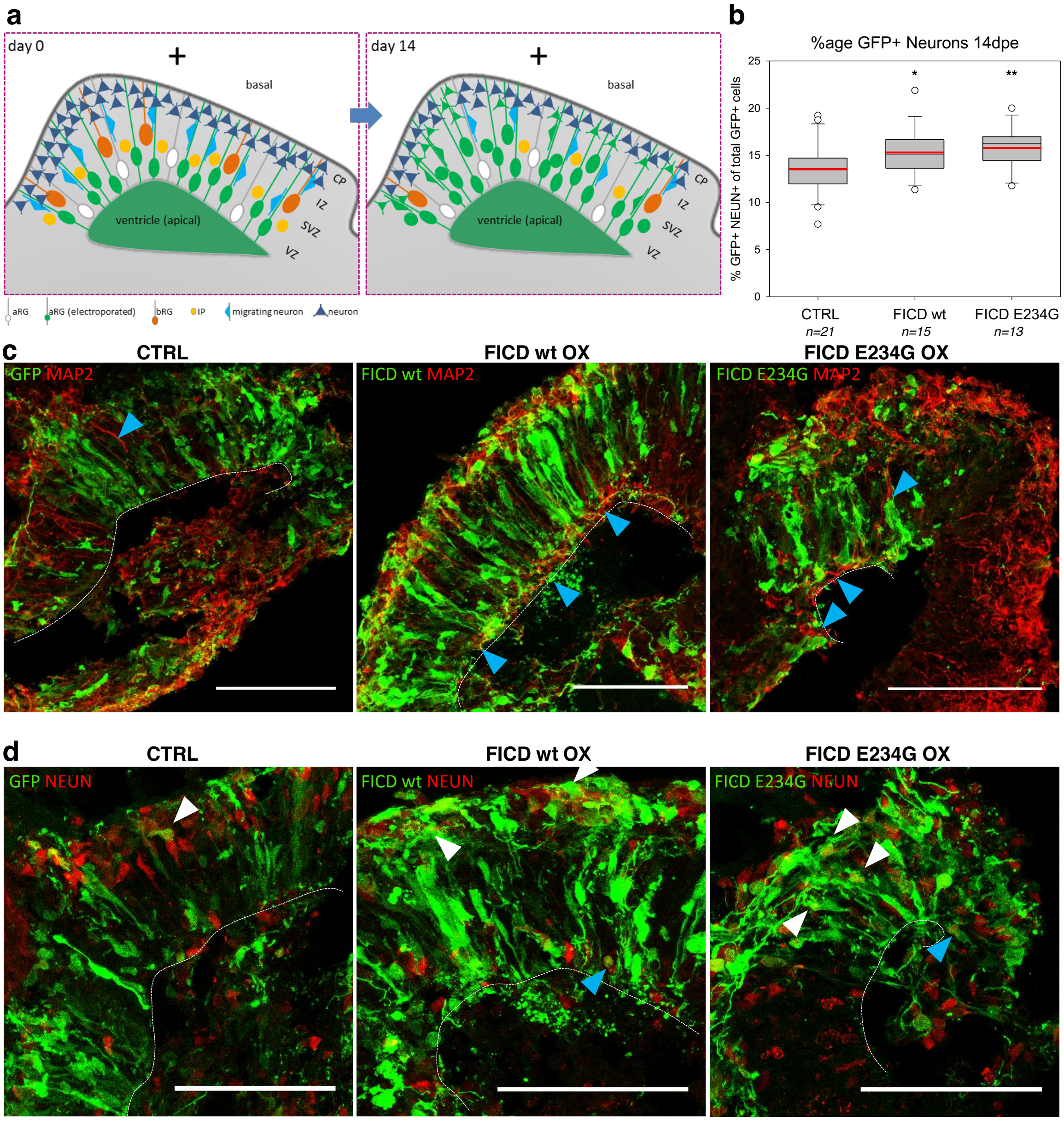


**Figure S22 | Overexpression of FICD in radial glia accelerates neurogenesis. a**, Scheme showing electroporated cells in CO germinal zone at day 0 and day 14: Over 14 days culture after electroporation, transfected (green) radial glia cells give rise to more differentiated cell types (green IPs, immature migrating neurons and differentiated neurons on day 14). **b**, Quantification of GFP$^+$ neurons 14 dpe shows natural increase in GFP$^+$ neurons over time (higher percentage than 7 dpe) and a significant increase in neuronal differentiation upon overexpression of FICD wt or E234G compared to control. **c,** Immunohistochemical (IHC) staining for electroporated cells (green) and neuronal dendrites (MAP2, red) shows neuronal processes in the progenitor zone. **d,** Immunostaining of electroporated cells (GFP, green) and nuclei of differentiated neurons (NEUN, red)

**a, e, f,** Scalebar = 50 µm, dotted line = apical surface; **b, d,** Box plots: mean (red line), median (black line); box represents 25th and 75th percentiles, whiskers extend to 10th and 90th percentiles, all outliers are shown; significance was tested using Kruskal-Wallis One-way ANOVA on Ranks and Dunn's Pairwise Multiple Comparison ($* \mathrm{P} < 0.05$; $** \mathrm{P} < 0.01$ ; $***P < 0.001$).

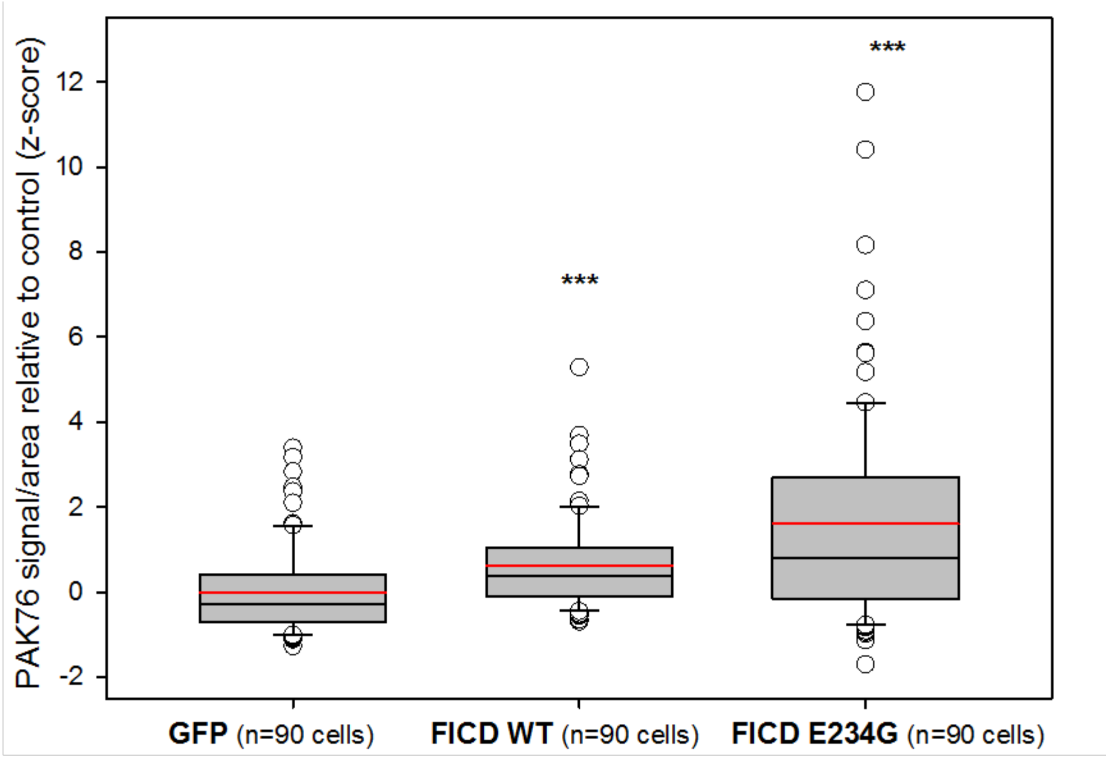


**Figure S23 | pro-N6pA signal upon FICD overexpression in SH-SY5Y cells.** Transfection of SH-SY5Y cells with overexpression construct vs. GFP (3 µg DNA per 1 Mio. cells), plating on 3 separate coverslips per condition. Pro-N6pA treatment for the last 16 h of culture. Fixation 48h post-transfection, followed by click-chemistry and nuclear counterstaining with DAPI. AMPylation signal was measured using Fiji, selecting transfected GFP+ cells and normalising to their area. Mean (red line), median (black line), box represents 25th and 75th percentiles, whiskers extend to 10th and 90th percentiles, all outliers are shown. One-tailed Student's *t*-test. $***P < 0.001$.

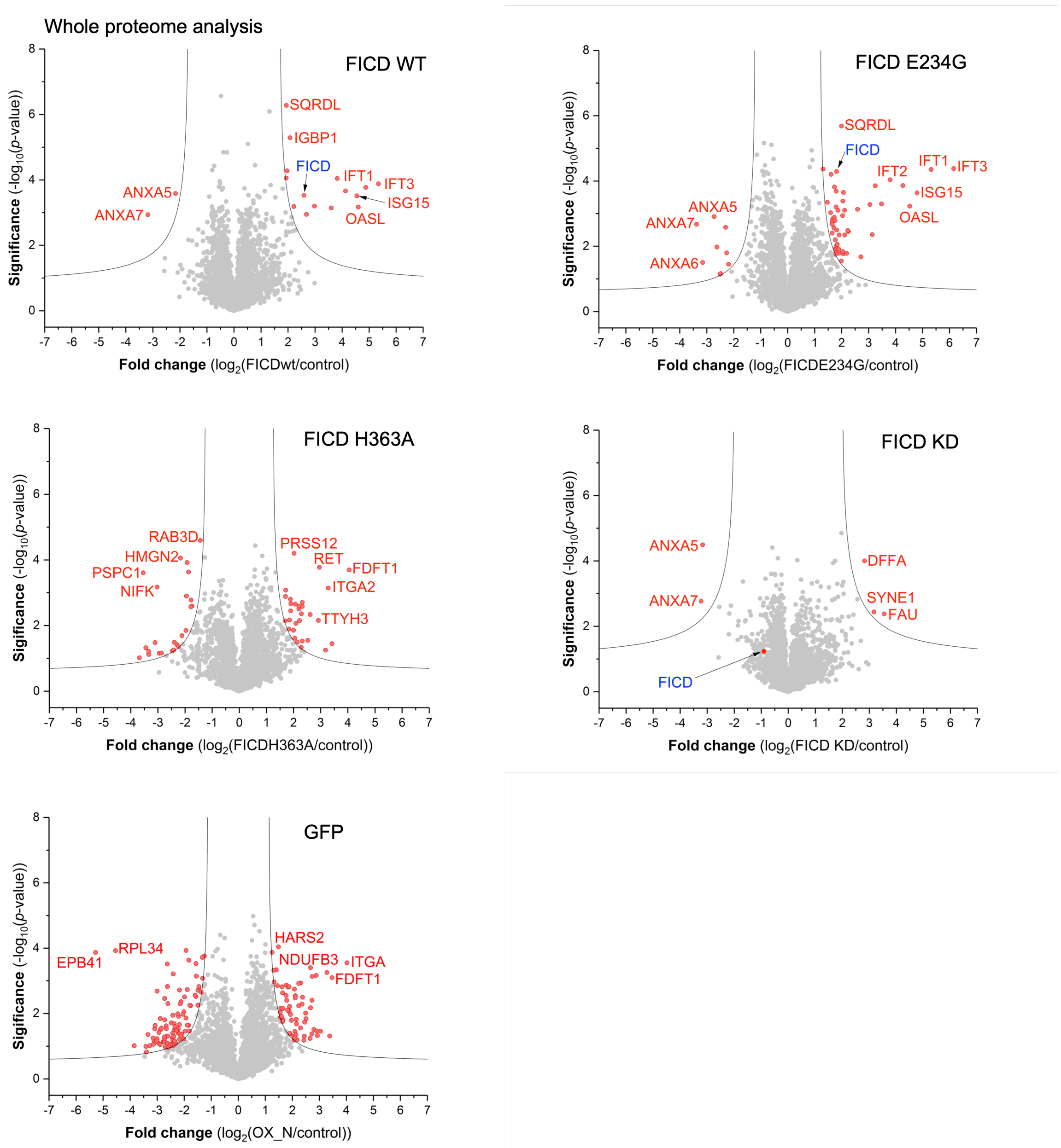


**Figure S24 | Whole proteome analysis of FICD OX and KD in SH-SY5Y cells.** In red are highlighted proteins with significant change. The proteins above the threshold are involved in innate immune response to viral infection, may be due to the exogenous DNA/plasmid transfection (IFT1, ISG15, OASL, ANXA7, ANXA5, PSPC1, DFFA). Cut-off line is at FDR 0.05; s0 2; *n* = 3.

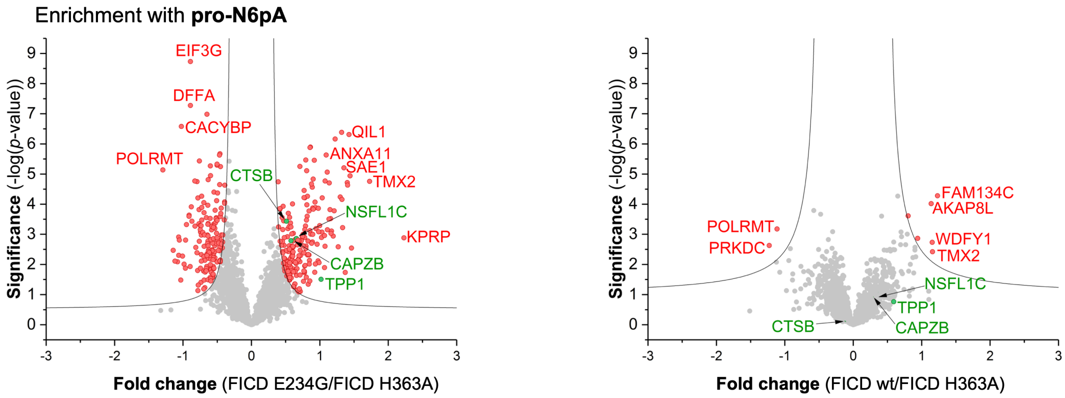


**Figure S25 | pro-N6pA pull-downs upon FICD overexpressions in SH-SY5Y cells.** Transfection of SH-SY5Y cells with overexpression construct (2 µg DNA per 1 Mio. cells). Volcano plots visualise the difference in AMPylation between FICD (E234G and wt) and FICD H363A OX. The x-axis is in $\log_2$ scale, (FDR 0.05; s0 0.3; $n$ = 7). Red circles represent all significantly enriched proteins. Green circles represent proteins found AMPylated in COs.

**Cytotoxicity.** Cytotoxicity of pro-N6pA and N6pA was measured by MTT assay. The $IC_{50}$ of pro-N6pA is 300 µM and $IC_{50}$ of N6pA is above 1 mM. pro-N6pA low cytotoxicity makes it suitable for the use in high concentrations for cell treatment (**Fig. S1**).

**Immunoprecipitation of PFKP and PPME1.** In order to further corroborate the efficiency of incorporation of pro-N6pA probe as a protein PTM, the PFKP and PPME1 proteins identified in HeLa cells were immunoprecipitated from pro-N6pA treated HeLa cell lysates and after the click-chemistry with rhodamine-azide and SDS-PAGE, in-gel fluorescence was scanned showing strong labelling at 85 kDa (PFKP) and 42 kDa (PPME1) while no signal was observed in control sample (without antibody) (**Fig. S3**).

**Cell type dependent changes in AMPylation during endoplasmic reticulum stress.** Proteome profiling under endoplasmic reticulum (ER) stress was performed to determine whether modifications are altered as a result of HSPA5 upregulation, as previously reported for thapsigargin (Tg)-treated HEK293T cells[4]. Only a slight increase in absolute AMPylated HSPA5 was observed in HeLa and A549 cells. Despite the moderate impact of ER stress on AMPylation in these cells, we found 145 dysregulated proteins in SH-SY5Y (**Fig. S7** and **S12**). This further corroborated the role of AMPylation in neuronal-like cells. Further, Tg induced ER stress in iPSCs, NPCs and neurons results in very distinguished changes in AMPylation (**Fig. S11**), with strong response in iPSCs and only two proteins were influenced in neurons.

**Site identifications on cathepsins in SH-SY5Y cells and cerebral organoids.** In addition to the AMPylated serines identified on cathepsins in HeLa cells (**Fig. 2** and **Fig. S4**) we confirmed the AMPylation on CTSB (S104 and S107) and CTSL (S137) also in SH-SY5Y cells (**Fig. S5**) and found two new sites on CTSV (S137) and CTSS (S135) in COs, the CTSB (S107) and CTSC (S254) AMPylation sites were confirmed (**Fig. S5**). See **Table S4** for complete characterization. We used similarly to previous site identifications approach employing the TEV-cleavable linker (**Fig. S4**).

**HSPA5 is AMPylated on T518 in U2OS cells.** In addition to the identified AMPylation sites on cathepsins using biotin-TEV-cleavable linker we have searched by MaxQuant MS spectra measured in ref. 50. These were acquired to identify the protein ADP-ribosylation sites in bone osteosarcoma epithelial cells (U2OS) on Orbitrap Fusion using a combination of higher-energy collision-induced dissociation (HCD) and electron transfer dissociation (ETD) acquisition. The ETD ion source was triggered by adenine diagnostic peak (136.062 Da). Since both ADP-ribosylated and AMPylated peptides contain the adenine base this setup inherently leads to the raw MS spectres which include also an intact AMPylated peptides. The MaxQuant search for AMPylated peptides (+329.052 Da) revealed HSPA5 AMPylation on T518. This corroborates the findings by *Preissler et al.* (ref. 4 and 5).

**ADP-ribosylation exclusion.** MS raw files obtained from site identification experiments using the TEV-cleavable linker were searched for potential ADP-ribosylation (+906.2786) on the amino acids EDSTR. However, none of the identified ADP-ribosylation sites was included as a hit in the chemical-proteomic pull-down experiments. A reference list of ADP-ribosylated proteins was adopted from ref. 27 and can be found in **Table S3**. To exclude experimentally incorporation of the pro-N6pA probe into ADP-ribosylation we have pre-treated HeLa cells with poly(ADP-ribose)polymerases (PARP) inhibitors 4-aminobenzamide (4-ABA) or olaparib (Ola) before the pro-N6pA labelling, similarly reported in ref. 25. For both PARP inhibitors we have not observed any influence on labelling intensity or change in labelling pattern thus concluding that majority of the labelling yield from the AMPylation (**Fig. S3**).

**Protein interaction and GO term analysis**. In order to understand if AMPylation plays a role in the regulation of specific biological processes or pathways, network and GO term analysis was performed using STRING.db[51]. Resulting networks for AMPylation targets identified in cancer cell lines and iPSCs have significantly more interactions than expected (cancer cell lines: PPI enrichment *p*-value *P* = 0.00028; iPSCs *P* = $1.7 \cdot 10^{-5}$ and are significantly enriched for proteins with metabolic function (**Table S1**). In NPCs, there is only an enrichment for a single KEGG pathway ("biosynthesis of amino acids") and the number of targets is very low. Their differentiation to neurons results in a state with a high number of AMPylated proteins again, which are different from those in pluripotent stem cells and NPCs and build a strongly interacting network (PPI enrichment *p*-value *P* = 0.000124). This network is, in addition to a new set of proteins involved in metabolic processes, significantly enriched for proteins with cytoskeletal and motor function (**Table S13**). Interestingly, these proteins are known to be important for neuronal development and function and have been shown to be modified by multiple other PTMs. Looking at AMPylated targets in iPSCs, NPCs and neurons, it becomes clear that during neuronal differentiation, cellular proteins undergo a process of complete remodelling of AMPylation that resembles an hourglass (**Fig. 4b** and **Fig. S13**): of many targets in iPSCS hardly any are left in NPCs and differentiation to mature neurons goes along with AMPylation of a big and unique group of new target proteins. As expected, additional targets were found in the more heterogeneous and complex COs. Thus, AMPylation of proteins known to play a role in neuronal development and function could represent one additional level of fine tuning and regulating cortical development.

**(B) Structures and synthesis**

**General methods.** Reagents and solvents were purchased from commercial suppliers (*Sigma-Aldrich Co. LLC*, *Thermo Fisher Scientific Inc.*, *Merck KGaA*, *TCI Europe GmbH*, *Fluorochem Ltd.* and *Alfa Aesar GmbH*) and used without further purification. HPLC-grade solvents or anhydrous solvents (max. 0.01 % water content, stored over molecular sieve under an argon atmosphere, *Sigma-Aldrich Co. LLC*) were used for all reactions. All experiments were monitored by analytical thin layer chromatography (TLC). TLC was performed on pre-coated silica gel plates (60 F-254, 0.25 mm, *Merck KGaA*) with detection by UV (λ = 254 and/or 366 nm) and/or by colouration using a potassium permanganate ($KMnO_4$) stain and subsequent heat treatment. Flash chromatography was performed on silica gel 60 (0.035 – 0.070 mm, mesh 60 Å, *Merck KGaA*) with the indicated eluent. $^{1}H$, proton-decoupled $^{13}C$ and proton-decoupled $^{31}P$ NMR spectra were recorded on a *Bruker* Avance III HD 300 (300 MHz), a *Bruker* Avance I 360 (360 MHz) or a *Bruker* Avance III HD (500 MHz) at 298 K. Chemical shifts are reported in delta (δ) units in parts per million (ppm) relative to distinguished solvent signals. The following abbreviations are used for the assignment of the signals: s – singlet, d – doublet, t – triplet, q – quartet, m – multiplet. Coupling constants *J* are given in Hertz [Hz]. HR-MS spectra were recorded in the ESI mode on a Thermo Scientific LTQ-FT Ultra (FT-ICR-MS) coupled with an UltiMate 3000 HPLC system (*Thermo Fisher Scientific Inc.*).

**4-(Prop-2-yn-1-ylamino)-7-(2',3'-*O*-Isopropylidene-β-D-ribofuranosyl)-1*H*-purine.** After suspending $N^6$-propargyl-adenosine (106 mg, 0.506 mmol) in acetone (13.0 mL), 2,2-dimethoxypropane (0.620 mL, 5.06 mmol) was added. Then *p*-toluenesulfonic acid monohydrate (1.61 g, 8.47 mmol) was added and the mixture was stirred for two hours at room temperature. The reaction was stopped with aqueous $NaHCO_3$ (24 mL), extracted with EtOAc (3×30 mL) and the combined organic phases were dried over $Na_2SO_4$ and filtered. The solvents were removed under reduced pressure and the residue was purified by column chromatography on silica gel (2% → 5% MeOH in DCM) to give the product as a pale yellow solid (149 mg, 85 %). Spectral analysis was in good accordance with the literature[52].

**Phenyl(benzyloxy-L-alaninyl)phosphorochloridate[53].** Anhydrous triethylamine (264 µL, 1.896 mmol) was added dropwise to a stirred solution of the phenyl dichlorophosphate (142 µL, 0.948 mmol) and the L-alanine benzyl ester *p*-toluenesulfonate salt (333 mg, 0.948 mmol) in anhydrous DCM (4 mL) at -78 °C. Following the addition, the reaction mixture was stirred at room temperature for 1 h. After this period the solvent was removed under reduced pressure and the residue triturated with anhydrous THF. The precipitate was filtered and the solution was concentrated to give a colourless oil. The crude product was used for the next step.

**4-(Prop-2-yn-1-ylamino)-7-(2′,3′-*O*-isopropylidene-β-D-ribofuranosyl)-1*H*-purine 5′-*O*-[phenyl(benzyloxy-L-alaninyl)]phosphate.** Protected $N^6$-propargyl-adenosine (89 mg, 0.26 mmol) was dissolved in THF (3 ml). Solution of *t*BuMgCl in THF (1M, 0.52 ml, 0.52 mmol) was added drop-wise and the reaction mixture was stirred for 15 minutes. Then solution of a phenyl(benzyloxy-L-alaninyl)phosphorochloridate (228 mg, 0.64 mmol) in THF (3 ml) was added and the reaction mixture was stirred overnight at room temperature. Reaction was quenched by saturated $NH_4Cl$ solution (1 ml) and mixture was extracted with EtOAc and water. Organic layer was dried over $Na_2SO_4$, filtered and evaporated under reduced pressure. Crude product was purified using column chromatography on silica gel (10% HEX in EtOAc) protected phosphoramidate (100 mg, 63 %) was obtained as colourless oil (mixture of diastereoisomers ~1:1). $^1$H NMR (300 MHz, $CDCl_3$) δ 8.43 (s, 1H), 8.41 (s, 1H), 7.94 (s, 1H), 7.91 (s, 1H), 7.37 – 6.97 (m, 20H), 6.38 (bs, 2H), 6.09 (d, *J* = 5.3 Hz, 1H), 6.08 (d, *J* = 5.3 Hz, 1H), 5.37 (dd, *J* = 6.3, 2.3 Hz, 1H), 5.16 – 4.99 (m, 4H), 4.92 (dd, *J* = 6.3, 2.6 Hz, 1H), 4.53 – 4.38 (m, 6H), 4.37 – 4.19 (m, 4H), 4.09 – 3.91 (m, 4H), 2.35 – 2.17 (m, 2H), 1.62 – 1.57 (m, 6H), 1.41 – 1.16 (m, 12H). $^{13}$C NMR (75 MHz, $CDCl_3$) δ 153.21, 139.71, 139.45, 129.74, 128.74, 128.58, 128.31, 128.25, 125.09, 125.03, 120.27, 120.21, 120.15, 120.08, 114.50, 91.62, 91.15, 85.70, 85.60, 84.37, 84.30, 81.57, 81.41, 80.15, 77.58, 77.36, 77.16, 76.74, 71.71, 71.65, 67.32, 66.48, 66.41, 50.51, 50.33, 27.26, 27.17, 25.47, 25.30, 21.08, 20.90. $^{31}$P NMR (121 MHz, $CDCl_3$) δ 2.56, 2.45. MS (ESI$^+$): *m/z* (*%*): 663.18 (100). [M + H]$^+$. HRMS (ESI$^+$): *m/z* [M + H]$^+$ calculated for $C_{32}H_{36}N_6O_8P^+$: 663.23323; found 663.23219.

**4-(Prop-2-yn-1-ylamino)-7-(β-D-ribofuranosyl)-1*H*-purine 5′-*O*-[phenyl(benzyloxy-L-alaninyl)]phosphate.** Protected phosphoramidate (100 mg, 0.15 mmol) was dissolved in 90% aqueous TFA (2 ml) and stirred for 2 h at room temperature. Then the solvent was repeatedly co-evaporated with methanol under reduced pressure. Product was purified using column chromatography on silica gel (10% MeOH in DCM). Several co-evaporations with chloroform gave phosphoramidate PRO-N6PA (61 mg, 65 %) as a white solid (mixture of diastereoisomers ~1:1). $^1$H NMR (500 MHz, MeOD) δ 8.26 (s, 2H), 8.20 (s, 1H), 8.17 (s, 1H), 7.32 – 7.20 (m, 14H), 7.19 – 7.10 (m, 6H), 6.00 (m, 2H), 5.11 – 4.99 (m, 4H), 4.63 – 4.57 (m, 2H), 4.40 – 4.31 (m, 8H), 4.31 – 4.23 (m, 2H), 4.22 – 4.18 (m, 2H), 2.58 (t, *J* = 2.5 Hz, 2H), 1.33 – 1.19 (m, 6H). $^{13}$C NMR (126 MHz, MeOD) δ 173.42, 173.39, 173.22, 173.18, 152.47, 152.43, 150.66, 150.61, 139.46, 139.41, 135.82, 135.74, 129.35, 128.14, 127.87, 127.84, 124.78, 120.03, 119.99, 119.93,

119.63, 88.52, 82.95, 82.91, 82.88, 82.84, 79.75, 73.96, 70.86, 70.14, 70.02, 66.53, 66.51, 66.12, 66.08, 65.69, 65.65, 50.26, 50.14, 19.01, 18.96, 18.85, 18.79. $^{31}$P NMR (121 MHz, MeOD) δ 3.89, 3.64. MS (ESI$^+$): *m/z* (*%*): 623.25 (100) [M + H]$^+$. HRMS (ESI$^+$): *m/z* [M + H]$^+$ calculated for $C_{29}H_{32}N_6O_8P^+$: 623.20193; found 623.20224.

**(C) Biological methods**

**Metabolomics samples preparation.** Samples were prepared according to protocol from Matano *et al.* with minor changes[54]. Cells were seeded in 60 mm cell culture dishes (approx. 800,000 cells/dish) till 90% confluency. Media was removed, cells washed with 3 mL PBS and incubated with 3 mL media, supplemented with **pro-N6pA** in DMSO (100 µM final conc.) or DMSO respectively. After the particular incubation times, cells were washed twice with 2 mL warm PBS and twice with 2 mL water (LC-MS grade, *Fisher Chemical*). Subsequently 4 mL of cold quenching solution (ACN:MeOH:$H_2O$ 2:2:1 incl. 0.5M FA, -20°C) was added, cells detached by scraping and transferred to falcons. Washing and quenching of cells was completed in 1-2 min. Cell lysis was carried out by sonication (30s, Sonorex RK 100, *Badelin*) while cooling followed by incubation on ice (15 min) prior to snap-freezing in liquid nitrogen. Samples were freeze-dried until complete dryness and stored at -80°C until day of measurement. For MS-analysis dried samples were dissolved in 20 µL water (LC-MS grade, *Fisher Chemical*), centrifuged (21000 x *g*, 10 min, 4°C) and transferred into LC-MS vials. A pooled sample was prepared from all samples for column equilibration

**Metabolomics sample measurement and evaluation.** Metabolic profiling and MS/MS analysis was carried out on an Ultimate 3000 RSLC system (*Thermo Scientific*) coupled to an LTQ Orbitrap XL mass spectrometer (*Thermo Scientific*). Chromatographic separation was performed using a SeQuant® ZIC®-pHILIC (250 x 2.1 mm, 5 µm, 100 Å) (Merck Millipore) at 40 °C. Gradient elution was carried out with 5 mM $NH_4OAc$ (LC-MS grade, *Fisher Chemical*) in 100% $H_2O$ (LC-MS grade, *Fisher Chemical*) (A) and 5 mM $NH_4OAc$ in 95% ACN (LC-MS grade, *Fisher Chemical*) (B). After 5 min pre-equilibration with 98% B, samples were eluted with a linear gradient from 98% to 53% B over 20 min at a flow rate of 250 µl/min followed by a 9 min washing step with 5% B and re-equilibration with 98% B for 4 min. Mass spectrometric measurements were accomplished in HESI- mode (H-ESI-II probe, Thermo Scientific) with following source parameters: capillary voltage 4.5 kV, tube lens -110 V, vaporizer temperature 80 °C, sheath gas 30 l/h, aux gas 10 l/h, capillary voltage -60 V and capillary temperature 275 °C. Parent ions of interest (**N6pATP** *m/z* = 544.00) was isolated (Isolation width 0.5 *m/z*) at 60,000 resolution in the orbitrap in profile mode and subjected to collision-induced dissociation (normalized collision energy 45 V) in the SRM mode and most intense daughter ion (**N6pATP** *m/z* = 446.03) isolated (SIM width 5 *m/z*) at a resolution of 7,500 in the orbitrap and used for quantification. Retention time, exact mass and MS/MS spectra of quantified compound were matched to an authentic synthetic standard. Prior to measurement, 15 pooled quality control (QC) samples were injected to equilibrate the column. In order to take into account metabolic degradation over time, the sample order was randomized. Data were processed with Xcalibur 2.2 SP1.48 Quan Browser (*Thermo Scientific*) using the Genesis algorithm for peak detection and manual integration mode. Peak detection was carried out with the following track peak parameters: Genesis Peak Detection: Highest Peak, Minimum peak height (S/N) = 3.0. Intensities of the corresponding daughter ion was extracted and plotted for each time point. (Experiments were carried out in two biological replicates).

**Cytotoxicity assay.** Human epitheloid cervix carcinoma cells (HeLa) cultivated in high glucose Dulbeccos´s Modified Eagle´s Medium supplemented with 10% (v/v) fetal bovine serum and 2 mM L-glutamine were seeded at a density of 5,000 cells per well (100 µL of a solution of 50,000 cells/mL) in a transparent, flat-bottom 96-well plate. Cells were grown over night in a humidified atmosphere at 37 °C and 5% $CO_2$. The next day the medium was removed and replaced by fresh medium supplemented with 1000, 700, 500, 300, 200, 100 or 30 µM pro-N6pA or N6pA or 1% (v/v) DMSO as a control. The cells were incubated for 24 h. To determine metabolic activity of the cells, 20 µL of 3-(4,5-dimethyl-2-thiazolyl)-2,5-diphenyl-2*H*-tetrazolium bromide solution (MTT, 5 mg/mL in PBS) were added to each well. The cells were incubated for 4 h. The medium was completely removed and the formazan crystals were resuspended in 200 µL DMSO and absorbance at 570 nm and at a reference wavelength of 630 nm was determined in an infinite F200 pro plate reader (*Tecan*). All data points were measured in triplicate. The data was normalized with respect to the DMSO control.

**Protein concentration.** Protein concentrations were determined by bicinchoniic acid assay (BCA, *Carl Roth GmbH + Co.*).

**Probe treatment, click-chemistry and enrichment.** Cells (6-cm dishes) were treated with the probes at 80-90% confluency. Culture medium was removed and the cells or COs were labelled in fresh media containing 100µM pro-N6pA or 100µM N6pA (both stocks 100mM in DMSO) for 16 h at 37 °C in cells incubator. Subsequent cell lysis was done in 1 mL ice cold lysis buffer (1% (v/v) NP40, 1% (w/v) sodium deoxycholate and 1 tablet protease inhibitor (cOmplete™, Mini, EDTA-free protease inhibitor cocktail, *Roche*) in 10 mL PBS) while mechanically detached by scraping and subsequently transferred into an eppendorf tube or by ultrasonication at 40 % intensity for 10 s. Lysis was done for 30 min at 4 °C while rotating. Insoluble fraction was pelletized (10 min, 14,000 g, 4 °C) and protein concentration was determined by BCA. To 500 µg (HeLa, A549, SH-SY5Y cells, COs and fibroblasts) or 250 µg (iPSCs, NPCs, neurons) of a protein lysate in total volume of 970 µL 0.2% (w/v) SDS in PBS 10 µL 5-rhodmamine-azide (10 mM in DMSO, *Jena Bioscience*; in-gel analysis) or 10 µL azide-$PEG_3$-biotin (10 mM in DMSO, *Jena Bioscience*; for MS samples), 10 µL Tris(2-carboxyethyl)phosphine (TCEP) (53 mM in $ddH_2O$) and 1.2 µL tris(benzyltriazolylmethyl)amine (TBTA) (83.5 mM in DMSO) were added. Samples were gently vortexed and the click reaction was initiated by the addition of 20 µL $CuSO_4$ solution (50 mM in $ddH_2O$). The mixture was incubated at 25 °C for 1.5 h. Afterwards proteins were precipitated by addition of 4 mL acetone and incubation overnight at -20 °C. The protein pellet was harvested by centrifugation at 9,000 g for 15 min at 4 °C and washed twice with 1 mL of ice cold methanol. Proteins were reconstituted in 500 µL 0.2% (w/v) SDS in PBS, the remaining not soluble particles were spun down at 9,000 g for 5 min and resulting supernatant was loaded onto the beads and incubated at 25 °C for 1.5 h while rotating. The 50 µL of avidin-agarose beads (*Sigma-Aldrich*) were washed in advance thrice with 1 mL 0.2% (w/v) SDS in PBS. Afterwards through the enrichment protocol, the beads were always spun at 400 g, 25 °C, for 2 min. Beads were subsequently washed thrice with 1 mL 0.2% (w/v) SDS in PBS, twice with 1 mL 6 M urea in $ddH_2O$ and thrice with 1 mL PBS.

**Endoplasmic reticulum stress induction.** For induction of unfolded protein response, cells or COs were treated with 100µM pro-N6pA for 16 h, then the media was exchanged with fresh media containing 100µM pro-N6pA and 1 µg/mL thapsigargin (stock 10 mg/mL in DMSO) and incubated for 24 h.

**4-aminobenzamide (4-ABA) and Olaparib (Ola) poly(ADP-ribose) polymerases inhibition.** In HeLa (6-cm dishes) cells 2 h before addition of the pro-N6pA probe 4 µL of 10M 4-ABA or 2 µL 10mM Ola (both stock solutions in DMSO) were added to the 2 mL of the fresh culture media, after two hours 2 µL of 100mM pro-N6pA was added and incubated for 16 h. Subsequent cell lysis was done in 1 mL ice cold lysis buffer (1% (v/v) NP40, 1% (w/v) sodium deoxycholate and 1 tablet protease inhibitor (cOmplete™, Mini, EDTA-free protease inhibitor cocktail, *Roche*) in PBS) while mechanically detached by scraping and subsequently transferred into an eppendorf tube or by ultrasonication at 40 % intensity for 10 s. Lysis was done for 30 min at 4 °C while rotating. Insoluble fraction was pelletized (10 min, 14,000 g, 4 °C) and protein concentration was determined by BCA. To 200 µg of a protein lysate in total volume of 200 µL 0.2% (w/v) SDS in PBS 2 µL 5-rhodmamine-azide (10 mM in DMSO, *Jena Bioscience*; in-gel analysis) 2 µL Tris(2-carboxyethyl)phosphine (TCEP) (53 mM in $ddH_2O$) and 0.24 µL tris(benzyltriazolylmethyl)amine (TBTA) (83.5 mM in DMSO) were added. Samples were gently vortexed and the click reaction was initiated by the addition of 4 µL $CuSO_4$ solution (50 mM in $ddH_2O$). The mixture was incubated at 25 °C for 1.5 h. Afterwards proteins were precipitated by addition of 900 µl acetone and incubation 1 h at -20 °C. The protein pellet was harvested by centrifugation at 9,000 g for 15 min at 4 °C and washed once with 500 µl of ice cold methanol. Finally, it was dissolved in 50 µL of gel loading buffer (4% SDS, 20% glycerol, 10% 2-mercaptoethanol, 0.004% bromphenol blue in 125mM Tris-HCl, pH 6.8) and loaded on a 12.5% SDS-PAGE.

Chemical-proteomic samples preparation. Cells were treated with Olaparib and pro-N6pA probe in a same way as for in-gel analysis, but it was continued with click reaction with azide-$PEG_3$-biotin and continued with enrichment as described above.

**In-gel analysis.** Enriched rhodamine-tagged proteins were released from the beads with 50 µL of gel loading buffer (4% SDS, 20% glycerol, 10% 2-mercaptoethanol, 0.004% bromphenol blue in 125mM Tris-HCl, pH 6.8) and loaded on a 12.5% SDS-PAGE. After electrophoretic separation, fluorescently labelled proteins were visualized using a Fujifilm LAS-4000 equipped with a Fujifilm Fujinon VR43LMD3 lens and a 575DF20 filter (Fujifilm) operated in Cy3 fluorescence detection mode.

**Chemical-proteomics.** Enriched proteins were on beads digested in 200 µL digestion buffer (20 mM HEPES, pH 7.5, 7 M urea, 2 M thiourea). Proteins were reduced (0.2 µL 1 M DTT, 45 min, 25 °C) and alkylated (2 µL, 30 min, 25 °C, in the dark). The alkylation reaction was quenched by addition of 0.8 µL 1M DTT and incubation for 30 min at 25 °C. Proteins were pre-digested with 1 µL LysC (*Wako*) at 25 °C for 4 h. 600 µL 50 mM TEAB buffer was added and the proteins were digested overnight with 1.5 µL sequencing grade trypsin (0.5 mg/mL, Promega) at 37 °C. The following day the beads were settled and the supernatant was acidified with 10 µL formic acid to a pH of 2 – 3. Peptides were desalted and on-column dimethyl labelled using 50 mg SepPak C18 cartridges (*Waters Corp.*) on a vacuum manifold. The cartridges were equilibrated with 1 mL acetonitrile, 1 mL 80% acetonitrile and 3 mL 0.5% formic acid. The samples were loaded on the cartridges and subsequently washed with 5 mL 0.5% formic acid. Peptides were labelled with either 5 mL labelling buffer "light" (90 mM sodium phosphate, pH 7.5, 30 mM $NaBH_3CN$, 0.2% $CH_2O$) or 5 mL labelling buffer "heavy" (90 mM sodium phosphate, pH 7.5, 30 mM $NaBD_3CN$, 0.2% $^{13}CDO$). Cartridges were subsequently washed with 2 mL 0.5% formic acid. The peptides were eluted with two times 200 µL 80% acetonitrile, 0.5% formic acid. DMSO and probe-treated samples were combined and dried by lyophilization. Peptides were reconstituted in 30 µL 1% (v/v) formic acid, prepared for mass spectrometry by filtering through a membrane filter (Ultrafree-MC and –LC, Durapore PVDF-0.22 µm, *Merck Millipore*) and transferred into mass vials. For label-free quantification (LFQ) samples were prepared as above without the use of labelling solutions "light" and "heavy" and subsequent washing with 3 mL 0.5% formic acid. Samples were not combined. Experiments were conducted in 7 to 12 replicates. Label switch was used for dimethyl-labelling samples.

**Site identification using TEV-cleavable linker.** HeLa, SH-SY5Y or COs lysates in total volume of 2 mL 0.2% (w/v) SDS in PBS were supplemented with 20 µL azide-TEV-biotin (10 mM in DMSO, gift from B. F. Cravatt laboratory)[28], 20 µL TCEP (53 mM in $ddH_2O$) and 2.5 µL TBTA (83.5 mM in DMSO). Samples were gently vortexed and the click reaction was initiated by the addition of 40 µL $CuSO_4$ solution (50 mM in $ddH_2O$). The mixture was incubated at 25 °C for 1.5 h. Afterwards, proteins were precipitated in 18 mL acetone at -20 °C overnight. Protein pellet washing, enrichment and on-beads trypsin digest was done as described above using 100 uL of avidin-beads slurry and digestion buffer without the thiourea. After the overnight incubation with trypsin, the beads were transferred onto the membrane filter (Ultrafree-MC and –LC, Durapore PVDF-0.22 µm, *Merck Millipore*) and spun down at 1,000 g for 1 min. The flow through was processed as described above for trypsin digest. Beads loaded on a membrane filter were washed twice with 50 µL of $ddH_2O$, thrice with 600 µL PBS and thrice with 600 µL $ddH_2O$. Beads were resuspended in 150 µL TEV buffer (141 µL of $ddH_2O$, 7.5 µL 20xTEV buffer supplied by manufacturer, 1.5 µL of 100µM DTT) and transferred from the membrane filter into eppendorf tube. Beads were spun down at 400 g for 2 min and supernatant was removed. Beads were resuspended in 150 µL of TEV buffer and incubated with agitation at 29 °C with 5 µL of AcTEV protease (10 U/µL, *Invitrogen*) overnight. The suspension was transferred onto the membrane filter and spun down at 1,000 g for 1 min, beads were further washed twice with 50 µL of $ddH_2O$. Flow through was collected and after addition of 4 µL of formic acid it was desalted on stage tips (double $C_{18}$ layer, Empore disc-C18, 47MM, *Agilent Technologies*). Stage tips were equilibrated with 70 µL acetonitrile, spun at 1,000 g for 2 min, then samples were loaded and spun at 1,000 g for 5 min. Stage tips were washed thrice with 70 µL 0.5% formic acid and TEV cleaved proteins eluted with two portions 30 µL of 80% acetonitrile and 0.5% formic acid. Samples were combined and dried by lyophilization. Peptides were reconstituted in 30 µL 1% (v/v) formic acid, prepared for mass spectrometry by filtering through a membrane filter (Ultrafree-MC and –LC, Durapore PVDF-0.22 µm, *Merck Millipore*) and transferred into mass vials.

**DSSO crosslinking.** HeLa cells were cultivated in 15 cm dishes and transfected with pCMV-SPORT6 plasmid with FICD wt or FICD E234G with GGGGS sequence connected to 6 x HIS tag at C-terminus. Cells were supplemented with 15 mL fresh DMEM media and transfected using 8 µL Lipofectamine®2000 (*Thermo Fisher Scientific*) and 16 µg of the pCMV-SPORT6 respective plasmid in 2 mL of the Opti-MEM™ (*Gibco™*) media according to manufacturer protocol. Cells were incubated for 36 h then washed twice with 3 mL PBS, scraped into the 1 mL PBS containing 2 mM DSSO crosslinker (20 µL of 100 mM DSSO stock in DMSO) and incubated at 37 °C for 1 h. Crosslinking reaction was stopped by addition of 0.5 mL 50 mM Tris pH 8.0. Cells were harvested at 600 rpm for 5 min and washed twice with 0.5 mL ice cold PBS and lysed in 1 mL ice cold lysis buffer (1% (v/v) NP40, 1% (w/v) sodium deoxycholate and 1 tablet protease inhibitor (cOmplete™, Mini, EDTA-free protease inhibitor cocktail, *Roche*) in 10 mL PBS). Lysis was done for 30 min at 4 °C while rotating. Insoluble fraction was pelletized (10 min, 14,000 g, 4 °C), supernatant was transferred to prewashed (3 x 1 mL of lysis buffer) Ni-NTA agarose (200 µL of beads slurry, *QIAGEN*) and incubated at 4 °C for 2 h. The beads were spin down at 2,000 rpm and supernatant was removed, beads were resuspended in 600 µL mL high salt wash buffer (25 mM Tris pH 8.0, 500 mM NaCl, 20 mM imidazol) and transferred on filter columns (Pierce® centrifuge columns), washed twice with 600 µL mL wash buffer (25 mM Tris pH 8.0, 150 mM NaCl, 80 mM imidazol). Beads were resuspended in 600 µL of PBS and transferred into 1.5 eppendorf tube, spin down at 2,000 rpm and supernatant was removed. Enriched proteins were digested as described above for chemical-proteomics experiments and identified using LC-MS/MS as described below.

**Whole proteome analysis.** SH-SY5Y were washed twice with PBS and homogenized in ice-cold lysis buffer (1% (v/v) NP40, 1% (w/v) sodium deoxycholate and 1 tablet protease inhibitor (cOmplete™, Mini, EDTA-free protease inhibitor cocktail, *Roche*) in 10 mL PBS) using ultrasonication at 40 % intensity for 10 s. Lysis was done for 30 min at 4 °C while rotating. Insoluble fraction was pelletized (10 min, 14,000 g, 4 °C) and protein concentration was determined by bicinchoniic acid assay (BCA, *Carl Roth GmbH + Co.*). For next analysis 100 µg of the total protein was used. Proteins were precipitated by addition of 900 µL acetone and incubation overnight at -20 °C. The protein pellet was harvested by centrifugation at 9,000 g for 15 min at 4 °C and washed once with 0.5 mL of ice-cold methanol. Proteins were reconstituted in 200 µL digestion buffer (20 mM HEPES, pH 7.5, 7 M urea, 2 M thiourea), reduced (0.2 µL 1 M DTT, 45 min, 25 °C) and alkylated (2 µL, 30 min, 25 °C, in the dark). The alkylation reaction was quenched by addition of 0.8 µL 1M DTT and incubation for 30 min at 25 °C. Proteins were pre-digested with 1 µL LysC (*Wako*) at 25 °C for 4 h. 600 µL 50 mM TEAB buffer was added and the proteins were digested overnight with 1.5 µL sequencing grade trypsin (0.5 mg/mL, *Promega*) at 37 °C. The following day the samples were acidified with 10 µL formic acid to a pH of 2 – 3. Peptides were desalted on 50 mg SepPak C18 cartridges (*Waters Corp.*) on a vacuum manifold. The cartridges were equilibrated with 1 mL acetonitrile, 1 mL 80% acetonitrile and 3 mL 0.5% formic acid. The samples were loaded on the cartridges and subsequently washed with 5 mL 0.5% formic acid. The peptides were eluted with two times 250 µL 80% acetonitrile, 0.5% formic acid. Samples were combined and dried by lyophilization. Peptides were reconstituted in 30 µL 1% (v/v) formic acid, prepared for mass spectrometry by filtering through a membrane filter (Ultrafree-MC and –LC, Durapore PVDF-0.22 µm, *Merck Millipore*) and transferred into mass vials.

**Mass Spectrometry.** MS analysis was either performed on an Orbitrap Fusion or an Q Exactive Plus instrument coupled to an Ultimate3000 Nano-HPLC via an electrospray easy source (all *Thermo Fisher Scientific*). Samples were loaded on a 2 cm PepMap RSLC C18 trap column (particles 3 µm, 100A, inner diameter 75 µm, *Thermo Fisher Scientific*) with 0.1% TFA and separated on a 50 cm PepMap RSLC C18 column (particles 2 µm, 100A, inner diameter 75 µm, *Thermo Fisher Scientific*) constantly heated at 50 °C. The gradient was run from 5-32% acetonitrile, 0.1% formic acid during a 152 min method (7 min 5%, 105 min to 22%, 10 min to 32%, 10 min to 90%, 10 min wash at 90%, 10 min equilibration at 5%) at a flow rate of 300 nL/min. For measurements of chemical-proteomic samples on the fusion instrument survey scans (m/z 300-1,500) were acquired in the orbitrap with a resolution of 120,000 at m/z 200 and the maximum injection time set to 50 ms (target value 2e5). Most intense ions of charge states 2-7 were selected for fragmentation with high-energy collisional dissociation at a collision energy of 30%. The instrument was operated in top speed mode and spectra acquired in the ion trap with the maximum injection time set to 50 ms (target value 1e4). The option to inject ions for all available parallelizable time was enabled. Dynamic exclusion of sequenced peptides was set to 60 s. Real-time mass calibration was based on internally generated fluoranthene ions. Data were acquired using Xcalibur software version 3.0sp2 (*Thermo Fisher Scientific*).

Binding site identifications of AMPylated peptides were performed on the fusion instrument with slightly changed parameters. The scan range for survey scans in the orbitrap was changed to 300-1,700 with an AGC target value of 4e5. HCD fragmentation was performed at a collision energy of 27% and spectra first acquired in the ion trap with a maximum injection time of 30 ms and an AGC target value of 2e4. Peptides with targeted masses of m/z 501.28, 251.14 and 167.76 (adenine_TEV singly, doubly and triply charged), as well as m/z 615.31, 308.16 and 205.77 (adenosine_TEV singly, doubly and triply charged) were further selected for ETD fragmentation while highest charge states were prioritized. ETD fragmentation scans were acquired in the orbitrap with a resolution of 30,000, a maximum injection time of 1,000 and an AGC target value of 2e5.

For measurements of chemical-proteomic samples on the Q Exactive Plus instrument survey scans (m/z 300-1,500) were acquired in the orbitrap with a resolution of 70,000 at m/z 200 and the maximum injection time set to 80 ms (target value 3e6). Data dependent HCD fragmentation scans of the 12 most intense ions of the survey scans were acquired in the orbitrap at a resolution of 17,500, maximum injection time of 50 ms as well as minimum and maximum AGC targets of 5e3 and 5e4, respectively. The isolation window was set to 1.6 m/z. Unassigned and singly charged ions were excluded for measurement and the dynamic exclusion of peptides enabled for 60 s. The lock-mass ion 445.12002 from ambient air was used for real-time mass calibration on the Q Exactive Plus. Data were acquired using Xcalibur software version 3.1sp3 (*Thermo Fisher Scientific*).

**Quick change point mutation**. Quick change point mutation of FICD H363A was performed using following primers 5´-CCGTTGCCATCAATGAAAGGGGCGATGTAAACGAGTTTATAATG-3´ and 5´-CATTATAAACTCGTTTACATCGCCCCTTTCATTGATGGCAACGG- 3´ and pCMV-SPORT6 with wt FICD as a template. PCR reaction mixture contained 5xPhusion buffer 10 µL, 10 mM dNTPs 1 µL, 10 µM each primer 0.75 µL, template 50 ng 1 µL, ddH$_2$O 32 µL, DMSO 4 µL and Phusion polymerase (*NEB*) 0.5 µL. PCR cycle (initial denaturation 98 °C, 3 min then 25 cycles of 95 °C 45 s, 65 °C 30 s, 72 °C 3 min; final elongation at 72 °C for 5 min). Template was digested using DpnI (8 µL of PCR mixture, 10xCutSmart buffer (*NEB*) 1 µL and DpnI (*NEB*) 1 µL) at 37 °C for 4 h. Resulting plasmid was amplified in *E. coli* XL1 blue. Identity of the isolated plasmid was verified by sequencing.

**Cloning of FICD with C-terminal 1xFLAG tag**. To construct the pCMV-SPORT6 plasmid containing FICD wt gene with C-terminal GSDYKDDDDK tag. The FICD gene was extended in PCR using following primers 5´-

GGCAGCGACTACAAAGACGATGACGACAAGTAACCCTAGAAATCCTCAG - 3´ and 5´-CTTTGTAGTCGCTGCCGGGCTTCACAGGAAGC - 3´. PCR reaction mixture contained 5xPhusion GC reaction buffer 10 µL, 10 mM dNTPs 1 µL, 10 µM each primer 0.75 µL, template 150 ng 0.3 µL, DMSO 1.5 µL, dd$H_2O$ 35.2 µL and Phusion polymerase (*NEB*) 0.5 µL. PCR cycle (initial denaturation 98 °C, 3 min then 25 cycles of 98 °C 45 s, 72 °C 3.5 min; final elongation at 72 °C for 7 min). Next, 8 µL of PCR product was digested supplemented with 1 µL of CutSmart buffer (NEB) and 1 µL DpnI (*NEB*) and incubated at 37 °C for 3 h. Resulting mixture was transformed into *E. coli* Xl1 blue. Identity of the isolated plasmid was verified by sequencing.

**ADP-ribosylation.** The reference list of ADP-ribosylated proteins was taken from ref. 27. In total, it includes 484 proteins identified as ADP-ribosylated by PARP-1, PARP-2 or PARP-3 with known site of modification (**Table S3**). All presented heatmaps and summaries of AMPylated proteins are without the AMPylation-ADP-ribosylation overlapping proteins.

**Immunoprecipitation-click-chemistry.** HeLa cells were cultivated in 15 cm dishes and treated with 100µM pro-N6pA (13 µL 100mM in DMSO) for 16 h as described above. Cells were washed twice with 3 mL ice cold PBS and lysed in 1mL ice cold IP lysis buffer (50mM Tris HCl, pH 8; 250mM NaCl; 1% NP-40, 5mM EDTA, with protease inhibitors) while mechanically detached by scraping. Lysis was done for 30 min at 4 °C while rotating. Insoluble fraction was pelletized (10 min, 14,000 g, 4 °C). Protein lysates (3.5 - 4 mg) from pro-N6pA or DMSO treated cells were incubated with 10 µL of primary antibodies (anti-PFKP (rabbit) or anti-PPME1 (rabbit) both 1 mg/mL, *Bethyl laboratories Inc.*) at 4 °C with gentle agitation overnight. Beads were spun at 400 g, 4 °C for 2 min. Approximately 70 µL Pierce Protein A/G agarose beads slurry (*Thermo Scientific*) was washed twice with 0.5 mL ice-cold IP lysis buffer, added to the lysates and incubated at 4 °C for 4 hours with gentle agitation. The beads were washed thrice with 0.5 mL of ice cold IP lysis buffer, then twice with ice cold IP wash buffer (50 mM Tris/HCl pH 8.5, 75 mM KCl) and finally resuspended in 97 µL of PBS supplied with 1 µL 5-rhodamine-azide (10 mM in DMSO, *Carl Roth GmbH + Co. KG*), 1 µL TCEP (53 mM in dd$H_2O$) and 0.2 µL TBTA (83.5 mM in DMSO). The click-reaction mixture was incubated at 25 °C for 1.5 h with agitation, supernatant was removed and 20 µL of gel loading buffer was added to the beads slurry. Immunoprecipitated proteins were released from the beads into the gel loading buffer at 95 °C for 5 min and separated on 12.5% SDS PAGE. In-gel fluorescence was scanned as described above.

**Immunoblot analysis.** SDS-PAGE separated proteins were blotted on a methanol-activated PVDF membrane for 1 h at 22 V (blotting buffer: 48 mM Tris, 39 mM glycine, 0.0375% SDS, 20% methanol) using a Bio-Rad Trans-Blot SD semi-dry transfer cell setup. The membrane was then blocked in blocking buffer (5% milk powder in PBS with 0.5% Tween-20), washed and incubated with primary antibody (Table S12) (diluted 1:1,000 in blocking buffer) for 16 h at 4 °C. The membrane was washed thrice 10 min in 0.5% Tween-20 in PBS, and then incubated with secondary antibody (goat anti-rabbit IgG-HRP (Santa Cruz), diluted 1:10,000 in blocking buffer) at 25 °C for 1 h. The membrane was washed again with 0.5% Tween-20 in PBS and subsequently incubated in ECL (Bio-Rad) and imaged using a Fujifilm LAS-4000 equipped with a Fujifilm Fujinon VR43LMD3 lens and chemoluminescence detection mode.

**Table S12 | Antibodies used in immunohistochemistry and immunoprecipitations**

| Antigen | Dilution | Vendor | Catalog # |
|---|---|---|---|
| AMPylated Tyrosine | 1:1000 | Merck | ABS184 |
| ARL13B | 1:200 | Proteintech | 17711-1-AP |
| CTIP2 | 1:500 | Abcam | ab18465 |
| Doublecortin (DCX) | 1:2,000 | Merck Millipore | ab2253 |
| F-Actin (Phalloidin) | 1:40 | Thermo Fisher | A12381 |
| FICD | 1:100 | Sigma | HPA021390 |
| FLAG | 1:200 | Sigma | F1804 |
| GFP | 1:1,000 | Aves Lab | GFP-1020 |
| GM130 | 1:500 | GeneTex | GTX130351 |
| HSPA5 | 1:1,000 | Abcam | ab21685 |
| KI67 | 1:100 | Dako | M7248 |
| MAP2 | 1:500 | Sigma Aldrich | M4403 |
| NESTIN | 1:200 | Millipore | MAB5326 |
| NEUN | 1:500 | Merck Millipore | MAB377 |
| PAX6 | 1:500 | Biolegend | PRB-278p |
| PDI | 1:500 | Abcam | ab2792 |

| | | | |
|---|---|---|---|
| PFKP | 1:100 | Bethyl Lab. Inc. | A304-283A |
| PH3 | 1:500 | Millipore | 06-570 |
| Phospho-TAU (Ser202, Thr205) | 1:500 | Thermo Fisher | MN1020 |
| PPME1 | 1:100 | Bethyl Lab. Inc. | A304-762A |
| TBR1 | 1:500 | Abcam | ab31940 |
| β-Catenin | 1:500 | BD Biosciences | 610154 |

**Transfection of SH-SY5Y cells by nucleofection.** SH-SY5Y cells were grown to 80 % confluency, washed with PBS, dissociated to single cells with 0.05 % trypsin-EDTA and counted. Per construct, 2 mio. cells were resuspended in 100 µl nucleofection buffer (50 mM Hepes, 90 mM $Na_3PO_4$, 5 mM KCl, 0.15 mM $CaCl_2$) and a total of 4 µg DNA of a 2:1 ratio of pCMV-SPORT6 plasmid containing wt FICD or FICD E234G/H363A mutant (gift from A. Itzen, TUM) and pCAG-IRES-GFP or of only pCAG-IRES-GFP (control) was added. The suspension was filled into an aluminium electrode cuvette and exposed to program G00.04 in an Amaxa II Nucleofector (*Lonza*). Sequentially, for the quantification of probe signal upon FICD OX, 30,000 cells/well were plated on coverslips in a 24-well-plate in triplicates and for proteome analyses, 10 Mio. cells were plated per 15 cm dish. Transfected cells were cultured for 48 h, adding pro-N6pA for the last 16 h of culture, before analysis.

**Probe staining of fixed cells.** Cells and organoids were cultured with medium containing 100 µM pro-N6pA or 1:1,000 dilution of DMSO for 16 h prior to fixation. PFA-fixed cells on coverslips and thawn, rehydrated organoid sections were permeabilized using 0.1 % Triton X100 in PBS for 5 min. Then they were incubated with click-chemistry staining mix (10 µM rhodamine-azide, 1 mM $CuSO_4$, 10 mM freshly prepared sodium ascorbate in PBS) at rt for 2 h in the dark, followed by several washes with PBS[47]. For nuclear counterstaining only, cells or sections were incubated with 0.1 µg/mL DAPI in PBS for 15 min. For co-staining by immunohistochemistry, 10 min postfixation with 4 % PFA in PBS was performed after the click-reaction with rhodamine azide, followed by the immunohistochemistry protocol (see above).

**Endoglycosidase H assay.** HeLa cells were transiently transfected with the pCMV-SPORT6-wtFICD plasmid as described above. Cells lysate (100 µg total protein) was digested by the Endo H (#P070S, New England Biolabs) as described by the manufacturer, but extending the digestion time to 1.5 h in total reaction volume of 50 µL.

**qPCR of FICD mRNA levels.** Cells were washed and scraped with cold PBS and the pellet was lysed in QIAzol® Lysis Reagent (#79306, Qiagen). RNA was extracted using RNA Clean & Concentrator Kit (#R1015, Zymo Research) and cDNA was synthesised from 1 µg RNA each using SuperScript III reverse transcriptase (#18080-044, Thermo Fisher) with Oligo(dT)$_{12\text{-}18}$ primers (#18418012, Thermo Fisher) according to the manufacturer's protocol. Subsequently, qPCR was performed in triplicates on a LightCycler® 480 II (Roche) using the LightCycler® 480 SYBR Green I Master (#04707516001, Roche) with the following reaction mix: 1 µL of each primer (5 µM), SYBR Green I Master 5 µL, $H_2O$ 2 µL and 1 µL of cDNA. The primer sequences were the following for FICD 5'-GCGGTTGGGGTTCTGAC- 3' and 5'-TGAAGCCATTGGTATGAGCA- 3' and GAPDH 5'-AATCCCATCACCATCTTCCAGGA- 3' and 5'-TGGACTCCACGACGTACTCAG- 3'. PCR cycle (initial denaturation 95 °C, 10 min then 45 cycles of 95 °C 10 s, 60 °C 10 s and 72 °C 10 s) with melting curve 95 °C to 65 °C with 15s per step. $C_p$ values were determined from the second derivative maximum of the amplification curve and fold expression was calculated using the $\Delta\Delta C_p$ method.

**Quantification of AMPylation upon FICD overexpression.** SH-SY5Y cells were transfected by nucleofection with GFP, wt FICD + GFP (2:1) or FICD E234G + GFP (2:1) as described above. Cells were cultured for 48 h after transfection, treating with pro-N6pA for the last 16 h. Then they were fixed with 4 % PFA, followed by click-reaction and counterstaining of GFP and of nuclei with DAPI as described above. Confocal images were taken at 3 positions per coverslip. For quantification of AMPylation, using Fiji, the outline of individual GFP$^+$ cells were drawn and then the rhodamine signal was measured and normalised to cell size (area). Results are shown for $n$ = 90 cells per condition, normalised to GFP$^+$ control.

**Reprogramming of fibroblasts to induced pluripotent stem cells (iPSCs).** Human iPSCs were prepared by reprogramming newborn foreskin fibroblasts (CRL-2522, ATCC). As feeders 2.5x10$^5$ NuFF3-RQ IRR human newborn foreskin fibroblasts (GSC-3404, GlobalStem) were seeded per well of a 6-well tissue culture dish with advanced MEM (12491015, Thermo Fisher Scientific) supplemented with 5% HyClone Fetal Bovine Serum (SV30160.03HI, GE Healthcare), 1% MEM NEAA and GlutaMAX (11140050; 35050061 Thermo Fisher Scientific). On day 1, a newborn foreskin fibroblast culture of 70-80% confluency was dissociated using 0.25% Trypsin-EDTA (25200056, Life Technologies), counted and seeded on the pre-plated NuFF3-RQ cells at two densities: $2x10^4$ and $4x10^4$ cells/well. On day 2, the medium was changed to Pluriton Reprogramming Medium (00-0070, Stemgent) supplemented with 500 ng/ml carrier-free B18R Recombinant Protein (03-0017, Stemgent). On days 3-18, a cocktail of modified mRNAs (mmRNAs) was transfected daily. For that, *OCT4*, *SOX2*, *LIN28A*, *CMYC*, and *KLF4* mmRNAs at a 3:1:1:1:1 stoichiometric ratio in Opti-MEM I Reduced Serum Medium (13778150, Thermo Fisher Scientific) in a total volume of 105µl was combined with a mix of 92µl Opti-MEM I Reduced Serum Medium and 13µl Lipofectamine RNAiMAX Transfection Reagent (31985062, Thermo Fisher Scientific) after an incubation at RT for 15 min. Cells were transfected for 4hrs, washed and fresh reprogramming medium supplemented with B18R was added to the cultures. The mmRNAs were provided by the RNA CORE unit of the Houston Methodist Hospital and contained the following modifications: 5-Methyl CTP, a 150 nt poly-A tail, ARCA cap and Pseudo-UTP. Already 5 days after the first transfection, initial morphological changes became apparent, and first induced pluripotent stem cell (iPSC) colonies appeared by day 12-15. On day 16, the medium was changed to STEMPRO hESC SFM (A1000701, Thermo Fisher Scientific) for five days. Afterwards, the iPSC colonies were harvested after 40min incubation at 37°C with 2mg/ml Collagenase Type IV (17104019, Thermo Fisher Scientific) solution in DMEM/F12 (31331093, Thermo Fisher Scientific). The iPSCs were plated on γ-irradiated mouse embryonic fibroblasts (MEFs) and grown in STEMPRO hESC SFM for 10 additional passages. Then the iPSCs were adapted to a feeder-free culture system, using mTeSR1 (05850, StemCell Technologies) and plates coated with LDEV-Free Geltrex (A1413302, Thermo Fisher Scientific).

**pro-N6pA / 5-Ethynyl-uridine and RNAse treatment.** In order to exclude incorporation of pro-N6pA into RNAs[55], HeLa cells were grown on coverslips. For the last 16 h of culture proior to PFA fixation, they were incubated with the pro-N6pA probe (100 µM = 1:1000 from stock), DMSO (1:1000) or 5-ethynyl-uridine (5-EU; 1 mM = 1:100 from stock in $H_20$) in medium. The fixed cells were treated with different amounts of RNAse $I_f$ prior to probe staining by click-chemistry: Cells were permeabilized applying 0.3 % Triton X100 in PBS for 5 min and the incubated with 5 to 100 U/coverslip RNAse $I_f$ (M0243S, New England Biolabs; unspecific RNAse cleaving both purine and pyrimidine residues) in 1 x NEB3 buffer for 30 min or 2h at 37 °C. NEB3 without RNAse served as control. After the RNAse treatment the fixed cells were washed thrice with PBS.

**Rating of MAP2+ neuronal processes intruding the progenitor zone in COs**: After IHC for GFP (to identify electroporated germinal zones) and MAP2 (for neuronal dendrites), combined with DAPI for nuclear counterstain, confocal laser scanning images were acquired. A rating system was applied to categorize ventricles according to invading MAP2 processes within the progenitor zone, recognized in the DAPI channel as cell-dense zone lining the ventricle-like lumen, with the grading: 0 = no invading processes, 1 = few invading processes, 2 = a lot of invading processes. Every one ventricle was taken as n.

**GO Term and Network Analysis using STRING.db.** STRING database was used to perform Network and GO Term Analysis of AMPylated proteins identified in the different cell lines[50]. Proteins overlapping with ADP-ribosylation were excluded. Network nodes represent the AMPylated proteins. Network edges represent protein-protein association. Thickness of the network edges was chosen to indicate edge confidence, meaning strength of the data support for the shown interaction. Minimum required interaction score was set to 0.400. The Table S13 contains all GO Term analyses for the cell types. Functional enrichments in each network were considered significant in case of a false discovery rate *FDR* < 0.05.

### (D) NMR spectra

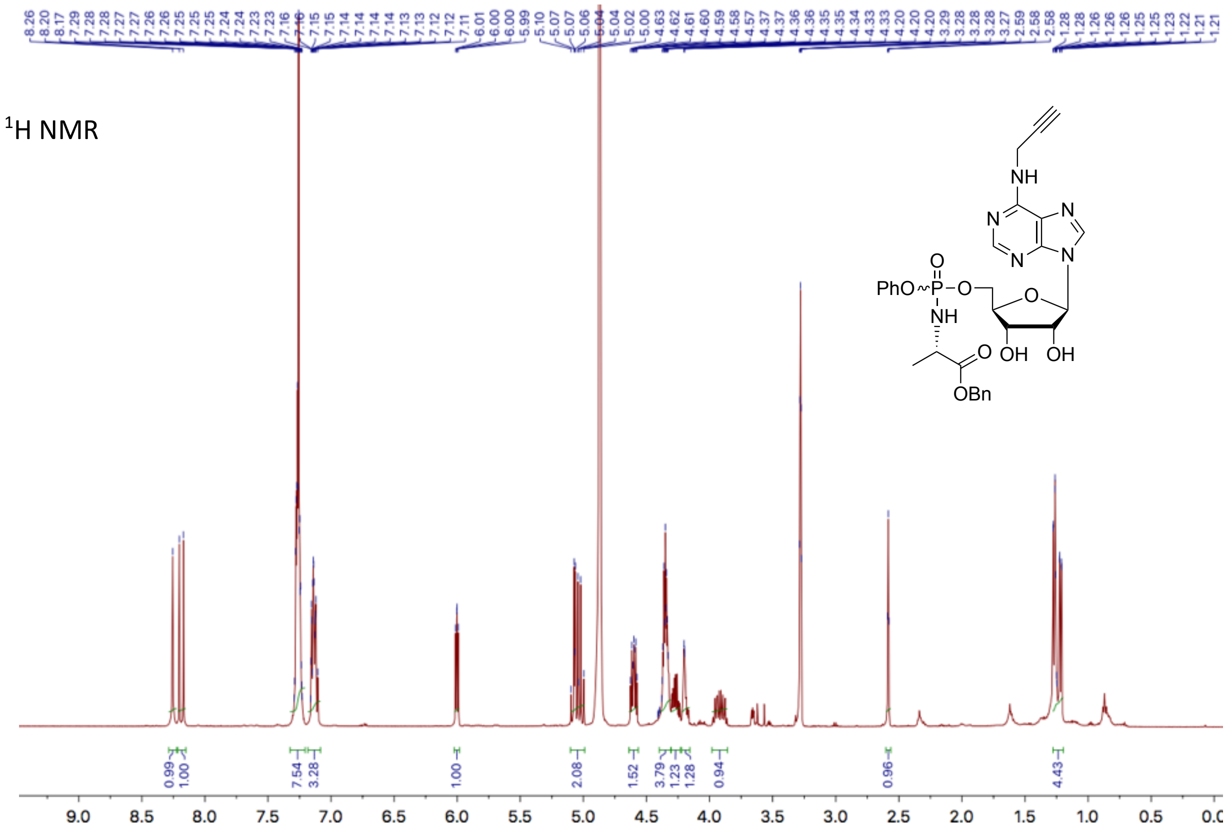


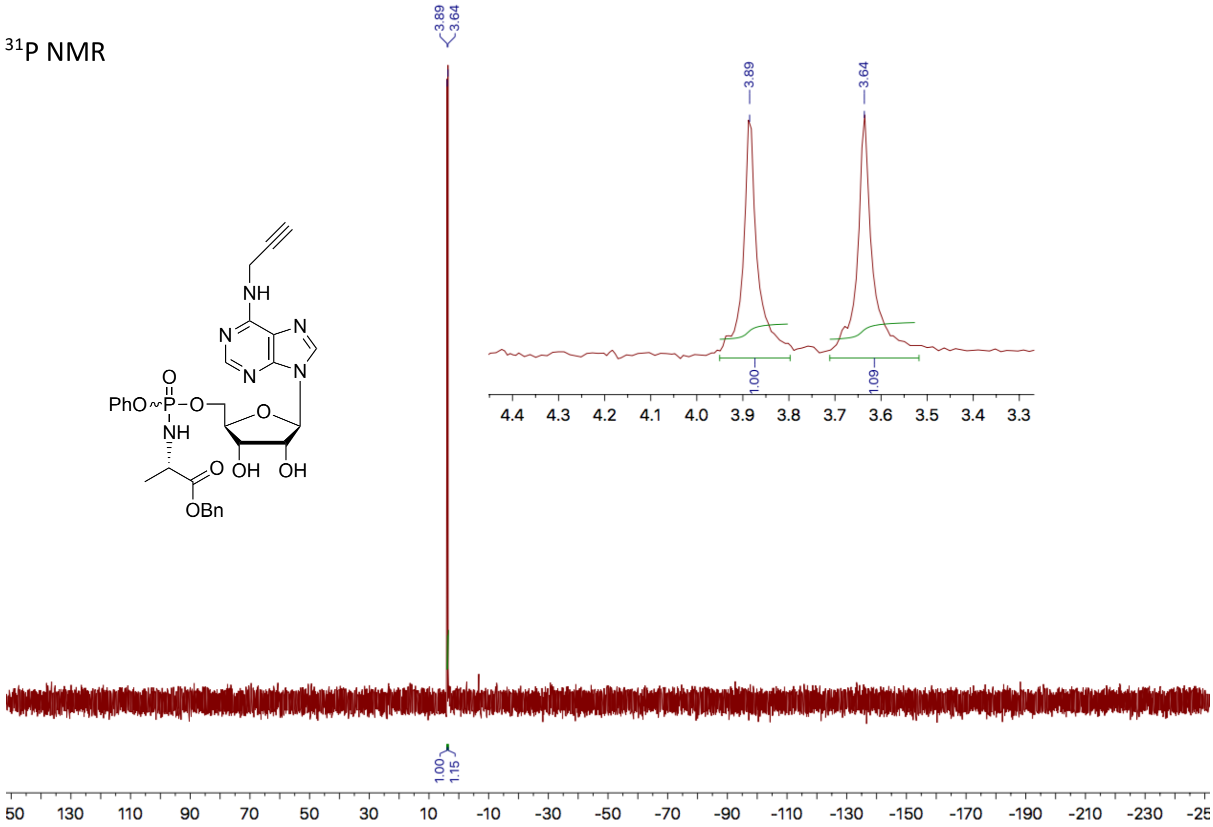


### (E) References
